# Near telomere-to-telomere *Linum* genomes reveal a lineage-specific DNA transposon associated with chromosome architecture remodeling

**DOI:** 10.64898/2026.05.01.722228

**Authors:** Frank M. You, Chunfang Zheng, Pingchuan Li, Tara Edwards, Sean Walkowiak, Liqiang He, Jin Xiao, Xiue Wang, Sylvie Cloutier

**Affiliations:** Ottawa Research and Development Centre, Agriculture and Agri-Food Canada, Ottawa, 960 Carling Avenue, Ottawa, ON, K1A 0C6, Canada; Grain Research Laboratory, Canadian Grain Commission, 196 Innovation Drive, Winnipeg, MB, R3T 2N2, Canada; School of Tropical Agriculture and Forestry, School of Tropical Crops, Hainan University, Haikou, China; Key Laboratory of Crop Genetics & Germplasm Enhancement and Utilization, College of Agriculture/Zhongshan Biological Breeding Laboratory/CIC-MCP, Nanjing Agricultural University/JCIC-MCP, Nanjing 210095, China

**Author notes:** Correspondence: Frank M. You; Sylvie Cloutier.

**Keywords:** *Linum*, flax, chromosome evolution, telomere-to-telomere genome assembly, pericentromeric regions, transposable elements, DNA transposons, karyotype evolution, post-polyploid diploidization, structural variation

## Abstract

Chromosome number variation and structural reorganization are key drivers of plant evolution, yet their genomic basis remains unclear due to incomplete representation of repetitive regions in existing assemblies. The *Linum* genus exhibits exceptional karyotypic diversity (*n* = 7–43), providing a powerful system to investigate chromosome evolution. Here, we generated near telomere-to-telomere (T2T) genome assemblies for four species, including cultivated flax (*L. usitatissimum* cv. CDC Bethune; *n* = 15), its wild progenitor (*L. bienne*; *n* = 15), and two related species (*L. decumbens* and *L. grandiflorum*; *n* = 8). Together with published genomes of *L. lewisii* (*n* = 9) and *L. tenue* (*n* = 10), these enabled reconstruction of chromosome evolution across six lineages. Phylogenomic analyses revealed a shared ancestral whole-genome duplication (WGD) associated with the *n* = 9 karyotype, followed by lineage-specific WGDs and divergent diploidization. The transition from *n* = 8 to the derived *n* = 15 flax lineage not only occurred without chromosome length expansion, but also with genome size reduction, indicating extensive internal restructuring. Comparative analyses showed that this restructuring was associated with lineage-specific expansion of a single DNA transposon family (*TE_00003234*; *hAT*), which is highly enriched in expansive pericentromeric regions that are characterized by low gene density and nucleotide diversity, suppressed recombination, segregation distortion, and extensive synteny disruption, unlike the LTR retrotransposon-rich pericentromeres typical of most plant genomes. These findings support a model in which lineage-specific DNA transposon expansion is associated with remodeling of pericentromeric architecture and large-scale chromosome restructuring following polyploidization.

## INTRODUCTION

*Linum* is an important land plant genus that includes cultivated flax (syn. flaxseed, linseed), *L. usitatissimum*, which is a food crop high in α-linolenic acid (ALA, omega-3 fatty acid), lignans, and fiber (Goyal et al. 2014), whose oil also has industrial applications, such as in paints and linoleum (Jhala and Hall 2010). In addition, tall flax morphotypes have been bred for their long stem fibers used in the fabrication of linen and other fiber products. Flax cultivation dates back more than 6,000 years; however, other species within the genus originated millions of years ago (MYA) and are important for understanding crop domestication, as well as evolution within the genus (Jhala and Hall 2010). The breadth of variation in the number of chromosomes within *Linum* sp., suggests its rich and complex evolutionary history (Goldblatt 2007; Rice et al. 2014).

Chromosome number variation is a fundamental feature of plant genome evolution and a major driver of diversification, speciation, and adaptation. Changes in chromosome number can arise through processes such as whole-genome duplication (WGD), aneuploidy, chromosome fusion, and chromosome fission, often accompanied by extensive genome restructuring (Stebbins 1950; Schubert and Lysak 2011). However, understanding the genomic mechanisms underlying chromosome evolution has remained challenging because centromeric and pericentromeric regions—where structural rearrangements and recombination suppression are often concentrated—are highly repetitive and typically fragmented or collapsed in many genome assemblies. Consequently, the architecture of chromosome centers and their roles in genome evolution remain poorly understood. Recent advances in long-read sequencing and assembly algorithms now enable telomere-to-telomere (T2T) genome assemblies, providing unprecedented opportunities to resolve complex genomic regions and investigate chromosome evolution at the nucleotide resolution (Miga et al. 2020; Nurk et al. 2022).

The *Linum* genus belongs to the Linaceae family and represents an excellent system for studying chromosome evolution due to its remarkable karyotypic diversity. The genus comprises over 180 diploid species with haploid chromosome numbers ranging from *n* = 7 to *n* = 43 (Goldblatt 2007; Rice et al. 2014). Phylogenetic analyses divide the *Linum* genus into two major evolutionary lineages: a blue-flowered clade, including the *Linum* and *Dasylinum* sections, and a yellow-flowered clade comprising the *Linopsis*, *Syllinum*, *Cathartolinum*, and their related groups (McDill et al. 2009; Bolsheva et al. 2017; Schneider et al. 2016; Villalvazo-Hernandez et al. 2022) (**Supplemental Figure S1**). The terms “blue-flowered” and “yellow-flowered” are historical terms for these two lineages and reflect predominant (and likely ancestral) flower-color states rather than an exclusive diagnostic phenotype, considering that multiple derived color transitions (e.g., red, pink, white) occur within both clades (Diederichsen 2007; Villalvazo-Hernandez et al. 2022).

Overlaying chromosome numbers onto this phylogeny suggests that the two lineages share a common ancestral karyotype with a base chromosome number of 9, supported by the prevalence of species with 9 chromosomes that are distributed across basal branches of both clades (Valdes-Florido et al. 2023; Chennaveeraiah and Joshi 1983) (**Supplemental Figure S1**). Subsequent diversification involved multiple lineage-specific departures from this ancestral state, including both chromosome number reductions and increases. For example, cytogenetic and phylogenetic evidence suggests that several blue-flowered lineage species with 8 chromosomes, such as *L. decumbens* and *L. grandiflorum*, are derived from an ancestral karyotype with 9 chromosomes, whereas the karyotypes with 15 chromosomes of cultivated flax (*L. usitatissimum*) and its wild progenitor (*L. bienne*) likely arose through chromosomal restructuring following polyploidization and subsequent diploidization (Chennaveeraiah and Joshi 1983; Fu et al. 2016; Sugiura 1940). Consistent with this evolutionary scenario, genomic analyses have identified at least two WGD events in flax, including an ancient duplication dating to ∼20–44 million years ago (MYA) and a more recent event somewhere between 3.7–9 MYA (Wang et al. 2012; Sveinsson et al. 2014; You et al. 2018a; You et al. 2018b). Although these duplication events suggest that polyploidization played an important role in *Linum* evolution, the structural mechanisms by which ancestral chromosome numbers diversified into the present-day karyotypes remain ill-defined.

Chromosome-scale genome assemblies have recently been generated for several *Linum usitatissimum* (cultivated flax) accessions, including CDC Bethune (You et al. 2018b), Gaosi (Lu et al. 2025), T397 (Yadav et al. 2025), K-3018 (Arkhipov et al. 2024), YY5 (Sa et al. 2021), K-1531 (Dvorianinova et al. 2023), and Neiya No. 9 (Zhao et al. 2023), as well as from wild species such as *L. lewisii* (*n* = 9) (Innes et al. 2023) and *L. tenue* (*n* = 10) (Gutierrez-Valencia et al. 2022). Despite these advances, many assemblies remain unresolved in highly repetitive regions, limiting our ability to clearly define centromeric and pericentromeric regions, and to characterize large-scale structural rearrangements that are key to chromosome evolution.

In particular, the genomic processes responsible for the transition from ancestral chromosome numbers *n* = 8–9 to the derived 15-chromosome karyotype of *L. bienne* and *L. usitatissimum* remain unresolved. The contribution of transposable elements (TEs) to chromosome reorganization, especially within centromeric and pericentromeric regions, has also not been systematically explored in *Linum*. In many plant genomes, pericentromeric regions are largely occupied by long terminal repeat (LTR) retrotransposons and are characterized by suppressed recombination, low gene density, and often reduced nucleotide diversity (Tian et al. 2009; Yin et al. 2015; Morata et al. 2018; Kent et al. 2017; Zhao and Ma 2013). Whether similar or alternative mechanisms underlie chromosome evolution in *Linum* remains unknown.

Here, we generated high-quality near-T2T genome assemblies for four strategically selected *Linum* accessions representing key evolutionary nodes: cultivated flax (*L. usitatissimum* cv. CDC Bethune), its wild progenitor (*L. bienne* accession LIN1917), and two closely related species with eight chromosomes (*L. decumbens* accession LIN1754 and *L. grandiflorum* accession LIN1530) (**Figure 1A-D**). We also incorporated genome assemblies of two previously sequenced species, *L. tenue* (*n* = 10) (Gutierrez-Valencia et al. 2022), representing the yellow-flowered lineage, and *L. lewisii* (*n* = 9) (Innes et al. 2023), which possesses the ancestral chromosome number of the blue-flowered lineage. By integrating cytogenetic analyses, optical mapping, genetic maps, and comparative genomics with previously published chromosome-level assemblies, we inferred chromosome evolution across six fully sequenced *Linum* species. Our analyses revealed a striking lineage-specific reorganization of chromosome architecture driven by transposable elements, including the emergence of a dominant DNA transposon family that is associated with large-scale reorganization of chromosome architecture during the evolution of *L. bienne* and *L. usitatissimum*.

**Figure 1.**
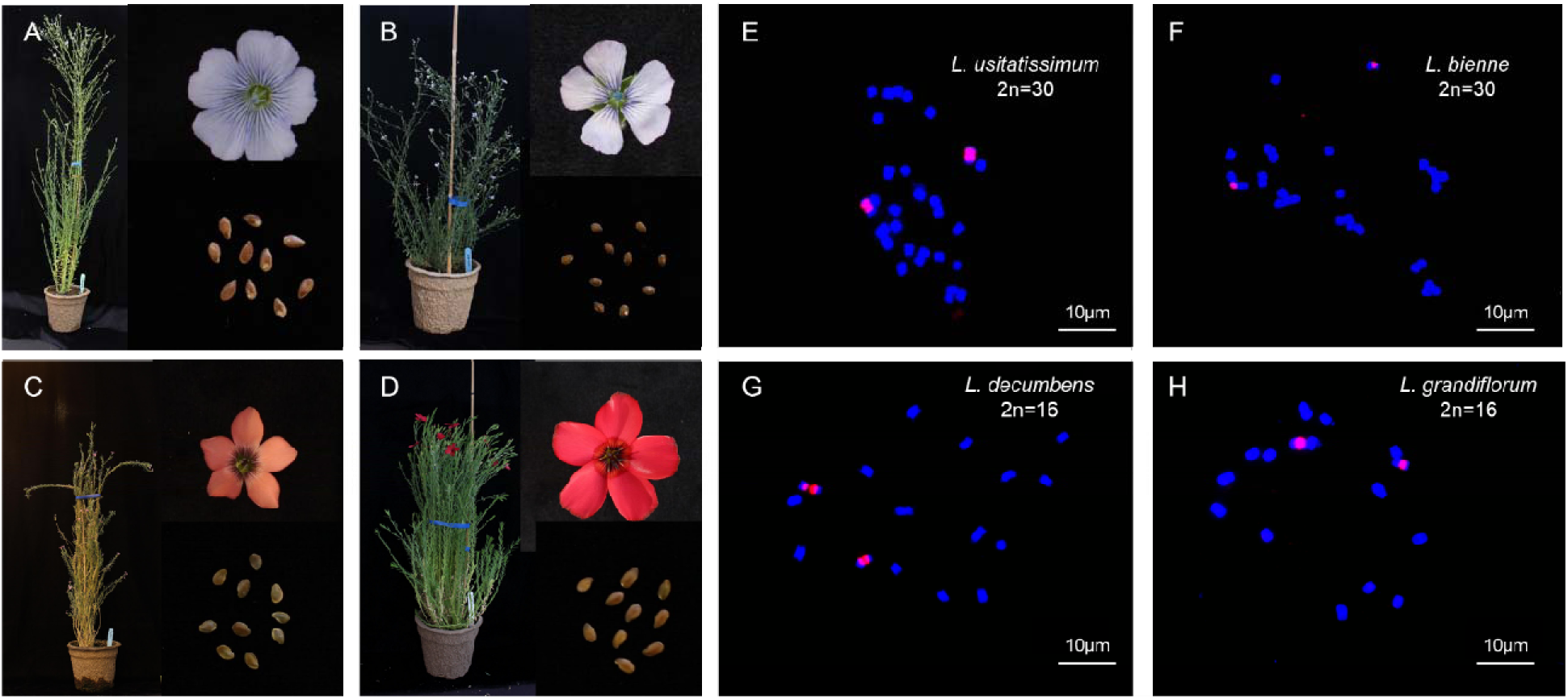
Morphological characteristics and chromosome cytogenetics of four *Linum* accessions. (**A–D**) Whole plants, flowers, and seeds of (**A**) *L. usitatissimum* cv. CDC Bethune, (**B**) *L. bienne* accession LIN1917, (**C**) *L. decumbens* accession LIN1754, and (**D**) *L. grandiflorum* accession LIN1530. (**E–H**) Fluorescence *in situ* hybridization (FISH) analysis of mitotic metaphase chromosomes from the corresponding accessions. The 45S rDNA probe is shown in pink, and chromosomes are counterstained with DAPI (blue).

## RESULTS

### Chromosome number increased without expansion of total chromosome length

Cytogenetic analysis revealed differences in chromosome number among the four *Linum* species studied. Cultivated flax (*L. usitatissimum* cv. CDC Bethune) and its wild progenitor (*L. bienne* accession LIN1917) possess 30 chromosomes, whereas *L. decumbens* (LIN1754) and *L. grandiflorum* (LIN1530) have 16 chromosomes at metaphase of somatic cells (**Figure 1E-H**). Despite this nearly two-fold increase in chromosome number, the total length of the chromosome complements was remarkably conserved across all four species, ranging from 44.27 μm in *L. decumbens* to 47.43 μm in *L. usitatissimum* (**Table 1**). In contrast, the average chromosome length differed substantially, with *n*=8 chromosome species having chromosomes that were approximately twice as long as *n*=15 chromosome species.

**Table 1.** Genome and chromosome sizes of four *Linum* species.

| Species | Accession | No. of chromosomes (*n*) | Total length of chromosomes (μm) | Average chromosome length (μm) | Genome size estimate (Mb) |  |  |
| --- | --- | --- | --- | --- | --- | --- | --- |
|  |  |  |  |  | BioNano assembly | K-mer | Flow cytometry |
| *L. usitatissimum* | CDC Bethune | 15 | 47.43 | 1.58 | 317 | 529.0 | 556.0 |
| *L. bienne* | LIN1917 | 15 | 46.02 | 1.53 | 356 | 527.8 | 581.3 |
| *L. decumbens* | LIN1754 | 8 | 44.27 | 2.77 | 875 | 680.3 | 752.3 |
|  |  |  |  |  | BioNano assembly | K-mer | Flow cytometry |
| *L. grandiflorum* | LIN1530 | 8 | 46.82 | 2.93 | 883 | 767.3 | 849.6 |

Independent genome size estimates based on BioNano optical mapping, k-mer analysis, and flow cytometry consistently showed that *L. decumbens* and *L. grandiflorum* had larger genomes than *L. usitatissimum* and *L. bienne* (**Table 1**). For example, the flow cytometry estimates for *L. decumbens* and *L. grandiflorum* were in the 750–850 Mb range, whereas those of *L. usitatissimum* and *L. bienne* ranged from 550–580 Mb, indicating that genome size reduction in the 15-chromosome lineage is uncoupled from both chromosome number and physical chromosome length, and pointing to internal genomic reorganization rather than whole-chromosome duplication.

### High-quality near telomere-to-telomere assemblies establish a comparative framework for *Linum* genomes

To investigate the genomic basis of chromosome evolution, we generated high-quality near-T2T genome assemblies for four *Linum* species using PacBio HiFi long-read sequencing combined with optical mapping and genetic maps (**Table 2**; **Supplemental Tables S1-S4**). Deep HiFi sequence coverage for all species (38×–148×) with average read lengths of approximately 10–12 kb was produced (**Supplemental Table S1**; **Supplemental Figure S2**). Assemblies generated using hifiasm were highly contiguous, with scaffold N50 values ranging from 30.9 to 60.0 Mb and total assembly sizes closely matching independent genome size estimates (**Tables 1 and 2**).

**Table 2.**
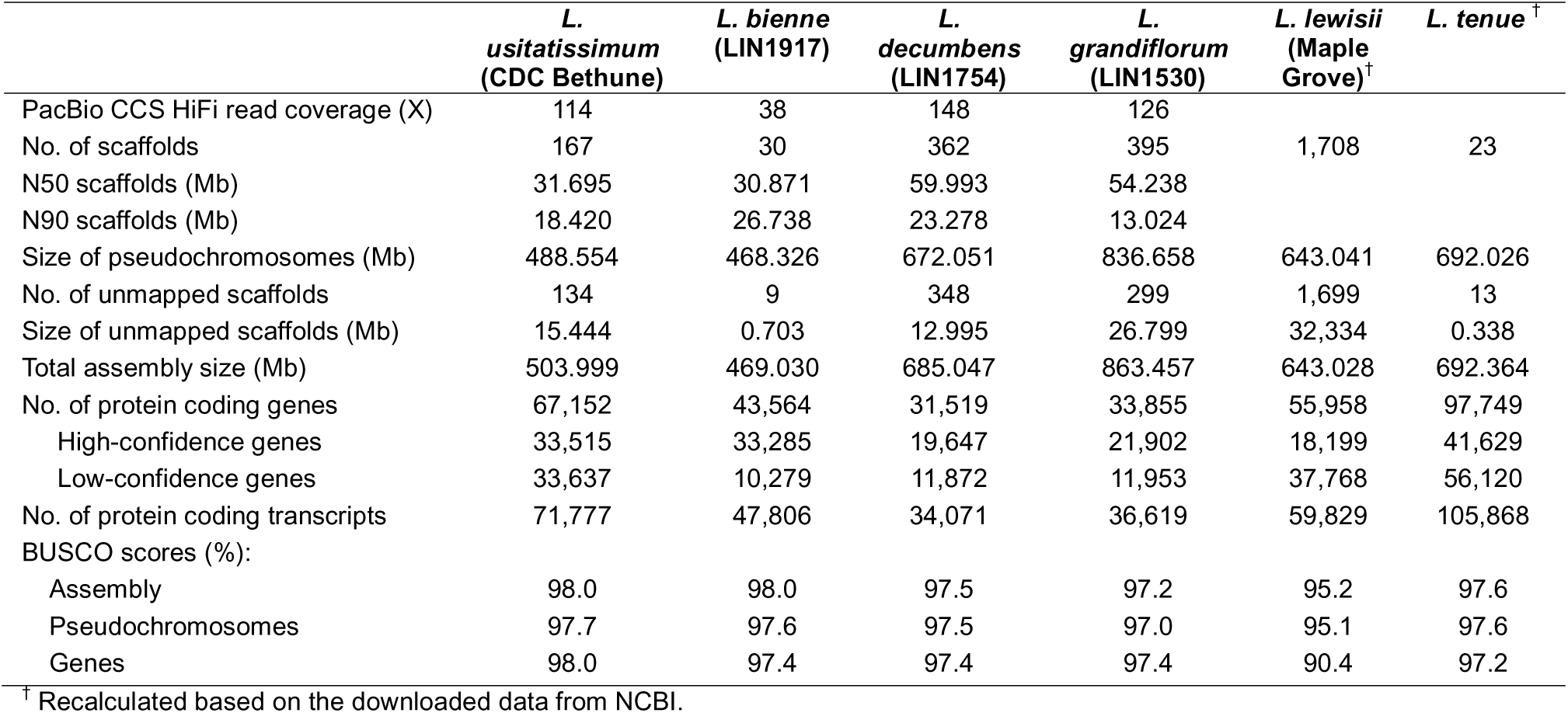
Genome assembly and gene prediction statistics for six *Linum* species.

Integration of BioNano optical maps (**Supplemental Table S2**) and three high-density genetic maps (**Supplemental Table S3; Supplemental Figures S3-S6**) enabled anchoring and orientation of a small number of scaffolds into chromosome-scale pseudomolecules for all species (**Supplemental Table S4**). Assembly completeness was supported by BUSCO scores exceeding 97% for all four species (**Table 2**; **Supplemental Figure S7**). Importantly, canonical telomeric repeat motifs (TTTAGGG) were detected at both termini of nearly all assembled pseudomolecules except for one arm of chromosome 3 of *L. decumbens* and both arms of chromosome 3 and one arm of chromosomes 7 and 8 of *L. grandiflorum* (**Figure 2**, **Track B**; **Supplemental Table S5**), providing evidence for near end-to-end chromosome continuity.

**Figure 2.**
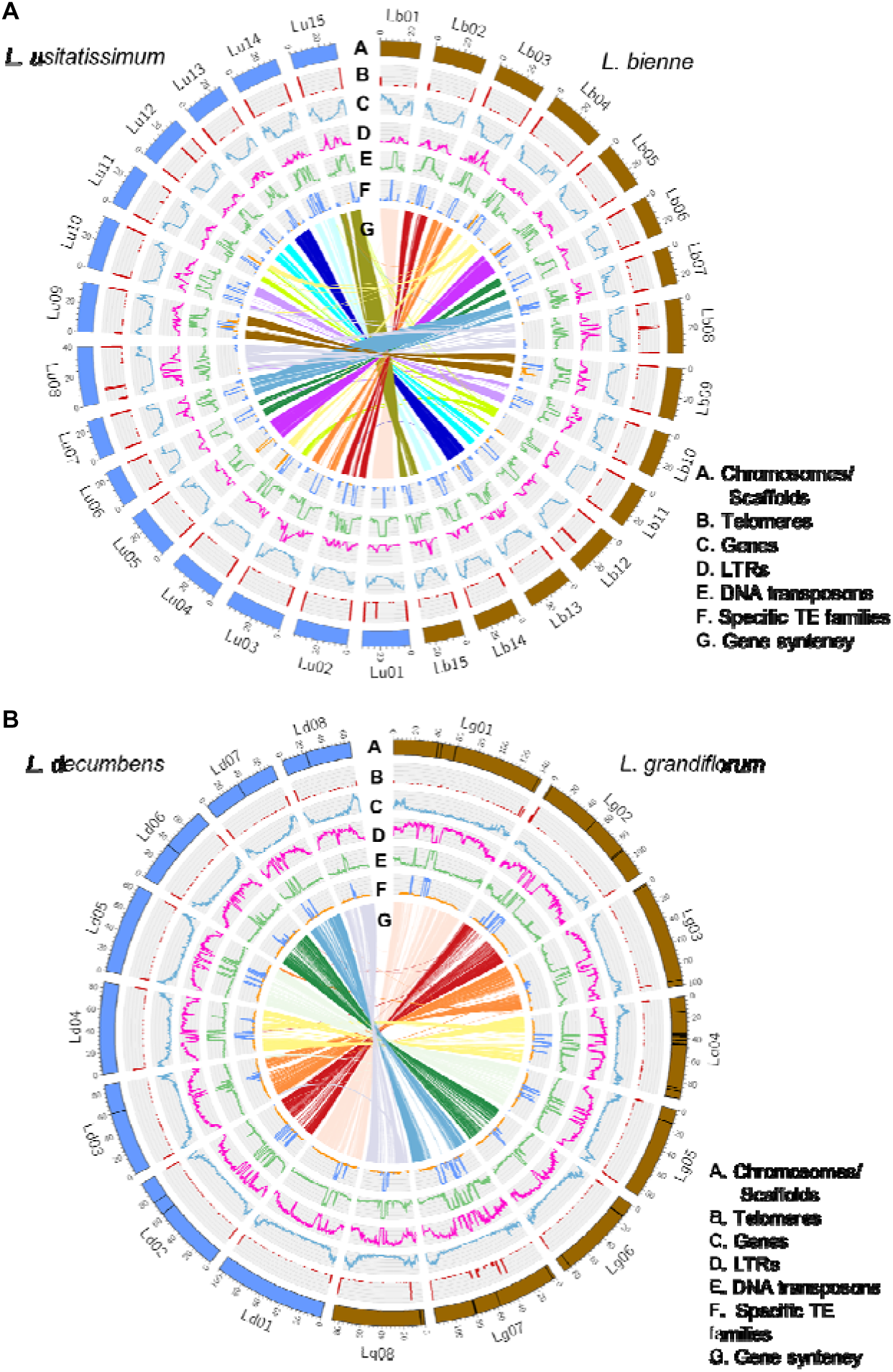
Comparative Circos maps reveal contrasting chromosome architectures and transposable element landscapes in *Linum*. (**A**) Circos map of the telomere-to-telomere genome assemblies for *L. usitatissimum* cv. CDC Bethune (left, blue) and *L. bienne* accession LIN1917 (right, brown). (**B**) Circos map of the telomere-to-telomere genome assemblies for *L. decumbens* accession LIN1754 (left, blue) and *L. grandiflorum* accession LIN1530 (right, brown). For each Circos map, concentric tracks represent: A, chromosomes with scaffold coordinates; B, telomeric regions identified by the simple repeat sequence TTTAGGG; C, gene density; D, long terminal repeat (LTR) transposable element (TE) density; E, DNA transposon density; F, density of predominant TE families (DNA transposon family *TE_00003234* and *TE_00000508* (second frequent) in the *n* = 15 lineage or *TE_00029583* in the *n* = 8 lineage); and G, gene collinearity (synteny) between chromosomes of the paired species.

For *L. usitatissimum* cv CDC Bethune, 15 near-T2T pseudomolecules were constructed, exhibiting strong collinearity between genetic map positions and physical coordinates (**Figure 2A; Supplemental Figures S3–S6**), thereby validating the assembly accuracy at the chromosome scale. Comparable pseudomolecule assemblies were obtained for *L. bienne*, *L. decumbens*, and *L. grandiflorum* (**Figure 2; Supplemental Table S4**), establishing a robust and unified framework for comparative genomic analyses across *Linum*.

These newly generated assemblies complement the previously published high-quality *Linum* genome assemblies of *L. tenue* (haploid chromosome number *n* = 10; assembly size 692.03 Mb) (Gutierrez-Valencia et al. 2022) and *L. lewisii* (*n* = 9; assembly size 643.04 Mb) (Innes et al. 2023) (**Table 2**). Both previously reported assemblies also exhibit a high degree of completeness, with BUSCO scores exceeding 95% (**Table 2**; **Supplemental Figure S7**), comparable to those obtained for the four genomes assembled here. Together, these six chromosome-level assemblies provide a comprehensive foundation for cross-species structural, evolutionary, and karyotypic analyses within *Linum*.

### Gene content, duplication history, and phylogeny provide evolutionary context

*Ab initio* gene prediction identified 31,519-67,152 protein-coding genes across the four newly assembled *Linum* genomes, with BUSCO completeness scores exceeding 97% for all gene sets (**Table 2**). Using the same annotation protocol, 55,958 genes were predicted in *L. lewisii* (Innes et al. 2023) whereas *L. tenue* harbored a substantially larger gene complement of 97,749 genes (Gutierrez-Valencia et al. 2022), consistent with its expanded duplication profile. Notably, *L. tenue* and *L. lewisii* exhibit highly similar gene structural features, including shorter gene lengths, fewer exons per gene, and higher proportions of single-exon genes compared with other *Linum* species, suggesting a shared gene architecture profile (**Supplemental Table S6**).

Analysis of the largest internal gene-free intervals of each chromosome revealed striking lineage-specific differences (**Supplemental Table S7**). *L. usitatissimum, L. bienne, L. decumbens* and *L. grandiflorum* have large internal gene-free regions across most chromosomes, typically spanning several megabases (on average 3.9-6.9 Mb across species), whereas *L. lewisii* and *L. tenue* largest internal gaps are thousand-fold smaller (102-407 kb). These results support the presence of expansive gene-poor pericentromeric regions in the lineage comprising *L. usitatissimum* and its closest related species, and indicate that genes are more continuously distributed across chromosomes in *L. lewisii* and *L. tenue*.

Orthogroup analysis across the six *Linum* genomes identified 40,115 orthogroups, revealing conserved and lineage-specific gene families (**Figure 3A–C**). Among these, 14,218 orthogroups were shared across all six species and constituted the conserved core gene set (**Figure 3C**; **Supplemental Table S8**), accounting for the largest proportion of gene copies in each genome, contrasting with accessory genes which showed substantial lineage-specific variation. Soft-core, shell, and rare orthogroups contributed moderately to genome composition, whereas private orthogroups were expanded in certain lineages, most notably in *L. tenue*. This species contained 27,173 private orthogroups comprising 55,489 gene copies, far exceeding those in the other species (**Figure 3A–B**; **Supplemental Table S8**). These results indicate that the large gene complement reported for *L. tenue* is likely attributable to lineage-specific gene expansion rather than proportional growth of conserved gene families. In addition, *L. usitatissimum* also exhibits significant gene expansion in private genes (10,389) compared to *L. bienne* (1,028) (**Supplemental Table S8**).

**Figure 3.**
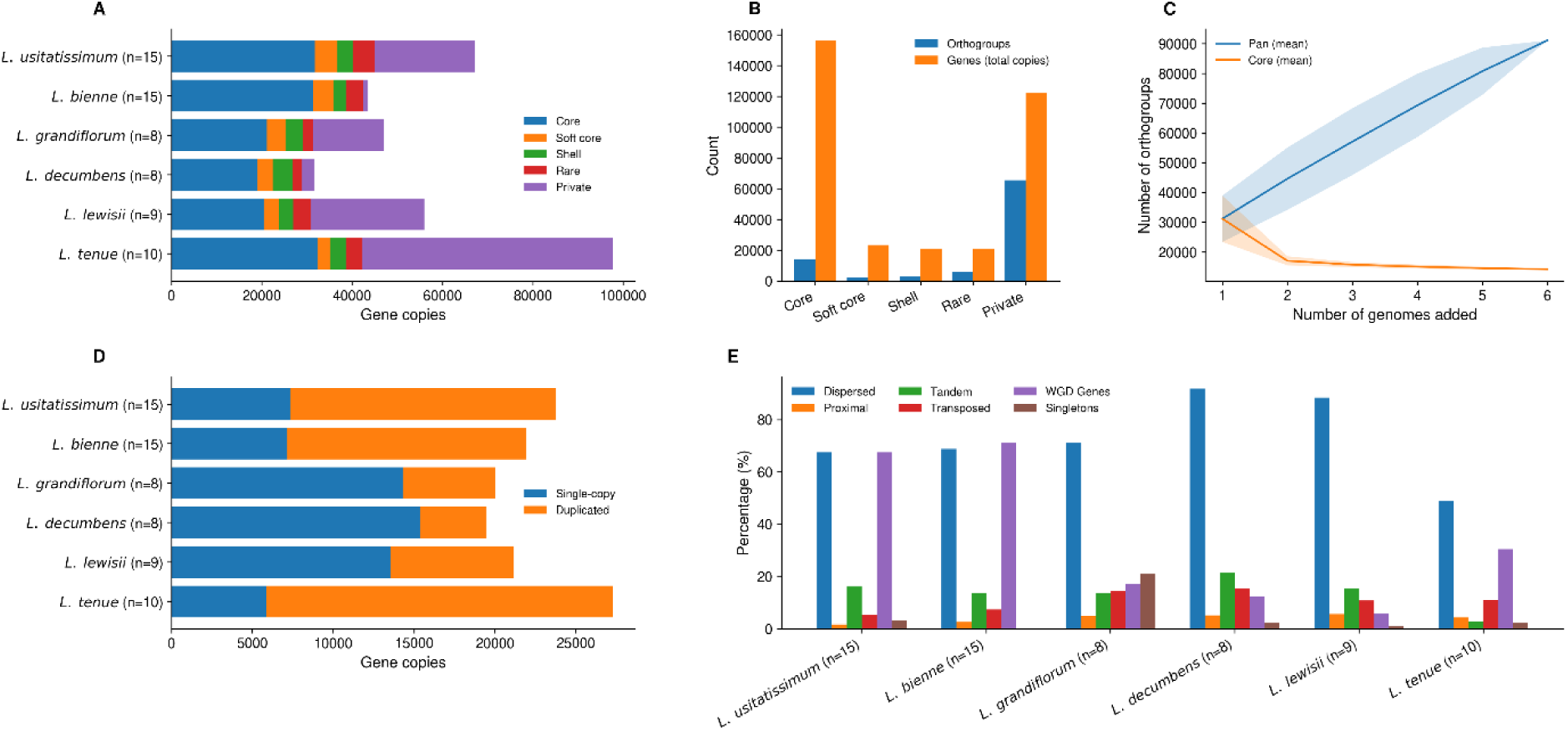
Pangenome structure and gene duplication patterns across six *Linum* species. (**A**) Gene composition of each genome partitioned into pangenome categories based on orthogroup prevalence across species (core, soft-core, shell, rare, and private). (**B**) Total numbers of orthogroups and gene copies within each pangenome category across the six species. **(C)** Pangenome accumulation curves showing the increase in total (pan) and conserved (core) orthogroups as additional genomes are sequentially incorporated. Lines represent mean values across random permutations of genome addition, with shaded areas indicating standard deviation. (**D**) Distribution of single-copy and duplicated orthogroups, illustrating variation in gene copy number across lineages. (**E**) Relative contributions of duplication modes, including whole-genome duplication (WGD), tandem, proximal, transposed, and dispersed duplication, inferred using DupGen_finder (Qiao et al. 2019).

Gene copy-number distributions further illustrate differences in duplication histories among species (**Figure 3D; Supplemental Table S9**). Species with higher chromosome numbers, including *L. bienne*, *L. usitatissimum*, and *L. tenue*, contained a larger proportion of duplicated orthogroups, whereas *L. decumbens*, *L. grandiflorum*, and *L. lewisii* had more single-copy orthogroups. Consistent with these patterns, BUSCO analysis revealed that duplicated core orthologs represented 70-77% of the *L. tenue*, *L. bienne*, and *L. usitatissimum* assemblies (**Supplemental Table S10**).

Gene duplication mode analysis using DupGen_finder (Qiao et al. 2019) revealed that both WGD and dispersed duplication contributed the largest fractions of duplicated genes in the *n* = 15 and *n* = 10 lineages, while only dispersed duplication predominated in the *n* = 8 and *n* = 9 lineages. Tandem, proximal, and transposed duplications contributed relatively minor fractions (**Figure 3E**), highlighting substantial variation in gene duplication patterns across *Linum* lineages.

Functional annotation further highlights the evolutionary nature of lineage-specific genes. While core genes exhibited relatively high GO annotation rates (∼54–57%), fewer private genes (3–11%) had GO annotations (**Supplemental Table S8**), indicating that many lineage-specific genes either lack recognizable functional domains or represent rapidly evolving or poorly characterized gene families. In *L. tenue* and *L. usitatissimum*, only ∼6-8% of the private gene copies were associated with GO annotations, supporting that its expanded gene content largely reflects lineage-specific gene duplication and diversification rather than expansion of conserved functional gene families.

Phylogenomic analysis based on 908 near single-copy nuclear protein-coding genes resolved well-supported relationships among the six species (**Figure 4**). Divergence times were estimated under a relaxed molecular clock model implemented in MCMCTree (PAML v4.10.9) (Yang 2007), using a calibration at the crown node of *Linum* (split between the yellow-flowered lineage, represented by *L. tenue*, and the blue-flowered lineage) at 38.02 MYA (95% highest posterior density (HPD): 36.17–39.55 MYA). The analysis placed *L. tenue* as the basal lineage, followed by *L. lewisii*, which diverged from the common ancestor of the *n* = 8 and *n* = 15 lineages approximately 25–27 MYA. Within the *n* = 8 clade, *L. decumbens* and *L. grandiflorum* diverged approximately 8 MYA, whereas *L. bienne* and *L. usitatissimum* formed a well-supported sister pair that separated approximately 0.7–1 MYA.

**Figure 4.**
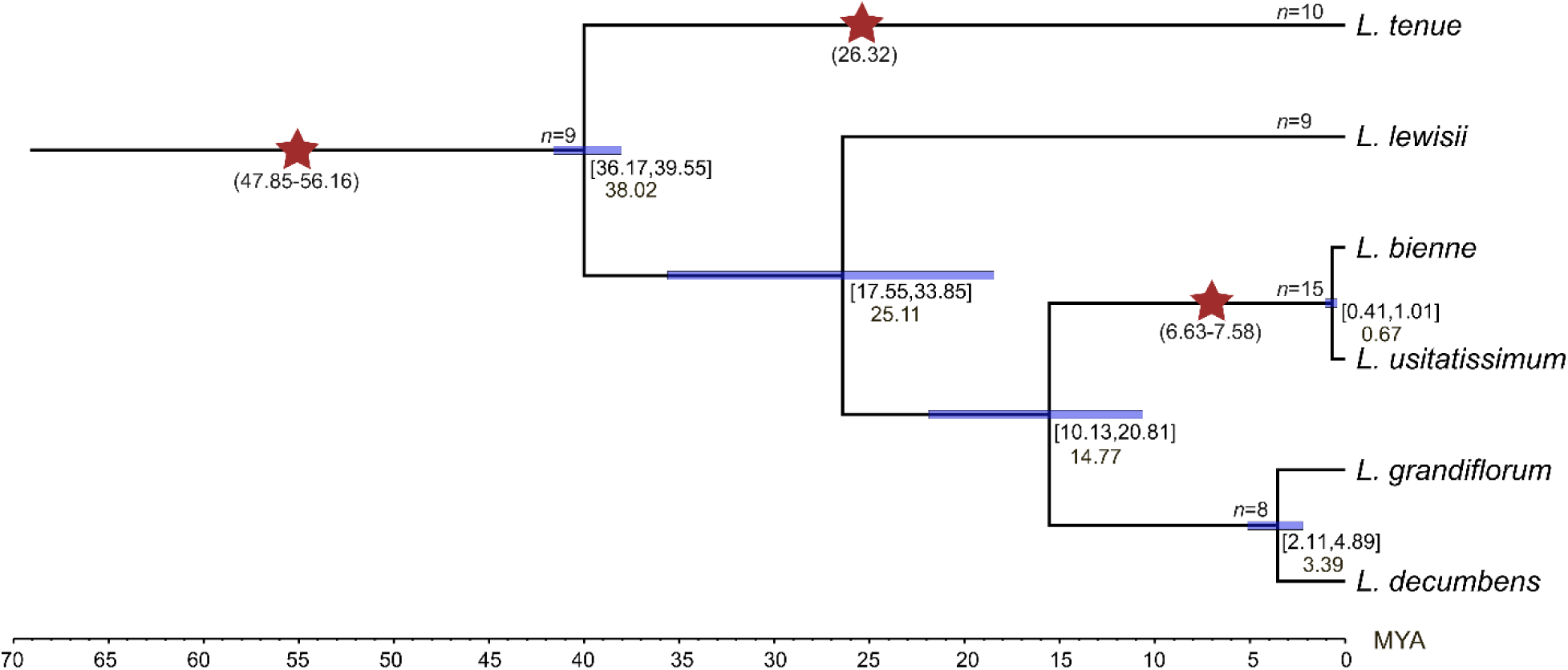
Phylogenetic relationships and divergence times of six *Linum* species. A maximum-likelihood phylogenetic tree was reconstructed using 908 single-copy nuclear protein-coding genes from six *Linum* species assemblies, with *L. tenue* designated as the outgroup. Divergence times and their 95% highest posterior density (HPD) intervals (blue bars and values in square brackets) were estimated under a relaxed molecular clock model implemented in MCMCTree (PAML v4.10.9). The *Linum* crown node was calibrated at 38.02 MYA, with a 95% HPD interval of 36.17–39.55 MYA, based on Villalvazo-Hernandez et al. (2022). Whole-genome duplication (WGD) events and their timing were inferred using WGD v2.0. Inferred WGDs are indicated by red stars, with estimated time of duplication (MYA) shown below in brackets.

WGD events were inferred using WGD v2.0 (Chen et al. 2024) (**Supplemental Table S11**; **Figure 4**). An ancient WGD event was detected across all six species, with estimated times ranging from approximately 48 to 56 MYA, predating the diversification of the *Linum* genus. In addition, lineage-specific WGDs were identified in species with expanded chromosome numbers. Both *L. bienne* and *L. usitatissimum* exhibited recent WGD events dated at ∼7.6 and ∼6.6 MYA, respectively, while *L. tenue* showed a distinct WGD event at ∼26.3 MYA.

A sharp and narrow *Ks* peak was observed in *L. grandiflorium* (∼3.1 MYA; corresponding to a *Ks* mode of ∼0.03) under default peak-detection parameters (**Supplemental Figure S8C**). Similar minor peaks at *Ks* ≈ 0.03 were also detected in *L. decumbens*, *L. lewisii*, and *L. tenue* when the peak-detection threshold (-prct) was relaxed to 0.01 (not labeled in the figure). However, these weak and parameter-sensitive peaks may reflect recent small-scale duplications, residual heterozygosity, or assembly-related artifacts rather than independent lineage-specific WGD events. Collectively, these analyses integrate gene content, duplication modes, and temporal phylogenetic relationships into a coherent evolutionary framework that supports an ancient shared WGD preceding divergence of the species, followed by subsequent lineage-specific polyploidization and differential duplicate retention contributing to chromosome number variations and gene repertoire expansion across *Linum* species.

### A single DNA transposon family dominates pericentromeric regions in the flax lineage

We constructed a pangenomic transposable element (TE) library comprising 31,159 TE families spanning the six *Linum* species. Genome-wide TE annotation using this library revealed that TEs accounted for 63–84% of *Linum* genomes (**Supplemental Table S12**). Species with eight to ten chromosomes (*L. decumbens*, *L. grandiflorum*, *L. lewisii*, and *L. tenue*) were dominated by Class I LTR retrotransposons, with the most abundant being those of the *Gypsy* superfamily in total and intact LTRs (**Figure 5A**; **Supplemental Table S12; Supplemental Figure S9**). LTR elements accounted for 19.51% of the genome in *L. tenue* to 34.64% in *L. decumbens*. In contrast, *L. bienne* and *L. usitatissimum* harbored fewer Class I retrotransposons and a marked expansion of Class II DNA transposons, which alone accounted for more than one-third of their genomes, i.e., 35.10 and 36.12%, respectively (**Supplemental Table S12**). Consistent TE landscapes were also observed across additional flax (*L. usitatissimum*) cultivars, including the published assemblies of T397 (Yadav et al. 2025), Gaosi (Lu et al. 2025), and Atlant (Dmitriev et al. 2020) (**Supplemental Figure S10A**). In contrast, representative grass genomes, including wheat, *Aegilops tauschii*, barley, and *Brachypodium*, are predominantly composed of Class I LTR retrotransposons, with DNA transposons contributing a much smaller fraction (**Supplemental Figure S10B**).

**Figure 5.**
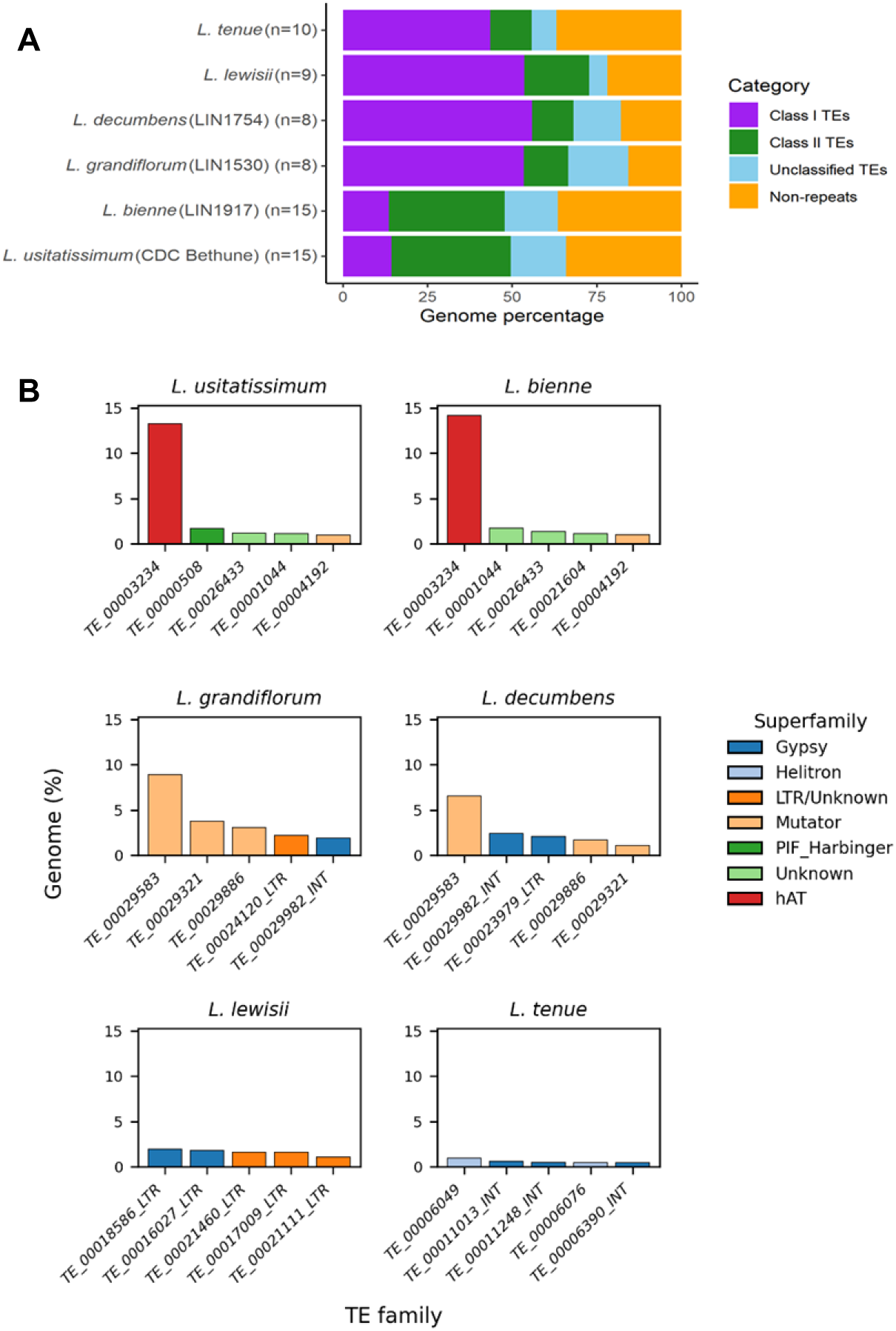
Genome-wide composition of transposable elements across six *Linum* species. (**A**) Stacked bars show the percentage contribution of Class I retrotransposons, Class II DNA transposons, unclassified TEs, and non-repetitive sequences to the total genome sizes. (**B**) Lineage-specific enrichment of major transposable element families in six *Linum* genomes. The top five TE families in each species are ranked by genomic contribution (% of genome). Bar colors represent TE superfamilies, illustrating differences in the dominant TE types among lineages.

This contrasting TE composition within the *n* = 15 lineage was primarily driven by the DNA transposon family *TE_00003234* **(Figure 5B; Supplemental Figure S10A).** This *hAT* element was characterized by 8-bp target site duplications (TSDs) and ∼8–12 bp terminal inverted repeats (TIRs), with an internal sequence spanning approximately 1.1–1.35 kb (**Supplemental Figure S11**). This single family accounted for ∼14% of the genomes of *L. bienne* and *L. usitatissimum*, but was nearly absent from the other *Linum* species analyzed (**Table 3**; **Figure 5B**). Circos map showed that *TE_00003234* copies were highly enriched in large central chromosomal regions, which collectively constituted 28–60% of individual chromosomes. These regions coincided with intervals of low gene density and extremely low nucleotide variation.

**Table 3.** Most frequent transposable elements in six *Linum* species.

| Species | Accession | TE family | TE class/<br>superfamily | Consensus<br>TE family<br>length (bp) | Total<br>blast<br>hits | Total<br>bases<br>(Mb) | Genome<br>(%) | Mean<br>divergence<br>(K) | Average<br>insertion<br>time (MY) ±<br>Std |
| --- | --- | --- | --- | --- | --- | --- | --- | --- | --- |
| *L. tenue* | | *TE_00011013_INT* | LTR/Gypsy | 3,465 | 708 | 4.30 | 0.62 | 0.09 | 3.42 ± 1.45 |
| *L. lewisii* | Maple<br>Grove | *TE_00018586_LTR* | LTR/Gypsy | 1,518 | 6,134 | 12.67 | 1.97 | 0.15 | 5.44 ± 1.95 |
| *L. decumbens* | LIN1754 | *TE_00029583* | TIR/*Mutator* | 351 | 572 | 44.95 | 6.56 | 0.27 | 10.25 ± 1.28 |
| *L. grandiflorum* | LIN1530 | *TE_00029583* | TIR/*Mutator* | 351 | 1,244 | 76.89 | 8.91 | 0.28 | 10.53 ± 1.32 |
| *L. bienne* | LIN1917 | *TE_00003234* | TIR/*hAT* | 1,266 | 1,406 | 66.38 | 14.15 | 0.31 | 11.87 ± 1.91 |
| *L. usitatissimum* | CDC<br>Bethune | *TE_00003234* | TIR/*hAT* | 1,266 | 1,571 | 66.97 | 13.29 | 0.31 | 11.75 ± 1.94 |

Std: standard deviation

Although *L. decumbens* and *L. grandiflorum* were overall dominated by Class I TEs, the Mutator DNA transposon family *TE_00029583* was the most abundant individual family in these species, with 572 and 1,244 hits, accounting for 6.6 % and 8.9% of their genome, respectively (**Table 3**; **Figure 5B**). Conversely, *TE_00029583* was absent from the *L. lewisii* and *L. tenue* genome assemblies, and rare (only three hits each) in *L. bienne* and *L. usitatissimum*. No dominant TE family was identified in the *L. lewisii* or *L. tenue* genomes, where top five TE families individually accounted for less than 2% of the genome and consisted primarily of Class I LTR retrotransposons.

Insertion time distributions of the most abundant TE families revealed distinct lineage-specific amplification histories (**Figure 6**; **Table 3**). In *L. tenue* and *L. lewisii*, dominant LTR*/Gypsy* families exhibited recent insertions at approximately 3–6 MYA, suggesting ongoing or comparatively recent transposition activity. In *L. decumbens* and *L. grandiflorum*, amplification of the DNA transposon family *TE_00029583* (TIR/Mutator) is dated to mean insertion times around 10–11 MYA.

**Figure 6.**
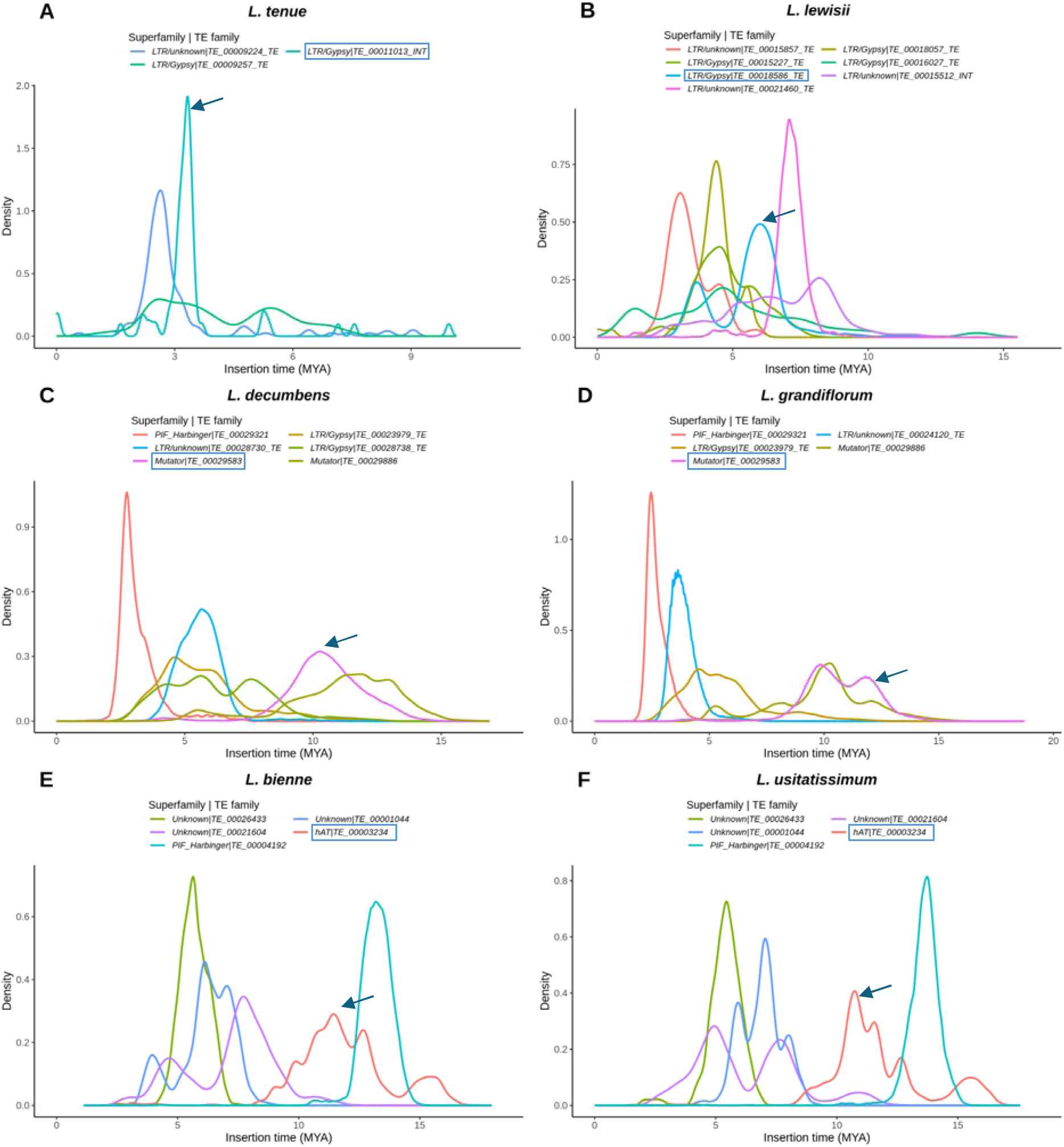
Insertion times in million years ago (MYA) of major transposable element (TE) families in *six Linum* species. (**A**) *L. tenue*, (**B**) *L. lewisii* accession Maple Grove, (**C**) *L. decumbens* accession LIN1754, (**D**) *L. grandiflorum* accession LIN1530, (**E**) *L. bienne* accession LIN1917, and (**F**) *L. usitatissimum* cultivar CDC Bethune. Predominant TE families are highlighted with blue boxes and their corresponding peaks are indicated by blue arrows. Curves represent kernel density estimates of TE insertion times and their density values are normalized with out unit so that the area under each curve equals 1.

In the *L. bienne* and *L. usitatissimum* lineage, the dominant *TIR/hAT* family *TE_00003234* exhibited a concentrated insertion peak corresponding to *K* ≈ 0.30, translating to approximately 10–12 MYA (**Figure 6E–6F**), indicating that its major amplification occurred after the divergence of the *n* = 15 lineage from its closest relatives *L. decumbens* and *L. grandiflorum* (14.77 MYA; **Figure 4**).

Although the divergence peak (∼10–12 MYA) predates the most recent WGD event in the *n* = 15 lineage (∼6–7 MYA), direct temporal comparisons between TE divergence (*K*) and gene-based synonymous substitution (*Ks*) should be interpreted cautiously. Copy-to-consensus divergence time estimates in repetitive elements (Rodriguez and Arkhipova 2023; Zhang et al. 2023) integrate mutations accumulated across multiple amplification waves and may inflate apparent time of divergence relative to that obtained from single-copy gene comparisons. In addition, divergence-based estimates in repetitive elements are sensitive to consensus reconstruction and mutation-rate heterogeneity, and therefore should be interpreted as approximate temporal signals rather than precise chronological events. Therefore, we interpret the *TE_00003234* burst as broadly occurring during the early evolutionary phase of the *n* = 15 lineage, potentially overlapping with or preceding the polyploidization and subsequent chromosomal reorganization, rather than asserting strict chronological ordering relative to the recent WGD event.

Together, these results illustrate the distinct TE amplification episodes of each *Linum* lineage, namely the *Mutator* expansion in the *n* = 8 lineage (∼10 MYA), the *hAT* expansion in the *n* = 15 lineage (∼10–12 MYA), and the more recent LTR retrotransposon activity in the *n* = 9 and *n* = 10 lineages. These distinct TE amplification histories contributed to lineage-specific genome restructuring and chromosome evolution across the genus.

### DNA transposon-rich central chromosomal domains are strongly depleted in genes and SNPs and are associated with segregation distortion

To characterize the genomic features of TE-rich central regions in flax, we examined gene density, SNP density, and short-read mapping depth along each chromosome using resequencing data from 407 accessions mapped to the CDC Bethune v3.0 reference genome. Across all 15 chromosomes, extended central regions consistently exhibited markedly reduced gene and SNP densities relative to chromosome arms (**Figure 7**). These regions often span several megabases and contain few annotated genes and extremely low SNP density (nucleotide diversity), despite read depth comparable to that of chromosome arms, indicating that the observed depletion is unlikely the result of mapping bias or assembly artifacts.

**Figure 7.**
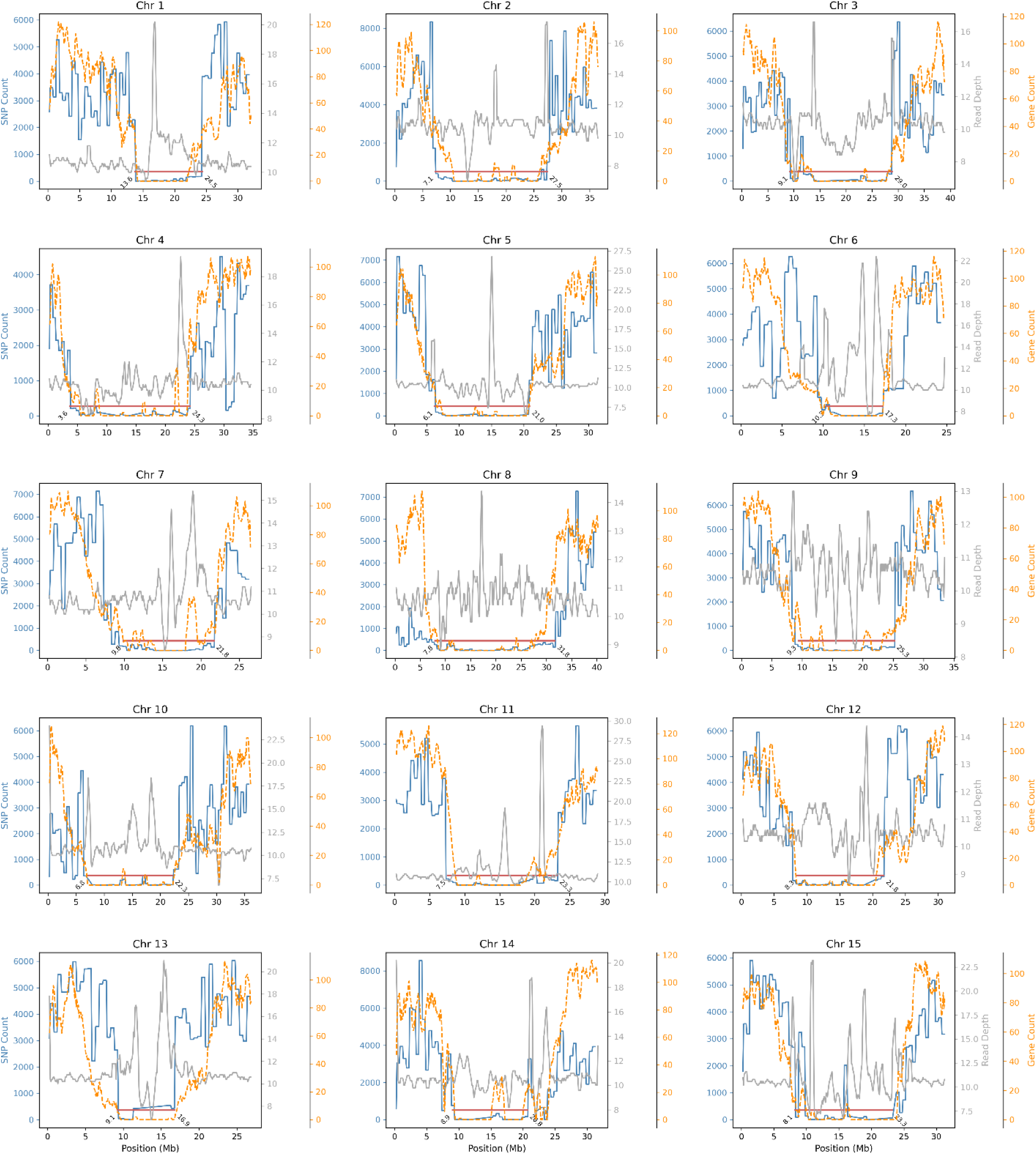
Central repeat-rich chromosomal domains in flax are exceptionally depleted in genes and SNPs despite broadly uniform read depth. Distribution of SNP counts (blue), gene counts (orange), and read depth (gray), along the 15 chromosomes of *L. usitatissimum* cv. CDC Bethune. Read depth represents the average number of Illumina short reads from a core collection of 407 accessions mapped to CDC Bethune v3.0. SNPs were identified from the same set of accessions using the same reference genome. Red horizontal lines indicate contiguous chromosomal regions of low SNP and gene densities. Numbers adjacent to each end of the red line mark the approximate start and end coordinates of these low SNP and gene regions for each chromosome. Values were calculated using sliding windows of 100 kb with a 10 kb step.

The scale of these low-density regions is consistent with the large internal gene gaps summarized in **Supplemental Table S7**, where maximum distances between flanking genes average approximately 4.5–5.5 Mb across chromosomes in the n = 15 lineage. Comparable gap sizes were also observed in *n* = 8 species, suggesting that large gene-poor intervals are a general feature of *Linum* chromosomes, although the underlying sequence composition differs among lineages.

These low-SNP/low-gene-density regions largely coincide with *TE_00003234*-enriched domains identified in the Circos maps (**Figure 2A**). To quantify this relationship, we compared *TE_00003234* distribution between low-density regions and the remainder of the genome. *TE_00003234* occupies a substantial fraction of sequence within low-density regions, reaching ∼20–85% coverage across chromosomes, while remaining low in the other regions (typically <10%) (**Supplemental Table S13**; **Supplemental Figure S12A**). Consistently, strong enrichment was observed across most chromosomes, with enrichment ranging from ∼8- to >30-fold, except for chromosomes 5, 8, and 15, which contain fewer *TE_00003234* elements (**Supplemental Table S13**; **Supplemental Figure S12B**).

To test whether this enrichment reflects non-random genomic organization, we performed chromosome-wise permutation analyses that preserve interval size and number. In all cases, observed TE coverage in low-density regions exceeded permutation-based expectations (**Supplemental Figure S12C**), indicating that the concentration of *TE_00003234* is unlikely to arise by chance. Together, these results demonstrate that *TE_00003234* is highly and non-randomly concentrated within these regions.

To assess potential genetic consequences, we examined segregation distortion in two *L. usitatissimum* × *L. usitatissimum* recombinant inbred line (RIL) populations and one interspecific *L. bienne* × *L. usitatissimum* RIL population (**Supplemental Table S3**). Across all populations, elevated distortion signals were consistently detected in central chromosomal regions, whereas chromosome arms showed predominantly background levels (**Supplemental Figures S13–S16**). Regions of elevated distortion frequently overlapped intervals of extremely low SNP density within *TE_00003234*-enriched domains.

Taken together, these results show that flax chromosomes contain large central regions characterized by low gene density, extremely low SNP density, and high repeat content. These regions are strongly enriched for the lineage-specific DNA transposon *TE_00003234* and are consistently associated with segregation distortion across multiple populations.

### Contrasting TE landscapes correlate with karyotype divergence

Comparative analysis of TE family composition across six *Linum* species revealed strong correlations among closely related species. *L. decumbens* and *L. grandiflorum* had similar TE composition (*r* = 0.92), followed by *L. bienne* and *L. usitatissimum* (*r* = 0.87) (**Supplemental Figure S17**). In contrast, *L. lewisii* and *L. tenue* displayed distinct TE profiles, consistent with independent evolutionary histories. These patterns indicate that chromosome evolution in *Linum* involved shifts not only in chromosome number, but also in dominant TE classes and expansion dynamics. Species retaining low chromosome numbers had LTR-dominated genomes, whereas the lineages with 15-chromosomes experienced lineage-specific DNA transposon expansion that reshaped their chromosome architecture.

### Chromosome rearrangements are concentrated in TE-rich regions

Comparative gene synteny and sequence collinearity analyses revealed extensive conservation of gene order between *L. decumbens* and *L. grandiflorum*, punctuated by several large inversions and localized translocations (**Figure 2B**; **Supplemental Figure S18**). In contrast, comparisons between *L. bienne* and *L. usitatissimum* (**Figure 2A**; **Supplemental Figure S19**) revealed widespread chromosome fragmentation and reorganization. Notably, alignments between *L. decumbens* or *L. grandiflorum* and *L. bienne* or *L. usitatissimum* showed little or no synteny within the central chromosomal regions enriched for *TE_00003234* (**Supplemental Figures S20–S23**). These TE-dominated regions disrupted the synteny that likely suppressed recombination, thereby facilitating chromosome breakage and rearrangement during karyotype evolution.

### Chromosome restructuring underlying the transition from *n* = 8 to *n* = 15

To investigate the structural basis of the chromosome number transition from *L. decumbens* (*n* = 8) to *L. usitatissimum* (*n* = 15), genome-wide gene synteny and collinearity analyses were performed. The combined visualization of orthologous gene distribution and pairwise dot plots revealed extensive fragmentation and redistribution of ancestral chromosomal segments across the derived *L. usitatissimum* karyotype (**Supplemental Figure S20**; **Figure 8**). Rather than exhibiting a simple one-to-one correspondence or whole-chromosome duplication pattern, each *L. usitatissimum* chromosome was composed of syntenic blocks derived from multiple *L. decumbens* chromosomes, forming a pervasive many-to-many relationship.

**Figure 8.**
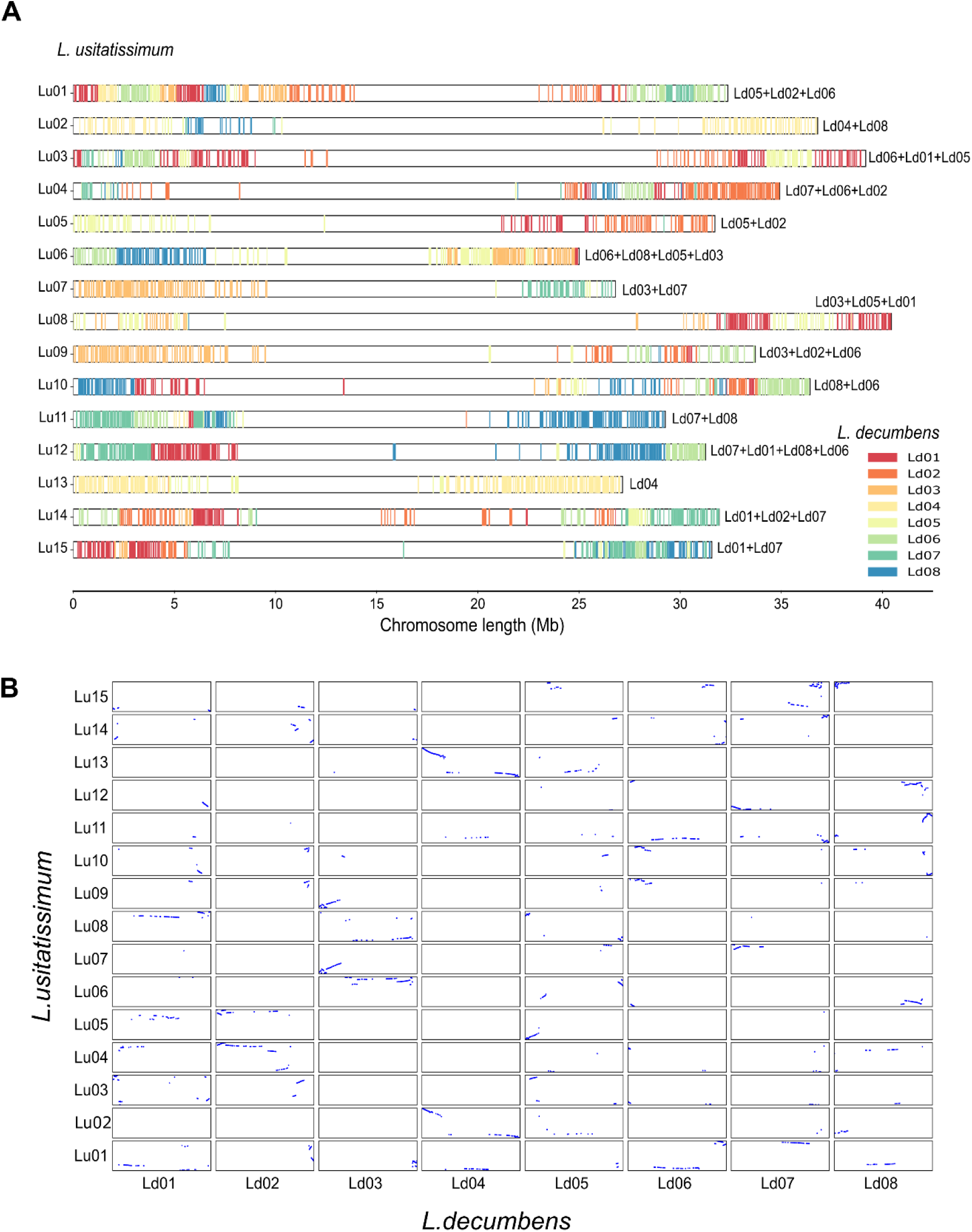
Distribution and collinearity of orthologous genes between *L. usitatissimum* and *L. decumbens*. (**A**) Chromosomal distribution of *L. decumbens* orthologs along the chromosomes of *L. usitatissimum*. Each horizontal line represents one *L. usitatissimum* chromosome, and colored ticks indicate the positions of orthologous genes, with colors corresponding to their chromosome of origin in *L. decumbens* (Ld01–Ld08). Multiple labels at chromosome termini denote the major contributing *L. decumbens* chromosomes. (**B**) Dot plot showing pairwise collinearity between *L. usitatissimum* (y-axis) and *L. decumbens* (x-axis) chromosomes based on orthologous gene pairs. Blue dots represent collinear gene matches.

The dot plot analysis further demonstrates that collinear regions are highly discontinuous and dispersed, with no evidence of intact chromosome-scale conservation between the two genomes (**Figure 8B**). Most *L. usitatissimum* chromosomes appear to be mosaics, typically combining segments derived from two or more ancestral chromosomes. For example, Lu07 contains major segments from Ld03 and Ld07, whereas Lu15 is primarily derived from Ld01 and Ld07. In contrast, only a small subset of chromosomes retain relatively continuous ancestry from a single *n* = 8 chromosome. In particular, Lu02 and Lu13 both preserve substantial collinear segments originating from Ld04, representing the closest approximation to chromosome-scale conservation observed in the *n* = 15 lineage.

Similar patterns of extensive chromosomal fragmentation and reassortment were consistently observed in additional comparisons, including *L. bienne* versus *L. decumbens* (**Supplemental Figures S21, S24**), *L. usitatissimum* versus *L. grandiflorum* (**Supplemental Figure S22, S25**), and *L. bienne* versus *L. grandiflorum* (**Supplemental Figure S23, S26**). These concordant results indicate that the observed restructuring is not specific to a single species pair but reflects a general feature of chromosome evolution within the *n* = 15 lineage.

Together, these results do not support a simple model of whole-genome duplication followed by limited chromosome loss. Instead, the pervasive intermixing of ancestral segments and the absence of chromosome-scale collinearity are consistent with extensive post-polyploid diploidization involving repeated chromosome fission, fusion, and interchromosomal translocation events. This large-scale genome reorganization underlies the increase and reshaping of chromosome number in the *n* = 15 *Linum* species and highlights a decoupling of chromosome number from genome size and overall chromosome length.

## DISCUSSION

### Near telomere-to-telomere reference genome assemblies resolve the genomic architecture of chromosome evolution in *Linum*

The near-T2T genome assemblies generated in this study provide a comprehensive and contiguous view of chromosome organization in *Linum*, overcoming a long-standing limitation in plant genomics: the fragmented representation of repetitive centromeric and pericentromeric regions in short-read assemblies (Nurk et al. 2022; Miga et al. 2020). Complete chromosome sequences are essential for resolving the structural dynamics underlying karyotype evolution, particularly in lineages characterized by extensive chromosomal reorganization (Schubert and Lysak 2011).

Our near-T2T assembly of cultivated flax (*L. usitatissimum* cv. CDC Bethune; v3.0) represents a substantial improvement over the previous chromosome-scale reference genome (v2.0) (You et al. 2018b), which comprised 316 Mb of anchored sequence and contained unresolved gaps and several misoriented scaffolds (**Supplemental Figure S27**), thereby limiting interpretation of central chromosome regions. By integrating PacBio HiFi sequencing, optical mapping, and multiple high-density genetic maps, we resolved all 15 chromosomes end-to-end, including telomeric and pericentromeric domains. Telomeric repeats were detected at both ends of all pseudomolecules, and strong collinearity between genetic and physical maps validated the structural accuracy of the assembly. Consistent read-mapping rates across hundreds of accessions (**Supplemental Figure S28**) further support the degree of completeness of the assembly and establish CDC Bethune v3.0 as the superior reference genome for the flax research community.

Comparable near-T2T assemblies for *L. bienne*, *L. decumbens*, and *L. grandiflorum* enabled unbiased cross-lineage comparisons of chromosome architecture. These assemblies reveal that major interspecific structural differences are concentrated in and around the centromeric regions, underscoring the importance of T2T contiguity for resolving the structural bases of chromosome evolution.

### Diversification of *Linum* karyotypes through ancestral polyploidy and lineage-specific evolutionary trajectories

Whole genome duplication provides genomic redundancy that permits chromosome number evolution, but WGD alone does not determine the final karyotype. Classical models of post-polyploid diploidization emphasize chromosome fusion, fission, and segmental rearrangements as key mechanisms shaping chromosome number after genome doubling (Mandáková and Lysak 2018; Schubert and Lysak 2011). The evolution of some *Linum* species falls within this framework while also highlighting lineage-specific genomic processes accompanying these structural changes.

Our analyses indicate that extant *Linum* karyotypes originated from a shared ancestral WGD dating ∼48–56 MYA, likely associated with the establishment of the ancestral *n* = 9 chromosomes. This event predates the diversification of the major lineages and represents a foundational step within the genus. Previous studies inferred an ancient WGD at ∼20–44 MYA, but these were based on fragmented assemblies or limited gene sequences (Sveinsson et al. 2014; You et al. 2018b). The multi-species near-T2T analysis presented here suggests an earlier polyploid origin than previously estimated.

Subsequent karyotype evolution followed distinct lineage-specific trajectories. In the lineage leading to cultivated flax (*L. usitatissimum*) and its wild progenitor (*L. bienne*), a more recent WGD (∼6–7 MYA) occurred after divergence from the *n* = 8 lineage (∼14–15 MYA) and is associated with the emergence of the *n* = 15 karyotype. The nearly identical WGD timing and structural configurations of these two species support a shared polyploid origin preceding domestication.

In contrast, *L. tenue* experienced an independent and older lineage-specific WGD (∼26 MYA) associated with a modest increase in chromosome number from *n* = 9 to *n* = 10. Despite this chromosome-number increase, *L. tenue* does not exhibit the pronounced genome-size expansion typically expected following polyploidization. Instead, its genome contains an unusually large number of lineage-specific genes, many of which occur as duplicated copies and lack functional annotation. This pattern suggests that the *L. tenue* lineage followed a distinct post-WGD evolutionary trajectory characterized by extensive retention or expansion of lineage-specific gene copies without corresponding increases in overall genome size. Such a pattern contrasts with the *L. usitatissimum* lineage, where chromosome-number increase to *n* **=** 15 was accompanied by extensive TE-associated chromosomal restructuring and genome downsizing rather than expansion of the gene content. Other species, such as *L. lewisii* (*n* = 9), *L. decumbens* and *L. grandiflorum* (*n* = 8), show no evidence of recent lineage-specific WGDs. In these species, chromosome number reduction likely reflects descending dysploidy through chromosome fusion and structural rearrangement following the ancestral WGD rather than additional duplications. These contrasting outcomes illustrate that polyploidy in *Linum* has been succeeded by multiple diploidization trajectories, consistent with broader models of post-polyploid genome evolution in plants in which gene retention, genome fractionation, and structural reorganization proceed at different rates among lineages. The pangenome analysis further supports these disparate evolutionary trajectories, as *L. tenue* exhibits a striking expansion of duplicated private genes (**Figure 3**), whereas the *L*. *usitatissimum*/*L*. *bienne* lineage maintains comparable gene numbers despite extensive chromosome restructuring.

### Chromosome number increases without genome expansion: evidence for extensive internal reorganization

Chromosome number evolution in plants is commonly associated with WGD followed by genome expansion or contraction (Mandáková and Lysak 2018; Stebbins 1950). However, our analyses indicate that the transition from *n* = 8 to *n* = 15 in the *Linum* lineage occurred without proportional genome expansion and was incongruously accompanied by a reduction in total genome size. This decoupling of chromosome number, total chromosome length, and genome size suggests that karyotype evolution in this *Linum* lineage did not follow a simple duplication-based trajectory. Instead, the data support a model of extensive internal reorganization involving chromosome fragmentation, fusion, and reassortment that reshaped chromosome architecture without increasing genome size. Similar dysploid processes have been described in other plant lineages (Schubert and Lysak 2011; Mandáková and Lysak 2018), but in *Linum* these changes appear to be intricately associated with lineage-specific remodeling of repetitive elements within pericentromeric regions.

### A lineage-specific DNA transposon dominates pericentromeric regions

LTR retrotransposons are widely recognized as the principal drivers of pericentromeric expansion and structural diversification in plant genomes (Bennetzen and Wang 2014; Vitte and Bennetzen 2006). Increasing evidence indicates that DNA transposons can also contribute to structural genome evolution across diverse lineages (Huang et al. 2012). Substantial amplification of DNA transposons has been documented in several animal genomes. For example, in the invertebrate *Hydra magnipapillata*, DNA transposons account for ∼21% of the genome, representing approximately half of all repeat sequences, with *hAT* and *Mariner* elements together contributing ∼10% of the genome (Chapman et al. 2010). In the brown bat *Myotis lucifugus*, lineage-specific amplification of Helitron elements constitutes roughly 3.4% of the genome (Pritham and Feschotte 2007). Similar examples have been reported in plant genomes. In wheat, *CACTA* elements are enriched at chromosomal breakpoints following polyploidization (Wicker et al. 2018). The novel DNA transposon superfamily “GingerRoot” was identified in *Selaginella lepidophylla*, where it accounts for 1.7% of the genome (Cerbin et al. 2019). In these cases, DNA transposons generally function as local mutagens or facilitators of chromosomal rearrangements within genomes that are otherwise dominated by LTR retrotransposons.

Previous analyses of *L. usitatissimum* TEs suggested that LTR retrotransposons dominate the repeat landscape (Duk et al. 2025; Demenou et al. 2025; Yadav et al. 2025; Lu et al. 2025). However, these observations were derived from fragmented assemblies that lacked complete pericentromeric regions or resulted from incomplete repeat annotation methods. Our results revealed a striking departure from these reports. In the *L. usitatissimum* and *L. bienne* lineages, Class II DNA transposons occupy ∼35% of the genomes, corresponding to 2.5-fold more than Class I retrotransposons (∼14%) (**Supplemental Table S12**). This finding revises the current view of repeat composition in these genomes and highlights the importance of fully resolving chromosome centromeres for accurate transposable element annotation.

In *L. usitatissimum* and *L. bienne*, the *TE_00003234* (*hAT*) DNA transposon family has undergone massive amplification, and now dominates pericentromeric regions, accounting for ∼14% of the entire genome and covering 28–60% of the pericentromeric regions of individual chromosomes. To our knowledge, this represents the first documented example in plants in which a DNA transposon family, rather than LTR retrotransposons, constitutes the major structural component of pericentromeric regions associated with chromosome evolution. This unusual genomic architecture suggests that the lineage-specific expansion of *TE_00003234* may have played a role in remodeling chromosome centers during post-polyploid diploidization, thereby broadening the range of mechanisms through which transposable elements can shape chromosome structure. While the shift toward DNA transposon dominance is evident in both cultivated flax and its wild progenitor *L. bienne*, its role in driving genome evolution requires further study. Nevertheless, this conserved pattern was supported by our analysis of the resequencing of 407 flax accessions, which were compared to the newly generated *L. usitatissimum* reference genome (**Supplemental Figure S10A**).

These findings expand current models of plant chromosome evolution by demonstrating that DNA transposons, in addition to LTR retrotransposons, can dominate pericentromeric architecture and may influence karyotype diversification. More broadly, this work highlights the transformative impact of telomere-to-telomere genome assemblies in uncovering structural events that could not have been described using fragmented genome references.

### Future directions and applications

The near-T2T assemblies and identification of *TE_00003234* as a dominant structural component of chromosome centers open avenues for functional and epigenetic studies of TE-rich regions. Comparative analyses across additional *Linum* species will refine the timing of this TE expansion and clarify its relationship to karyotype change. A detailed structural and evolutionary characterization of this lineage-specific *hAT* family across *Linum* genomes will be addressed in future work.

## CONCLUSIONS

By leveraging telomere-to-telomere genome assemblies, this study uncovers an unexpected mechanistic framework for chromosome evolution in *Linum*. Our analyses showed that extant *Linum* karyotypes originated from an ancestral WGD, followed by lineage-specific polyploidization events and divergent diploidization trajectories. Contrary to classical expectations, major chromosome number increases, particularly the emergence of the *n* = 15 karyotype, occurred without genome expansion and instead involved extensive internal chromosome reorganization.

This reorganization was associated with lineage-specific transposable element activity. A single DNA transposon family (*TE_00003234*, *hAT* superfamily) underwent a concentrated expansion in the *L*. *usitatissimum*/*L*. *bienne* lineage and came to dominate pericentromeric regions, reshaping chromosome architecture, and was associated with recombination and widespread chromosomal fragmentation and reassortment. These findings reveal that DNA transposons, in addition to LTR retrotransposons, may play a role in pericentromeric remodeling and karyotype diversification.

Overall, our results support a hierarchical model of chromosome evolution in which WGD establishes genomic redundancy, where chromosome fusion and fission reshape karyotypes during diploidization, and lineage-specific transposable elements may act as architectural modifiers that stabilize and extend newly formed chromosome configurations. More broadly, this study highlights the power of telomere-to-telomere genome assemblies to uncover links between polyploidy, repetitive element dynamics, and chromosome evolution. From a practical perspective, the CDC Bethune v3.0 genome provides an important resource for flax breeding and genomic selection, particularly given the recombination-suppressed nature of TE-rich pericentromeric domains. This assembly version (v3.0) replaces the previous flax reference genome and provides a complete coordinate framework for future evolutionary, genetic, and breeding studies.

## MATERIALS AND METHODS

### Plant materials

*L. usitatissimum* L. subsp. *usitatissimum* accession CN116251 (cultivar ‘CDC Bethune’) was obtained from Plant Gene Resources Canada (PGRC, Saskatoon, Canada). ‘CDC Bethune’ is an oilseed flax cultivar developed by the University of Saskatchewan, Saskatoon, Canada (Rowland et al. 2002) (**Figure 1A**). *L. bienne* Mill. accession LIN1917, *L. decumbens* Desf. accession LIN1754 and *L. grandiflorum* Desf. accession LIN1530 were obtained from the Leibniz Institute of Plant Genetics and Crop Plant Research (IPK, Gatersleben, Germany). *L. bienne* is a biennial species, and accession LIN1917 is classified as a traditional cultivar/landrace that originated from Spain (DOI:10.25642/IPK/GBIS/97852) (**Figure 1B**). *L. decumbens* is a wild species, and accession LIN1754 originated from Italy (DOI: 10.25642/IPK/GBIS/50485) (**Figure 1C**). *L. grandiflorum* is an annual species, and accession LIN1530 is of unknown geographic origin (DOI:10.25642/IPK/GBIS/35729) (**Figure 1D**).

### Cytogenetic analysis

#### Cell cycle synchronization and mitotic metaphase chromosome preparation

Seeds of the four accessions were soaked in distilled water for 24 h, then placed on moist filter paper for 12 h at 25℃ to germinate. When the roots reached a length of 2-4 mm, they were cut and treated in a nitrous oxide gas chamber (1 MPa) for 2 h. The treated roots were subsequently fixed on ice in cold 90% acetic acid for 5 min, washed in water and stored in 70 % ethanol at −20°C until use.

For chromosome preparation, the fixed roots were washed in distilled water (ddH_2_O) for 5 min. The root tips were cut and placed in 20 μL of an enzyme mixture consisting of 4% cellulase Onozuka R-10 (Yakult, Japan, Tokyo cat. 201069), 1% pectolyase Y23 (Karlan cat. 8006) in KCl buffer (75 mM KCl, 7.5 mM EDTA, pH 4), and incubated at 37°C for 50 min. The digested root tip meristems were washed in TE buffer for 5 min, and twice in 70 % ethanol. The meristems were then dispersed with a needle in 20 μL 9:1 acetic acid–methanol mix and immediately dropped onto precleaned glass slides in a humid chamber. The dried preparations were UV cross-linked and ready for fluorescence *in situ* hybridization (FISH).

#### Fluorescence in situ hybridization (FISH)

For FISH analysis, 45S rDNA labeled with digoxigenin-11-dUTP (Roche 11093088910) was used as a probe. The hybridization mixture (15 μL) contained 50% formamide, 2×SSC, 10% dextran sulfate, salmon sperm DNA, 50 ng 45S rDNA probe and ddH_2_O to make up the final volume. The mixture was denatured by boiling for 13 min, then cooled on ice for at least 10 min. Chromosome preparations were denatured in 0.15 M NaOH, washed twice in 70% ethanol, and once in 100% ethanol. The denatured hybridization mixture was applied to the denatured preparations, covered with a cover glass, and placed in a moist chamber for hybridization at 37°C for 6 h. The cover glass was then removed, and Anti-Dig-Rhodamine (Roche 11207750910) was applied for the antigen-antibody reaction for 2 h. Finally, the preparations were washed in ddH₂O for 10 min and mounted in Vectashield antifade solution (H1200) for microscopy and imaging.

#### Fluorescence microscopy and imaging

The preparations were examined under an OLYMPUS fluorescence microscope (OLYMPUS BX53). Images were captured using a CMOS camera (OLYMPUS DP74) for each filter set. Chromosome counting and size measurements were performed using the OLYMPUS CellSense software. The images were then optimized for contrast and brightness using Adobe Photoshop.

### DNA preparation for sequencing

Seedlings were grown under controlled growth conditions in a growth cabinet providing 20 h light and 4 h darkness with daytime and nighttime temperatures of 24°C and 18°C, respectively. Prior to sampling, i.e., when the first small cluster of true leaves was visible, plants were kept in the dark for 2-3 days. Young leaf tissue was weighed, wrapped in a weigh paper and placed in small envelopes that were immediately frozen in liquid nitrogen where they were kept until DNA extraction.

High molecular weight genomic DNA (HMW gDNA) was extracted from 1.5 g leaf tissue using the NucleoBond HMW DNA Kit (Macherey-Nagel, Allentown, PA, USA), following the manufacturer’s instructions for lysis with liquid nitrogen and mortar and pestle. Final pellets were resuspended in 60 µL low TE (10 mM Tris, 0.1 mM EDTA, pH 8). Following extraction, DNA was kept at 4°C for 2-4 d to ensure its complete resuspension. DNA was quantified using Quant-iT PicoGreen dsDNA assay kit (ThermoFisher Scientific, Waltham, MA, USA) and a preliminary quality control (QC) was performed by pulse-field gel electrophoresis. All DNA samples were then stored at -20°C. A total of 6-9 µg of HMW gDNA per sample was sent to Centre d’expertise et de services Génome Québec (Montreal, QC, Canada) for all four accessions, and to Canadian Grain Commission (Winnipeg, MB, Canada) for CDC Bethune only, for library construction and sequencing.

### Library construction and genome sequencing

HMW gDNA was used to construct SMRTbell libraries following the Preparing Whole Genome and Metagenome Libraries Using SMRTbell® Prep Kit 3.0 protocol (Pacific Biosciences, Menlo Park, CA, USA) or by g-Tubes (Covaris, Woburn, MA, USA). Purified DNA was sheared using a Megaruptor 3 system (Diagenode Inc., Denville, NJ, USA). DNA damage repair, end repair, and SMRTbell adapter ligation were performed according to the same SMRTbell Template Prep Kit 3.0 protocol. Libraries were size-selected using the diluted AMPure PB bead–based protocol (Pacific Biosciences). For CDC Bethune, size selection was also performed using BluePippin using High Pass Plus Cassettes (Sage Science Inc., Beverly, MA, USA). Sequencing primer 3.2 was annealed, and Sequel II Polymerase 2.2 was bound to the SMRTbell templates. Following SMRTbell Cleanup bead purification (Pacific Biosciences) and library concentration estimation using the SMRT Link v11 calculator, libraries were sequenced on a PacBio Sequel II or Sequel IIe platform at a loading concentration of 80 pM using the adaptive loading protocol. Sequencing was performed with the Sequel II Sequencing Kit 2.0 on SMRT Cell 8M, with 30 h movie collections and a 2 h pre-extension.

### Genome assembly

For each accession, PacBio HiFi reads were assembled using hifiasm v0.19.9 (Cheng et al. 2024) with default parameters. To remove organellar contamination, the resulting contigs were aligned to the complete chloroplast genome sequence (NC_002762) from National Center for Biotechnology Information (NCBI) using BLASTN, and chloroplast-derived contigs were removed using a custom Perl script. The remaining filtered nuclear contigs were processed with purge_dups v1.2.6 (https://github.com/dfguan/purge_dups) (Guan et al. 2020) to eliminate duplicated haplotigs and high-coverage artifacts. Contigs were further scaffolded with LRscaf (https://github.com/shingocat/lrscaf) (Qin et al. 2019) using raw PacBio HiFi reads. BioNano optical and genetic maps (see below) were subsequently incorporated to generate super-scaffolds and construct chromosome-scale pseudomolecules. Genome assembly completeness was evaluated using Benchmarking Universal Single-Copy Orthologs (BUSCO) (Simão et al. 2015) in genome mode with the embryophyta_odb10 dataset.

### Telomere identification

In plants, telomeres typically consist of tandem repeats of a short conserved sequence, most commonly TTTAGGG, which are synthesized by telomerase, an RNA-dependent DNA polymerase, and span approximately 2–75 kb at chromosome termini (Shakirov et al. 2022). To assess chromosome completeness and validate scaffold orientation, a custom script was developed to systematically identify telomeric repeat motifs at the ends of each contig or chromosome-scale pseudomolecule.

### BioNano optical maps

*De novo* optical maps of the four accessions were generated using the BioNano IRYS platform. HMW gDNA was isolated from young leaf tissue by Amplicon Express (http://ampliconexpress.com/). DNA molecules were labeled using the nicking endonuclease Nt.*Bsp*QI (New England BioLabs, Ipswich, Mass, USA), followed by fluorescent staining as previously described (Lam et al. 2012). Labeled DNA molecules were loaded onto the NanoChannel arrays of IrysChip devices (BioNano Genomics, San Diego, CA, USA) and automatically imaged using the IRYS system. Across 16 independent runs (168 unique scans), a total of 82 Gb of raw DNA molecules longer than 180 kb (approximately 200× genome coverage) were collected. TIFF images were converted into BNX files using AutoDetect software to extract molecule length and label position information. BNX files were aligned, clustered, and assembled into consensus maps (CMAPs) using the BioNano Assembler pipeline (BioNano Genomics), generating BioNano genome (BNG) maps (Lam et al. 2012; Cao et al. 2014). The P-value thresholds for pairwise assembly, extension/refinement, and final refinement were set to 5 × 10⁻⁸, 5 × 10⁻⁹, and 5 × 10⁻⁹, respectively. Initial BNG maps were examined to identify and remove potential chimeric contigs prior to downstream refinement.

### Genetic maps

To facilitate the construction and validation of chromosome-level pseudomolecules and to investigate structural variations, four high-density genetic linkage maps were developed from three biparental recombinant inbred line (RIL) populations: Bison × Novelty (BN) (Cloutier et al. 2024), Linda × Norman (LN), and LIN1917 × CDC Bethune (LB) (**Supplemental Table S3**). Bison, Novelty, Linda, Norman, and CDC Bethune are elite flax (*L. usitatissimum*) cultivars, while LIN1917 belongs to the progenitor species *L. bienne*. The BN, LN, and LB populations consisted of 702, 158, and 166 RILs, respectively (**Supplemental Table S3**).

All RILs and their respective parental lines were grown in root trainers (72 seedlings per unit) under controlled environmental conditions in a growth chamber (20 h light at 24°C and 4 h dark at 18°C). Genomic DNA was extracted from leaf tissue, and genomic libraries were constructed and sequenced following protocols previously described by Cloutier et al. (2024). Illumina short-read data from each individual were aligned to the flax reference genome CDC Bethune v3.0 (this study) using BWA v0.6.1 (Jo and Koh 2015). Resulting alignments were processed for single nucleotide polymorphism (SNP) discovery using SAMtools v1.12 (Li and Durbin 2009), with all steps integrated into the AGSNP pipeline (You et al. 2012; You et al. 2011) and its subsequent update (Kumar et al. 2012). For the LB population, reads were also mapped to the *L. bienne* (LIN1917) genome assembly to enable SNP identification relative to the progenitor reference genome.

SNPs were filtered to retain high-quality variants with a minor allele frequency (MAF) > 0.05 and a call rate > 80%. Genetic maps were constructed with QTL IciMapping (Meng et al. 2015), which involved the following sequential steps: testing for segregation distortion and removal of SNPs that significantly deviated from the expected Mendelian ratio (1:1), redundancy reduction through binning (P-value > 0.05), linkage group formation based on a recombination frequency threshold of 0.3, and marker ordering using the K-Optimality and 3-OptMap algorithms. In instances where linkage groups contained markers from multiple chromosomes, the “Split into Two” function was applied to resolve and separate the groups into distinct linkage groups corresponding to their respective chromosomes.

### Flax core collection and whole-genome resequencing

The 407 accessions of the flax core collection (You et al. 2017; Diederichsen et al. 2013; Soto-Cerda et al. 2013) were subjected to whole-genome resequencing using the Illumina HiSeq 2000 platform (Illumina Inc., San Diego, USA), generating 100-bp paired-end reads. SNP discovery was performed following previously described procedures (You et al. 2022). Briefly, sequencing reads from each accession were aligned to the CDC Bethune v3.0 reference genome generated in this study using BWA v0.6.1 (Jo and Koh 2015) with a minimum base quality score of Q20 (Phred scale) and default parameters. The resulting alignment files were processed for variant calling using SAMtools (Li and Durbin 2009). Identified variants were further filtered to obtain a high-quality SNP dataset following established criteria (Kumar et al. 2012). All analyses were conducted using the AGSNP pipeline (You et al. 2012; You et al. 2011) and its updated genotyping-by-sequencing (GBS) version (Kumar et al. 2012).

### Genome size estimation

Genome sizes were estimated using flow cytometry and an independent k-mer–based approach. For flow cytometry, young leaf tissue was collected from each *Linum* accession together with tissue from an internal standard and stored in moist paper towels on ice until processing on the same day. Approximately 2 mm² of leaf tissue from each *Linum* sample was co-chopped with the internal standard tissue using a razor blade in 750 µL of cold Galbraith buffer (45 mM MgCl₂, 30 mM sodium citrate, 20 mM MOPS, 0.1% (v/v) Triton X-100, pH 7). The internal standard for *L. usitatissimum*, *L. bienne* and *L. decumbens* was *Camelina sativa* while *Raphanus sativus* was used for *L. grandiflorum*.

Nuclear suspensions were filtered through a 30-µm nylon mesh into glass test tubes, followed by the addition of 50 µL RNase (1 mg mL⁻¹). Samples were stained in the dark for 30 min with 250 µL propidium iodide (100 µg mL⁻¹ in Galbraith buffer). Three biological replicates per accession were analyzed on three separate days using a Gallios flow cytometer (Beckman Coulter, Mississauga, ON, Canada). Flow cytometry data were processed using the R package flowPloidy v3.19 to estimate genome sizes (Smith et al. 2018).

In addition, genome sizes were independently estimated using a long-read-based k-mer approach. K-mer frequencies (*k* = 17) were counted from PacBio reads using kmerfreq (https://github.com/fanagislab/kmerfreq). The resulting frequency profiles were used as input for the GCE (Genomic Character Estimator) pipeline (gce-1.0.2; https://github.com/fanagislab/GCE) to infer genome sizes (Liu et al. 2013).

### Repeat sequence annotation

Repeat annotation was conducted in three major steps. First, transposable elements (TEs) were identified *de novo* for each genome assembly using the Extensive De novo Transposable Element Annotator (EDTA) v2.0.0 pipeline (Ou et al. 2019) with parameters --overwrite 1 --sensitive 1 --anno 1 --evaluate 1. EDTA integrates multiple complementary tools, including LTRharvest (Ellinghaus et al. 2008), LTR_FINDER (Ou and Jiang 2019), LTR_retriever (Ou and Jiang 2018), Generic Repeat Finder (GRF) (Shi and Liang 2019), TIR-Learner (Su et al. 2019), HelitronScanner (Xiong et al. 2014), RepeatModeler (Flynn et al. 2020), and RepeatMasker v4.1.2 (Tarailo-Graovac and Chen 2009), to identify and classify TE families and to generate species-specific, non-redundant TE libraries. Second, these species-specific TE libraries were merged to construct a *Linum* pangenome consensus TE library using the EDTA utility script make_panTElib.pl. The pangenome TE library incorporated nine individual TE libraries, including those from the four assemblies reported herein (*L. usitatissimum* accession CDC Bethune, *L. bienne* accession LIN1917, *L. decumbens* accession LIN1754 and *L. grandiflorum* accession LIN1530), the previously published assemblies of *L. tenue* (Gutierrez-Valencia et al. 2022) and *L. lewisii* (Innes et al. 2023), as well as the assemblies of *L. usitatissimum* cultivars Laura, Bison and Novelty that we generated (NCBI BioProject PRJNA1061544, PRJNA1061532 and PRJNA1061538, respectively). The resulting pangenome TE library was used to annotate the repeats of each genome assembly using RepeatMasker v4.1.2 (Tarailo-Graovac and Chen 2009), ensuring consistent repeat annotation across all assemblies.

Intact long terminal repeat retrotransposons (LTR-RTs) were identified using LTR_FINDER (parallel v1.2; default parameters: -w 2 -C -D 15000 -d 1000 -L 7000 -l 100 -p 20 -M 0.85) and LTRharvest (parallel v1.1; default parameters: -minlenltr 100 -maxlenltr 7000 -mintsd 4 -maxtsd 6 -motif TGCA -motifmis 1 -similar 85), followed by filtering and annotation with LTR_retriever (Ou and Jiang 2018). Only intact LTR-RTs containing identifiable structural protein domains were retained for downstream analyses. Insertion times (T) of intact LTR-RTs were estimated based on sequence divergence between the 5′ and 3′ LTRs. LTR pairs were globally aligned using CLUSTALW (Larkin et al. 2007). Insertion time was calculated as *T = K / (2r)*, where *r* is the neutral substitution rate (1.3 × 10⁻⁸ substitutions per site per year) (Bowen and McDonald 2001; Ma and Bennetzen 2004), and *K* represents the nucleotide divergence between paired LTRs. Divergence (*K*) was estimated using the Kimura two-parameter (K2P) model (Kimura 1980), and standard errors of *K* and *T* were calculated following established methods.

To estimate insertion times for fragmented TEs, TE fragments were aligned to the pangenome consensus sequences of their corresponding TE families using BLASTN. Only fragments meeting the following criteria were retained to reduce estimation bias: length ≥100 bp, coverage of ≥50% of the TE family sequence, and a minimum aligned length of 50 bp. Divergence (*K*) was estimated using the Kimura two-parameter (K2P) model (Kimura 1980). For illustrative purposes, insertion times were calculated as described above. However, because substitution rates may differ between coding and noncoding regions, divergence values (*K*) were also compared directly with *Ks* peaks from whole-genome duplication analyses.

### Protein-coding gene annotation

Structural gene annotation was performed on repeat-masked genome assemblies generated using RepeatMasker v4.1.2 (Tarailo-Graovac and Chen 2009). *Ab initio* gene prediction was conducted using AUGUSTUS v3.4.0 (Stanke et al. 2006) implemented within Braker v3.0.3 (Bruna et al. 2021; Hoff et al. 2016), incorporating extrinsic hints derived from spliced alignments of RNA-seq reads and a curated protein reference dataset. The protein reference dataset was constructed by combining Viridiplantae protein sequences obtained from OrthoDB (odb10) (Kriventseva et al. 2019) with previously published annotated protein sets from *L. usitatissimum* (Wang et al. 2012; Sa et al. 2021; Dvorianinova et al. 2023; Zhao et al. 2023; Arkhipov et al. 2024) and the relative species *L. lewisii* (*n* = 9) (Innes et al. 2023) and *L. tenue* (*n* = 10) (Gutierrez-Valencia et al. 2022). Gene models predicted by the Braker3 pipeline were further curated using a custom post-processing workflow consisting of four steps. First, gene models with incomplete coding sequences (lacking start or stop codons) and short coding sequences (<17 amino acids) were removed. Second, predicted proteins were searched against the TREP database (Wicker et al. 2002) using DIAMOND (Buchfink et al. 2015) with an e-value cutoff of 1e−05 to eliminate repeat-related genes. Third, proteins were queried against the UniProtKB database (UniProt 2023) using DIAMOND with an e-value cutoff of 1e−05, and gene models without significant hits were excluded. Fourth, predicted proteins were aligned to the NCBI non-redundant (*nr*) database to classify genes into high-confidence (HC) and low-confidence (LC) sets. Gene models with alignment coverage ≥80% for both query and subject sequences and an e-value ≤1e−10 were designated as HC; all others were classified as LC. Functional annotations were assigned based on the top-scoring hit. The completeness of the final gene annotation sets was evaluated using Benchmarking Universal Single-Copy Orthologs (BUSCO) v5.2.2 (Simão et al. 2015) in gene mode with the *embryophyta_odb10* dataset. Using DupGen_finder (Qiao et al. 2019), duplicated genes were categorized into five classes: dispersed, proximal, tandem, transposed, and WGD.

Functional annotation of predicted protein-coding genes was performed using eggNOG-mapper v2 (Cantalapiedra et al. 2021) based on the eggNOG v5.0 database (Huerta-Cepas et al. 2019). Protein sequences from each genome were assigned to orthologous groups through sequence similarity searches, allowing functional inference based on evolutionary relationships. Gene Ontology (GO) terms and Kyoto Encyclopedia of Genes and Genomes (KEGG) pathway annotations were obtained directly from eggNOG-mapper outputs.

### Orthogroup analysis and phylogenetic analysis

Orthogroup inference was performed with OrthoFinder v2.5.5 (Emms and Kelly 2019) using protein-coding gene sets from six *Linum* species. Given the extensive gene duplication observed in several *Linum* genomes, loci for phylogenetic reconstruction were identified using a relaxed, gene-tree-based “near-single-copy” criterion. Orthogroups were required to contain at least one gene from each of the six species. Orthogroups exhibiting gene duplication in more than two species were excluded from further analysis. For the remaining orthogroups, a single representative gene was selected per species. When multiple paralogs were present in a species, all combinations of selecting one paralog per duplicated species were enumerated. Each combination was evaluated using the corresponding OrthoFinder gene tree by calculating the sum of pairwise patristic distances among the selected terminal nodes. The combination with the minimum total distance, representing the most compact cross-species clustering consistent with orthology, was retained. In cases of equal scores, ties were resolved by selecting the combination with the greatest total sequence length to minimize potential biases from truncated gene models. Representative sequences from all retained near-single-copy orthogroups were aligned using MAFFT (Katoh and Standley 2013), and individual alignments were concatenated to generate a supermatrix for phylogenetic analysis. Maximum-likelihood phylogenetic inference was performed with RAxML-NG v1.2.2 (Kozlov et al. 2019) under the LG+G+F substitution model with 1,000 bootstrap replicates.

Divergence times were estimated using a relaxed molecular clock as implemented in MCMCTree within the PAML v4.10.9 package (Dos Reis and Yang 2019). The correlated rate model (clock = 3) and the HKY85 substitution model were employed. To account for rate heterogeneity across codon positions, the concatenated supermatrix was partitioned into three independent subsets (first, second, and third codon positions). The MCMC analysis was run for 10,000,000 iterations, sampling every 100 iterations following a burn-in of 3,000,000 iterations. Convergence was assessed by ensuring that all parameter Effective Sample Sizes (ESS) exceeded 200. Temporal calibration was applied to the *Linum* clade node at 38.32 MYA, with a 95% highest probability density (HPD) interval of 36.1-39.5 MYA (Schneider et al. 2016), which is highly consistent with the corresponding estimate reported by Villalvazo-Hernandez et al. (2022).

Synteny and gene collinearity analyses were performed using the SYNY pipeline (Julian and Pombert 2024), which reconstructs conserved gene clusters based on homologous protein pairs identified by DIAMOND. Whole-genome pairwise alignments were additionally generated using minimap2 (Li 2021), and collinearity dot plots were visualized using SYNY or custom scripts.

### Pangenome analysis

Orthogroups inferred across the *Linum* genomes were classified into core and accessory categories following the pangenome framework commonly applied in plant comparative genomics (Golicz et al. 2016). Orthogroups present in all genomes were defined as core, whereas those missing in only one genome were classified as soft-core. Orthogroups detected in several but not all genomes were assigned to the shell category, those present in only a few genomes were designated as rare, and orthogroups restricted to a single genome were defined as private. For each genome, gene copy numbers were summarized within these categories to quantify the relative contributions of conserved and lineage-specific gene sets to genome composition across the *Linum* pangenome.

### Whole genome duplication detection and event time estimation

Whole-genome duplication events and their timeline estimates were inferred for all six *Linum* species using WGD v2.0 (Chen et al. 2024) with default parameters. For each genome, analyses followed a standardized multi-step pipeline provided in WGD 2.0. First, synonymous substitution rate (*Ks*) distributions were generated for the whole-paranome, defined as the complete set of duplicated gene pairs within each genome. Second, intragenomic and intergenomic collinearity relationships were identified to distinguish WGD-derived duplicates from small-scale duplications. Third, mixture models were fitted to the *Ks* distributions to detect statistically significant peaks corresponding to putative WGD events. Fourth, lineage-specific substitution rate variation was corrected by incorporating phylogenetic information from all six *Linum* species, enabling accurate phylogenetic placement of duplication events. Finally, ancient WGDs were dated through phylogenetic dating of WGD-retained duplicate gene pairs under the relaxed molecular clock framework implemented in WGD v2.0. For all species, a minor adjustment to the parameter *-prct* was applied during *Ks* peak identification (modified from the default value of 0.1 to 0.05) to improve resolution of partially overlapping duplication signals. All other parameters were set to the default settings.

### Data availability

All chromosome-scale pseudomolecule assemblies and raw reads generated in this study have been deposited in NCBI under BioProject accessions PRJNA1060737, PRJNA1061532, PRJNA1446840 and PRJNA1446097. In addition, the corresponding genome assemblies, structural and functional gene annotations, repeat annotations, and the flax pangenome repeat library for RepeatMasker generated in this study have been deposited in Zenodo (https://zenodo.org/records/19186083).

## Acknowledgements

This work was conducted as part of the Total Utilization Flax GENomics (TUFGEN) project funded by Genome Canada and other stakeholders, and the A-base project no. 1142 funded by Agriculture and Agri-Food Canada, and the Diverse Field Crop Cluster (DFCC) managed by Ag-West Bio Inc (J-003426). We acknowledge Scott Duguid, Gordon Penner, Elsa Reimer, Andrzej Walichnowski, Natasa Radovanovic, Evelyn Miranda and Helen Booker for their contributions to the development of the mapping populations, Ming-Cheng Luo for assistance with BioNano genome mapping, Tracey James and Sara Martin for assistance with flow cytometry analyses and Madeleine Lévesque-Lemay for photos.

## Author contributions

F.M.Y. and S.C. designed the study; X.W. and J.X. conducted cytogenetic experiments and analyses; S.C., S.W., T.E., and F.M.Y. generated the sequence data; F.M.Y., C.Z., P.L., and L.H. performed genome assembly, annotation, and bioinformatics analyses; F.M.Y. drafted the manuscript; and S.C., S.W., T.E., X.W., L.H., and J.X. revised the manuscript. All authors contributed to manuscript preparation and read, commented on, and approved the final version.

