## Supplemental tables and figures for "Near telomere-to-telomere *Linum* genomes reveal a lineage-specific DNA transposon associated with chromosome architecture remodeling"

**Supplementary tables and figures**

**Supplemental Table S1.** Statistics of PacBio HiFi reads for *L. usitatissimum* cv. CDC Bethune, *L. bienne* accession LIN1917, *L. decumbens* accession LIN1754 and *L. grandiflorum* accession LIN1530

| **Species** | **Accession** | **No. reads** | **Total length (bp)** | **Max length (bp)** | | **Average read length (bp)** | **Coverage* (X)** |
| --- | --- | --- | --- | --- | --- | --- | --- |
| *L. usitatissimum* | CDC Bethune | 2,350,375 | 28,567,596,020 | 66,365 | 12,155 | | 114 |
| *L. bienne* | LIN1917 | 2,066,577 | 20,622,375,659 | 44,462 | 9,979 | | 38 |
| *L. decumbens* | LIN1754 | 8,967,002 | 108,394,116,319 | 62,448 | 12,088 | | 148 |
| *L. grandiflorum* | LIN1530 | 8,683,418 | 106,886,051,426 | 56,197 | 12,309 | | 126 |

* The coverage was estimated based on the K-mer distribution in the assemblies using Hifiasm

**Supplemental Table S2.** BioNano map statistics of four flax genomes

| **Species** | **Accession** | **No. of contigs** | **Total size (Mb)** | **N50 (Mb)** | **Largest contig (Mb)** |
| --- | --- | --- | --- | --- | --- |
| *L. usitatissimum* | CDC Bethune | 251 | 317 | 2.19 | 7.02 |
| *L. bienne* | LIN1917 | 105 | 384 | 9.92 | 14.98 |
| *L. decumbens* | LIN1754 | 946 | 875 | 1.50 | 9.40 |
| *L. grandiflorum* | LIN1530 | 832 | 883 | 1.70 | 17.99 |

**Supplemental Table S3.** Summary of genetic maps developed from three biparental recombinant inbred line (RIL) populations

| **RIL population** | **No. of individuals** | **Reference genome for SNPs** | **No. SNPs^(a)^** | **No. SNPs with sig. seg. dist. (p < 0.05)** | **SNPs with sig. seg. dist. (%)** | **No. of SNPs in genetic map^(b)^** | **No. chr.** | **Total len. of genetic maps (cM)** | **Avg. SNP int. len. (cM)** | **Avg. chr. len. (cM)** | **Genetic distance per chr. (cM)** | **No. SNPs per chr.** |
| --- | --- | --- | --- | --- | --- | --- | --- | --- | --- | --- | --- | --- |
| Bison x Novelty | 703 | CDC Bethune v3.0 | 3,636 | 875 | 24.06 | 1,404 | 15 | 1,506.41 | 1.08 | 100.43 | 93.38-117.24 | 65-160 |
| Linda x Norman | 158 | CDC Bethune v3.0 | 2,010 | 477 | 23.73 | 708 | 15 | 1,447.93 | 2.09 | 96.53 | 61.3-120.27 | 24-75 |
| LIN1917 x CDC Bethune | 166 | CDC Bethune v3.0 | 13,839 | 5,675 | 41.01 | 3,558 | 15 | 3,170.15 | 0.89 | 211.34 | 87.00-312.51 | 105-383 |
| LIN1917 x CDC Bethune | 166 | LIN1917 | 13,756 | 4,873 | 35.42 | 3,604 | 15 | 3,117.03 | 0.87 | 207.8 | 98.52-313.09 | 111-386 |

^(a)^ SNPs after filtering with a call rate >0.8 and minor allele frequency (MAF) >0.05

^(b)^ SNPs after binning and removing markers with significant segregation distortion

**No.**: number; **sig. seg. dist.:** significant segregation distortion; **chr**: chromosome(s); **len.**: length; **int.**: interval; **Avg.**: average.

**Supplemental Table S4.** Pseudochromosomes of *L. usitatissimum* cultivar CDC Bethune, *L. bienne* accession LIN1917, *L. decumbens* accession LIN1754 and *L. grandiflorum* accession LIN1530

| **Chromosome** | **CDC Bethune**  **(*L. usitatissimum*)** | | **LIN1917**  **(*L. bienne*)** | | **LIN1754**  **(*L. decumbens*)** | | **LIN1530**  **(*L. grandiflorum*)** | |
| --- | --- | --- | --- | --- | --- | --- | --- | --- |
|  | **No. of scaffolds** | **Chr. size (bp)** | **No. of scaffolds** | **Chr. size (bp)** | **No. of scaffolds** | **Chr. size (bp)** | **No. of scaffolds** | **Chr. size (bp)** |
| 1 | 2 | 32,352,746 | 1 | 27,549,123 | 1 | 104,652,720 | 12 | 143,123,099 |
| 2 | 1 | 36,797,904 | 1 | 35,061,533 | 3 | 96,753,766 | 11 | 116,634,129 |
| 3 | 1 | 39,172,353 | 1 | 33,073,062 | 2 | 87,375,547 | 10 | 101,080,509 |
| 4 | 1 | 34,927,835 | 2 | 39,175,414 | 1 | 85,721,745 | 16 | 95,880,742 |
| 5 | 2 | 31,717,593 | 2 | 31,650,770 | 1 | 84,243,894 | 6 | 87,355,777 |
| 6 | 1 | 25,011,845 | 1 | 26,737,787 | 2 | 77,338,037 | 12 | 87,008,858 |
| 7 | 3 | 26,806,467 | 2 | 25,439,370 | 2 | 70,879,426 | 21 | 118,658,842 |
| 8 | 1 | 40,448,020 | 1 | 37,125,979 | 2 | 65,085,933 | 6 | 86,841,630 |
| 9 | 3 | 33,717,904 | 2 | 35,439,608 |  |  |  |  |
| 10 | 4 | 36,426,562 | 1 | 30,435,201 |  |  |  |  |
| 11 | 3 | 29,269,339 | 1 | 28,886,650 |  |  |  |  |
| 12 | 2 | 31,253,724 | 2 | 30,871,055 |  |  |  |  |
| 13 | 2 | 27,152,020 | 1 | 30,631,534 |  |  |  |  |
| 14 | 5 | 31,933,221 | 1 | 29,136,146 |  |  |  |  |
| 15 | 2 | 31,567,185 | 2 | 27,113,738 |  |  |  |  |
| Aligned to chromosomes | 33 | 488,554,718 | 21 | 468,326,970 | 14 | 672,051,068 | 96 | 836,658,200 |
| Unaligned to chromosomes | 134 | 15,444,289 | 9 | 703,914 | 348 | 12,995,952 | 299 | 26,799,754 |
| Total | 167 | 503,999,007 | 30 | 469,030,884 | 362 | 685,047,020 | 395 | 863,457,954 |

**chr.**: chromosome

**Supplemental Table S5.** Telomere identification in *L. usitatissimum* (CDC Bethune), *L. bienne* (LIN1917), *L. decumbens* (LIN1754) and *L. grandiflorum* (LIN1530)

| **Species** | **Accession** | **Chromosome** | **Total hits (start)** | **Distance to start (bp)** | **Telomere length at start (bp)** | **Total hits (end)** | **Distance to end (bp)** | **Telomere length at end (bp)** |
| --- | --- | --- | --- | --- | --- | --- | --- | --- |
| *L. usitatissimum* | CDC Bethune | Lu01 | 1,797 | 6 | 12,579 | 2,325 | 38,420 | 16,275 |
|  |  | Lu02 | 1,489 | 119 | 10,423 | 3,520 | 2,284 | 24,640 |
|  |  | Lu03 | 810 | 1,650 | 5,670 | 1,541 | 2,771 | 10,787 |
|  |  | Lu04 | 1,486 | 6 | 10,402 | 1,304 | 155 | 9,128 |
|  |  | Lu05 | 1,396 | 4 | 9,772 | 2,241 | 21,893 | 15,687 |
|  |  | Lu06 | 665 | 107 | 4,655 | 2,324 | 71 | 16,268 |
|  |  | Lu07 | 924 | 1,754 | 6,468 | 2,302 | 2,019 | 16,114 |
|  |  | Lu08 | 603 | 52 | 4,221 | 2,322 | 2,418 | 16,254 |
|  |  | Lu09 | 404 | 110 | 2,828 | 657 | 23,475 | 4,599 |
|  |  | Lu10 | 1,152 | 203,401 | 8,064 | 1,422 | 76 | 9,954 |
|  |  | Lu11 | 2,273 | 508 | 15,911 | 344 | 1,080 | 2,408 |
|  |  | Lu12 | 2,020 | 41 | 14,140 | 1,551 | 517 | 10,857 |
|  |  | Lu13 | 1,471 | 53,790 | 10,297 | 1,169 | 289 | 8,183 |
|  |  | Lu14 | 8,597 | 216,318 | 60,179 | 1,167 | 207 | 8,169 |
|  |  | Lu15 | 2,234 | 18,434 | 15,638 | 1,634 | 482 | 11,438 |
| *L. bienne* | LIN1917 | Lb01 | 180 | 3,155 | 1,260 | 207 | 100 | 1,449 |
|  |  | Lb02 | 232 | 1,466 | 1,624 | 966 | 1,856 | 6,762 |
|  |  | Lb03 | 1,813 | 205 | 12,691 | 1,537 | 1,382 | 10,759 |
|  |  | Lb04 | 1,390 | 32 | 9,730 | 1,570 | 6,345 | 10,990 |
|  |  | Lb05 | 1,322 | 51 | 9,254 | 1,092 | 339 | 7,644 |
|  |  | Lb06 | 1,609 | 83 | 11,263 | 1,254 | 2,013 | 8,778 |
|  |  | Lb07 | 1,207 | 3 | 8,449 | 942 | 42,033 | 6,594 |
|  |  | Lb08 | 839 | 3 | 5,873 | 1,611 | 624 | 11,277 |
|  |  | Lb09 | 2,071 | 3 | 14,497 | 1,767 | 297 | 12,369 |
|  |  | Lb10 | 275 | 241,826 | 1,925 | 371 | 999 | 2,597 |
|  |  | Lb11 | 1,608 | 0 | 11,256 | 634 | 282 | 4,438 |
|  |  | Lb12 | 1,438 | 4 | 10,066 | 795 | 190 | 5,565 |
|  |  | Lb13 | 247 | 64,508 | 1,729 | 13 | 907 | 91 |
|  |  | Lb14 | 966 | 123,644 | 6,762 | 2,017 | 2,466 | 14,119 |
|  |  | Lb15 | 1,131 | 4 | 7,917 | 314 | 34,507 | 2,198 |
| *L. decumbens* | LIN1754 | Ld01 | 447 | 1,419 | 3,129 | 823 | 271 | 6,662 |
|  |  | Ld02 | 512 | 610 | 189 | 880 | 223 | 6,432 |
|  |  | Ld03 | 743 | 180 | 5,495 |  |  |  |
|  |  | Ld04 | 529 | 3,010 | 4,033 | 821 | 3,515 | 6,503 |
|  |  | Ld05 | 226 | 619 | 203 | 1,027 | 7,351 | 7,189 |
|  |  | Ld06 | 819 | 492 | 5,841 | 1,071 | 201 | 7,969 |
|  |  | Ld07 | 931 | 11 | 6,523 | 452 | 589 | 4,295 |
|  |  | Ld08 | 921 | 11 | 6,447 | 285 | 2,001 | 1,995 |
| *L. grandiflorum* | LIN1530 | Lg01 | 98 | 3 | 686 | 427 | 310 | 2,989 |
|  |  | Lg02 | 1,014 | 0 | 7,098 | 870 | 445 | 6,090 |
|  |  | Lg03 |  |  |  |  |  |  |
|  |  | Lg04 | 240 | 1,676 | 1,680 |  |  |  |
|  |  | Lg05 | 528 | 210 | 3,696 | 422 | 474 | 2,954 |
|  |  | Lg06 | 135 | 1,233 | 945 | 1,486 | 236 | 10,402 |
|  |  | Lg07 |  |  |  | 746 | 553 | 5,222 |
|  |  | Lg08 |  |  |  | 1,035 | 70 | 7,245 |

**Supplemental Table S6.** Gene annotation statistics of six *Linum* species

| **Species** | ***L. usitatissimum*** | ***L. bienne*** | ***L. decumbens*** | ***L. grandiflorum*** | ***L. lewisii*** | ***L. tenue*** |
| --- | --- | --- | --- | --- | --- | --- |
| Accession | CDC Bethune | LIN1917 | LIN1754 | LIN1530 | Maple Grove |  |
| Genome size in Mb | 504 | 469 | 685 | 863 | 643 | 692 |
| Total genes | 67,152 | 43,564 | 31,519 | 46,902 | 55,958 | 97,749 |
| Total transcripts | 71,846 | 48,022 | 34,426 | 49,506 | 60,106 | 106,308 |
| Mean gene length (bp) | 1,623 | 2,134 | 1,979 | 1,731 | 1,390 | 1,414 |
| Median gene length (bp) | 1,064 | 1,744 | 1,547 | 1,237 | 836 | 793 |
| Min gene length (bp) | 102 | 102 | 102 | 102 | 102 | 102 |
| Max gene length (bp) | 182,008 | 30,621 | 30,711 | 32,180 | 38,234 | 31,411 |
| Mean exons per gene | 4.04 | 5.25 | 4.52 | 3.99 | 3.4 | 3.35 |
| Median exons per gene | 2 | 4 | 3 | 3 | 2 | 2 |
| Single exon genes | 24195 | 9031 | 8047 | 12479 | 20748 | 38597 |
| Single exon genes (%) | 36.03 | 20.73 | 25.53 | 26.61 | 37.08 | 39.49 |
| Mean exon length (bp) | 232 | 233 | 250 | 249 | 234 | 239 |
| Median exon length (bp) | 142 | 133 | 141 | 153 | 147 | 150 |
| Mean intron length (bp) | 225.42 | 214.43 | 241.28 | 246.34 | 246.33 | 259.74 |
| Median intron length (bp) | 109 | 107 | 109 | 112 | 110 | 121 |
| Genes per Mb | 133.24 | 92.88 | 46.01 | 54.32 | 87.02 | 141.18 |
| Total merged genic region (bp) | 108.80 | 92.85 | 62.30 | 81.04 | 77.51 | 137.61 |
| Total genic region (%) | 21.58 | 19.80 | 9.09 | 9.39 | 12.05 | 19.87 |

**Supplemental Table S7.** Characterization of the largest internal gene-free intervals across chromosomes in six Linum species.

| **Species** | **Name** | **Chromosome** | **Number of genes** | **Max internal gap start (bp)** | **Max internal gap end (bp)** | **Max internal gap length (bp)** | **Left flanking gene id** | **Right flanking gene id** | **Average gap length of species (bp)** |
| --- | --- | --- | --- | --- | --- | --- | --- | --- | --- |
| *L. usitatissimum* | CDC Bethune | Lu1 | 4,890 | 13,904,367 | 20,522,682 | 6,618,316 | CDCBethune01G031500 | CDCBethune01G031510 | 4,497,264 |
|  |  | Lu2 | 3,529 | 10,415,320 | 14,121,308 | 3,705,989 | CDCBethune02G066220 | CDCBethune02G066230 | |
|  |  | Lu3 | 4,486 | 14,165,362 | 22,196,819 | 8,031,458 | CDCBethune03G109990 | CDCBethune03G110000 | |
|  |  | Lu4 | 3,586 | 8,741,606 | 12,822,290 | 4,080,685 | CDCBethune04G138470 | CDCBethune04G138480 | |
|  |  | Lu5 | 3,609 | 15,968,671 | 20,821,137 | 4,852,467 | CDCBethune05G182880 | CDCBethune05G182890 | |
|  |  | Lu6 | 3,497 | 11,448,646 | 17,459,140 | 6,010,495 | CDCBethune06G222330 | CDCBethune06G222340 | |
|  |  | Lu7 | 3,249 | 13,705,382 | 18,267,323 | 4,561,942 | CDCBethune07G258490 | CDCBethune07G258500 | |
|  |  | Lu8 | 4,150 | 24,198,723 | 26,864,469 | 2,665,747 | CDCBethune08G289840 | CDCBethune08G289850 | |
|  |  | Lu9 | 5,028 | 15,875,243 | 20,116,421 | 4,241,179 | CDCBethune09G346900 | CDCBethune09G346910 | |
|  |  | Lu10 | 3,886 | 6,766,939 | 10,427,111 | 3,660,173 | CDCBethune10G374900 | CDCBethune10G374910 | |
|  |  | Lu11 | 4,020 | 13,202,497 | 17,474,394 | 4,271,898 | CDCBethune11G425980 | CDCBethune11G425990 | |
|  |  | Lu12 | 3,755 | 15,926,371 | 20,579,009 | 4,652,639 | CDCBethune12G462370 | CDCBethune12G462380 | |
|  |  | Lu13 | 5,585 | 14,724,702 | 16,177,891 | 1,453,190 | CDCBethune13G519910 | CDCBethune13G519920 | |
|  |  | Lu14 | 4,042 | 9,123,230 | 15,230,685 | 6,107,456 | CDCBethune14G556540 | CDCBethune14G556550 | |
|  |  | Lu15 | 7,964 | 10,928,189 | 13,473,513 | 2,545,325 | CDCBethune15G597320 | CDCBethune15G597330 | |
| *L. bienne* | LIN1917 | Lb1 | 3,882 | 13,608,698 | 16,513,884 | 2,905,187 | LbLIN191701G025920 | LbLIN191701G025930 | 5,446,439 |
|  |  | Lb2 | 2,652 | 12,547,210 | 17,086,963 | 4,539,754 | LbLIN191702G052080 | LbLIN191702G052090 | |
|  |  | Lb3 | 3,333 | 14,333,145 | 18,724,796 | 4,391,652 | LbLIN191703G083840 | LbLIN191703G083850 | |
|  |  | Lb4 | 3,537 | 8,950,606 | 14,926,195 | 5,975,590 | LbLIN191704G109850 | LbLIN191704G109860 | |
|  |  | Lb5 | 2,487 | 10,455,661 | 18,393,305 | 7,937,645 | LbLIN191705G143810 | LbLIN191705G143820 | |
|  |  | Lb6 | 2,758 | 11,452,795 | 19,102,104 | 7,649,310 | LbLIN191706G173450 | LbLIN191706G173460 | |
|  |  | Lb7 | 2,455 | 13,166,440 | 15,118,274 | 1,951,835 | LbLIN191707G202730 | LbLIN191707G202740 | |
|  |  | Lb8 | 3,294 | 8,230,815 | 12,095,952 | 3,865,138 | LbLIN191708G222790 | LbLIN191708G222800 | |
|  |  | Lb9 | 2,751 | 12,583,974 | 18,865,536 | 6,281,563 | LbLIN191709G260130 | LbLIN191709G260140 | |
|  |  | Lb10 | 2,320 | 7,445,171 | 11,022,660 | 3,577,490 | LbLIN191710G280010 | LbLIN191710G280020 | |
|  |  | Lb11 | 2,400 | 16,825,083 | 24,884,622 | 8,059,540 | LbLIN191711G317740 | LbLIN191711G317750 | |
|  |  | Lb12 | 2,903 | 13,623,636 | 19,660,599 | 6,036,964 | LbLIN191712G334590 | LbLIN191712G334600 | |
|  |  | Lb13 | 2,480 | 12,405,716 | 18,895,579 | 6,489,864 | LbLIN191713G360460 | LbLIN191713G360470 | |
|  |  | Lb14 | 3,239 | 8,421,077 | 17,682,567 | 9,261,491 | LbLIN191714G389020 | LbLIN191714G389030 | |
|  |  | Lb15 | 2,799 | 11,078,501 | 13,852,069 | 2,773,569 | LbLIN191715G421090 | LbLIN191715G421100 | |
| *L. decumbens* | LIN1754 | Ld01 | 4,521 | 33,194,964 | 39,724,781 | 6,529,818 | LdLIN175401G016150 | LdLIN175401G016160 | 3,885,332 |
|  |  | Ld02 | 3,908 | 39,483,141 | 42,602,322 | 3,119,182 | LdLIN175402G060570 | LdLIN175402G060580 | |
|  |  | Ld03 | 3,805 | 55,048,803 | 59,554,888 | 4,506,086 | LdLIN175403G108310 | LdLIN175403G108320 | |
|  |  | Ld04 | 3,458 | 32,509,907 | 36,830,313 | 4,320,407 | LdLIN175404G141170 | LdLIN175404G141180 | |
|  |  | Ld05 | 3,708 | 50,680,589 | 53,969,184 | 3,288,596 | LdLIN175405G177940 | LdLIN175405G177950 | |
|  |  | Ld06 | 3,907 | 30,934,119 | 33,194,077 | 2,259,959 | LdLIN175406G217920 | LdLIN175406G217930 | |
|  |  | Ld07 | 3,633 | 23,187,843 | 25,369,637 | 2,181,795 | LdLIN175407G245790 | LdLIN175407G245800 | |
|  |  | Ld08 | 3,542 | 28,169,775 | 33,046,590 | 4,876,816 | LdLIN175408G282040 | LdLIN175408G282050 | |
| *L. grandiflorum* | LIN1530 | Lg01 | 5,082 | 52,765,408 | 58,797,159 | 6,031,752 | LgLIN153001G018310 | LgLIN153001G018320 | 6,908,501 |
|  |  | Lg02 | 4,165 | 46,324,925 | 52,612,025 | 6,287,101 | LgLIN153002G067090 | LgLIN153002G067100 | |
|  |  | Lg03 | 3,741 | 48,942,893 | 55,463,314 | 6,520,422 | LgLIN153003G111390 | LgLIN153003G111400 | |
|  |  | Lg04 | 3,780 | 49,918,798 | 57,758,649 | 7,839,852 | LgLIN153004G152560 | LgLIN153004G152570 | |
|  |  | Lg05 | 3,869 | 44,404,513 | 46,895,878 | 2,491,366 | LgLIN153005G187740 | LgLIN153005G187750 | |
|  |  | Lg06 | 4,201 | 40,804,746 | 44,355,317 | 3,550,572 | LgLIN153006G231350 | LgLIN153006G231360 | |
|  |  | Lg07 | 4,137 | 37,768,817 | 52,202,699 | 14,433,883 | LgLIN153007G263510 | LgLIN153007G263520 | |
|  |  | Lg08 | 4,461 | 41,017,087 | 49,130,149 | 8,113,063 | LgLIN153008G308870 | LgLIN153008G308880 | |
| *L. lewisii* | Maple Grove | Ll1 | 7,401 | 41,902,812 | 42,250,511 | 347,700 | LlMG01G040450 | LlMG01G040460 | 322,528 |
|  |  | Ll2 | 6,438 | 40,001,749 | 40,408,650 | 406,902 | LlMG02G107780 | LlMG02G107790 |  |
|  |  | Ll3 | 5,749 | 36,465,587 | 36,794,916 | 329,330 | LlMG03G171620 | LlMG03G171630 |  |
|  |  | Ll4 | 5,847 | 43,256,938 | 43,606,816 | 349,879 | LlMG04G237710 | LlMG04G237720 |  |
|  |  | Ll5 | 5,462 | 38,713,790 | 38,987,493 | 273,704 | LlMG05G287850 | LlMG05G287860 |  |
|  |  | Ll6 | 5,866 | 34,126,247 | 34,472,849 | 346,603 | LlMG06G347760 | LlMG06G347770 |  |
|  |  | Ll7 | 5,739 | 25,684,075 | 25,963,904 | 279,830 | LlMG07G399120 | LlMG07G399130 |  |
|  |  | Ll8 | 6,020 | 19,489,399 | 19,796,657 | 307,259 | LlMG08G452990 | LlMG08G453000 |  |
|  |  | Ll9 | 4,693 | 16,630,084 | 16,891,632 | 261,549 | LlMG09G510370 | LlMG09G510380 |  |
| *L. tenue* |  | Lt01 | 10,699 | 17,287,309 | 17,389,268 | 101,960 | Lt01G033140 | Lt01G033150 | 164,431 |
|  |  | Lt02 | 8,583 | 20,739,588 | 20,849,105 | 109,518 | Lt02G140550 | Lt02G140560 |  |
|  |  | Lt03 | 12,858 | 67,397,363 | 67,679,212 | 281,850 | Lt03G302220 | Lt03G302230 |  |
|  |  | Lt04 | 10,395 | 1,802,636 | 2,030,949 | 228,314 | Lt04G330010 | Lt04G330020 |  |
|  |  | Lt05 | 9,051 | 56,083,656 | 56,206,737 | 123,082 | Lt05G509540 | Lt05G509550 |  |
|  |  | Lt06 | 12,613 | 13,528,077 | 13,701,939 | 173,863 | Lt06G552090 | Lt06G552100 |  |
|  |  | Lt07 | 10,266 | 356,970 | 486,022 | 129,053 | Lt07G654330 | Lt07G654340 |  |
|  |  | Lt08 | 7,648 | 52,563,214 | 52,795,380 | 232,167 | Lt08G835150 | Lt08G835160 |  |
|  |  | Lt09 | 7,739 | 10,139,856 | 10,293,859 | 154,004 | Lt09G851850 | Lt09G851860 |  |
|  |  | Lt10 | 6,167 | 16,473,550 | 16,584,046 | 110,497 | Lt10G937730 | Lt10G937740 |  |

The maximum internal gap is defined as the longest contiguous region lacking annotated genes between adjacent gene models. Mean gap length is summarized for each species.

**Supplemental Table S8.** Pangenome and gene ontology (GO) analysis of orthogroups and genes for six *Linum* species

| **Species** | **Category** | **Total orthogroups** | **Total gene copies** | **Duplicate gene copies** | **GO annotated copies** | **GO annotated (%)** |
| --- | --- | --- | --- | --- | --- | --- |
| *L. usitatissimum* | Core | 14,218 | 31,705 | 17,487 | 18,140 | 57.22 |
|  | Soft_core | 2,396 | 4,967 | 2,571 | 1,995 | 40.17 |
|  | Shell | 2,039 | 3,511 | 1,472 | 1,112 | 31.67 |
|  | Rare | 4,013 | 4,796 | 783 | 2,050 | 42.74 |
|  | Private | 10,389 | 22,173 | 11,784 | 1,439 | 6.49 |
| *L. bienne* | Core | 14,218 | 31,406 | 17,188 | 18,062 | 57.51 |
|  | Soft_core | 2,350 | 4,559 | 2,209 | 1,956 | 42.90 |
|  | Shell | 1,790 | 2,653 | 863 | 1,014 | 38.22 |
|  | Rare | 3,553 | 3,874 | 321 | 1,977 | 51.03 |
|  | Private | 1,028 | 1,072 | 44 | 33 | 3.08 |
| *L. grandiflorum* | Core | 14,218 | 21,114 | 6,896 | 11,431 | 54.14 |
|  | Soft_core | 2,308 | 4,233 | 1,925 | 1,431 | 33.81 |
|  | Shell | 1,588 | 3,694 | 2,106 | 1,019 | 27.59 |
|  | Rare | 1,443 | 2,178 | 735 | 879 | 40.36 |
|  | Private | 12,747 | 15,683 | 2,936 | 490 | 3.12 |
| *L. decumbens* | Core | 14,218 | 19,010 | 4,792 | 10,493 | 55.20 |
|  | Soft_core | 2,320 | 3,366 | 1,046 | 1,252 | 37.20 |
|  | Shell | 1,441 | 4,372 | 2,931 | 897 | 20.52 |
|  | Rare | 1,261 | 2,120 | 859 | 896 | 42.26 |
|  | Private | 2,013 | 2,651 | 638 | 310 | 11.69 |
| *L. lewisii* | Core | 14,218 | 20,490 | 6,272 | 11,166 | 54.49 |
|  | Soft_core | 1,687 | 3,356 | 1,669 | 1,005 | 29.95 |
|  | Shell | 1,317 | 3,055 | 1,738 | 734 | 24.03 |
|  | Rare | 1,140 | 3,889 | 2,749 | 645 | 16.59 |
|  | Private | 12,228 | 25,168 | 12,940 | 2,093 | 8.32 |
| *L. tenue* | Core | 14,218 | 32,289 | 18,071 | 17,791 | 55.10 |
|  | Soft_core | 1,029 | 2,887 | 1,858 | 942 | 32.63 |
|  | Shell | 1,181 | 3,445 | 2,264 | 772 | 22.41 |
|  | Rare | 926 | 3,639 | 2,713 | 691 | 18.99 |
|  | Private | 27,173 | 55,489 | 28,316 | 4,469 | 8.05 |

**Supplemental Table S9**. Number of orthogroups with single-copy and duplicated genes in six *Linum* species

| **Species** | **Accession** | **Number of chromosomes (2*n*)** | **Single-copy** | | **Duplicated** | | **Total** |
| --- | --- | --- | --- | --- | --- | --- | --- |
|  |  |  | **No.** | **(%)** | **No.** | **(%)** |  |
| *L. tenue* | - | 10 | 5,883 | 21.55 | 21,415 | 78.45 | 27,298 |
| *L. lewisii* | Maple Grove | 9 | 13,548 | 63.98 | 7,626 | 36.02 | 21,174 |
| *L. decumbens* | LIN1754 | 8 | 15,387 | 79.10 | 4,065 | 20.90 | 19,452 |
| *L. grandiflorum* | LIN1530 | 8 | 14,320 | 71.47 | 5,715 | 28.53 | 20,035 |
| *L. bienne* | LIN1917 | 15 | 7,121 | 32.45 | 14,823 | 67.55 | 21,944 |
| *L. usitatissimum* | CDC Bethune | 15 | 7,327 | 30.81 | 16,458 | 69.19 | 23,785 |
| Total |  |  |  |  |  |  | 40,115 |

**Supplemental Table Table S10.** Number of single-copy and duplicated genes identified in the Benchmarking Universal Single-Copy Orthologs (BUSCO) analysis from the genome assemblies of six *Linum* species. The *embryophyte_odb10* dataset, comprising 1,614 conserved orthologs, was used

| **Species** | **Accession** | **Number of chromosomes (2*n*)** | **Complete single-copy** | | **Complete duplicated** | | **Total complete genes** |
| --- | --- | --- | --- | --- | --- | --- | --- |
|  |  |  | **No.** | **%** | **No.** | **%** |  |
| *L. tenue* | - | 10 | 346 | 30.0 | 808 | 70.0 | 1,154 |
| *L. lewisii* | Maple Grove | 9 | 1,253 | 85.9 | 206 | 14.1 | 1,459 |
| *L. decumbens* | LIN1754 | 8 | 1,355 | 86.3 | 216 | 13.7 | 1,571 |
| *L. grandiflorum* | LIN1530 | 8 | 1,248 | 79.3 | 325 | 20.7 | 1,573 |
| *L. bienne* | LIN1917 | 15 | 361 | 23.0 | 1,211 | 77.0 | 1,572 |
| *L. usitatissimum* | CDC Bethune | 15 | 394 | 24.9 | 1,188 | 75.1 | 1,582 |

**Supplemental Table S11.** Whole-genome duplication (WGD) events (MYA) estimated for six *Linum* species.

| **Species** | **Accession/Cultivar** | **Ancient WGD event (MYA)** | **Most recent WGD event (MYA)** |
| --- | --- | --- | --- |
| *L. tenue* |  | 56.16 | 26.32 |
| *L. lewisii* | Maple Grove | 48.00 | – |
| *L. grandiflorum* | LIN1530 | 53.86 | (3.14) ^†^ |
| *L. decumbens* | LIN1754 | 47.85 | – |
| *L. bienne* | LIN1917 | 51.03 | 7.58 |
| *L. usitatissimum* | CDC Bethune | 51.93 | 6.63 |

^†^ A minor Ks peak was detected in *L. grandiflorum* at ~3.14 MYA; however, this signal is more likely attributable to assembly artifacts associated with genome heterozygosity rather than a genuine WGD event.

**Supplemental Table S12.** Repeat sequence analysis of genome assemblies of six *Linum* species

| **Class** | **Subclass** | **Superfamily** | ***L. tenue***  **(n=10)** | | | ***L. lewisii***  **(Maple Grove)**  **(n=9)** | | | ***L. decumbens***  **(LIN1754)**  **(n=8)** | | | | ***L. grandiflorum***  **(LIN1530)**  **(n=8)** | | | | ***L. bienne***  **(LIN1917)**  **(n=15)** | | | | ***L. usitatissimum***  **(CDC Bethune v3.0)**  **(n=15)** | | |
| --- | --- | --- | --- | --- | --- | --- | --- | --- | --- | --- | --- | --- | --- | --- | --- | --- | --- | --- | --- | --- | --- | --- | --- |
|  |  |  | **No. of hits** | **Length (Mb)** | **%** | **No. of hits** | **Length (Mb)** | **%** | **No. of hits** | **Length (Mb)** | **%** | **No. of hits** | | **Length (Mb)** | **%** | **No. of hits** | | **Length (Mb)** | **%** | **No. of hits** | | **Length (Mb)** | **%** |
| **Class I: Retrotransposons** | | | **647,928** | **300.792** | **43.44** | **559,309** | **343.863** | **53.48** | **588,813** | **382.448** | **55.83** | **816,783** | | **461.432** | **53.44** | **193,742** | | **63.369** | **13.5** | **242,931** | | **71.690** | **14.22** |
|  | LTR | Copia | 167,878 | 77.635 | 11.21 | 64,424 | 30.279 | 4.71 | 48,761 | 35.412 | 5.17 | 57,352 | | 37.396 | 4.33 | 25,223 | | 17.045 | 3.63 | 26,106 | | 17.995 | 3.57 |
|  |  | Gypsy | 222,666 | 135.065 | 19.51 | 235,869 | 179.674 | 27.94 | 317,067 | 237.322 | 34.64 | 455,821 | | 279.336 | 32.35 | 65,118 | | 19.945 | 4.25 | 102,739 | | 26.112 | 5.18 |
|  |  | Unknown | 244,685 | 82.463 | 11.91 | 244,854 | 129.299 | 20.11 | 207,228 | 102.959 | 15.03 | 286,517 | | 137.256 | 15.9 | 92,127 | | 22.091 | 4.71 | 102,241 | | 23.006 | 4.56 |
|  | nonLTR | LINE | 12,699 | 5.629 | 0.81 | 14,162 | 4.611 | 0.72 | 15,757 | 6.756 | 0.99 | 17,093 | | 7.444 | 0.86 | 11,274 | | 4.288 | 0.91 | 11,845 | | 4.576 | 0.91 |
| **Class II: DNA transposons** | | | **277,382** | **85.128** | **12.29** | **500,363** | **129.836** | **20.18** | **440,667** | **145.763** | **21.28** | **572,095** | | **224.242** | **25.97** | **405,531** | | **164.606** | **35.10** | **444,155** | | **182.098** | **36.12** |
|  | TIR | CACTA | 14,310 | 2.026 | 0.29 | 98,508 | 26.670 | 4.15 | 84,564 | 20.114 | 2.94 | 102,762 | | 21.976 | 2.55 | 64,343 | | 18.035 | 3.85 | 75,523 | | 23.807 | 4.72 |
|  |  | Mutator | 26,190 | 5.591 | 0.81 | 193,391 | 50.880 | 7.91 | 202,266 | 95.438 | 13.93 | 303,017 | | 170.238 | 19.72 | 154,613 | | 42.356 | 9.03 | 172,995 | | 48.836 | 9.69 |
|  |  | PIF_Harbinger | 3,572 | 0.709 | 0.1 | 15,745 | 3.735 | 0.58 | 18,543 | 4.037 | 0.59 | 21,023 | | 4.264 | 0.49 | 27,980 | | 7.185 | 1.53 | 31,133 | | 12.672 | 2.51 |
|  |  | Tc1_Mariner | 1,492 | 0.150 | 0.02 | 8,379 | 1.603 | 0.25 | 8,647 | 2.228 | 0.33 | 9,014 | | 2.194 | 0.25 | 17,468 | | 2.092 | 0.45 | 16,104 | | 2.162 | 0.43 |
|  |  | hAT | 15,514 | 2.882 | 0.42 | 93,084 | 20.794 | 3.23 | 48,902 | 10.404 | 1.52 | 53,659 | | 11.082 | 1.28 | 42,206 | | 74.422 | 15.87 | 42,921 | | 73.788 | 14.64 |
|  | nonTIR | Helitron | 215,023 | 73.611 | 10.63 | 90,573 | 26.019 | 4.05 | 77,522 | 13.524 | 1.97 | 82,371 | | 14.463 | 1.68 | 98,658 | | 20.496 | 4.37 | 105,209 | | 20.810 | 4.13 |
|  | Others |  | 1,281 | 0.159 | 0.02 | 683 | 0.134 | 0.01 | 223 | 0.019 | 0 | 249 | | 0.025 | 0 | 263 | | 0.019 | 0 | 270 | | 0.021 | 0 |
| **Unknown** |  |  | **179,615** | **50.506** | **7.29** | **109,473** | **28.689** | **4.46** | **130,298** | **34.647** | **5.06** | **156,201** | | **42.567** | **4.93** | **336,281** | | **69.674** | **14.85** | **374,189** | | **78.685** | **15.61** |
| **Total** |  |  | **1,104,925** | **436.427** | **63.03** | **1,169,145** | **502.387** | **78.13** | **1,159,778** | **562.858** | **82.16** | **1,545,079** | | **728.241** | **84.35** | **935,554** | | **297.648** | **63.46** | **1,061,275** | | **332.472** | **65.97** |

**Supplemental Table S13.** Chromosome-wide enrichment of *TE_00003234* in low-SNP/low-gene-density regions of the flax CDC Bethune genome (v3.0).

| **Chr** | **Chr size (Mb)** | **LowRegion size (Mb)** | **NonLowRegion size (Mb)** | **TE_00003234 total size (Mb)** | **TE_00003234 in LowRegion (Mb)** | **TE_00003234 in NonLowRegion (Mb)** | **TE_00003234 in Chr (%)** | **TE_00003234 of LowRegion (%)** | **TE_00003234 of NonLowRegion (%)** | **Enrichment fold for Low vs NonLow** | **Fisher odds ratio** | **Fisher *P* value** |
| --- | --- | --- | --- | --- | --- | --- | --- | --- | --- | --- | --- | --- |
| Lu1 | 32.35 | 10.90 | 21.45 | 2.43 | 2.07 | 0.36 | 7.50 | 18.96 | 1.68 | 11.26 | 13.66 | 0.00 |
| Lu2 | 36.80 | 20.40 | 16.40 | 5.74 | 5.28 | 0.46 | 15.60 | 25.89 | 2.81 | 9.21 | 12.07 | 0.00 |
| Lu3 | 39.17 | 19.90 | 19.27 | 8.27 | 7.86 | 0.41 | 21.10 | 39.49 | 2.11 | 18.74 | 30.31 | 0.00 |
| Lu4 | 34.93 | 20.70 | 14.23 | 4.32 | 4.11 | 0.20 | 12.36 | 19.86 | 1.44 | 13.79 | 16.96 | 0.00 |
| Lu5 | 31.72 | 14.90 | 16.82 | 0.49 | 0.07 | 0.42 | 1.55 | 0.48 | 2.50 | 0.19 | 0.19 | 0.00 |
| Lu6 | 25.01 | 7.00 | 18.01 | 6.75 | 6.04 | 0.71 | 27.00 | 86.29 | 3.95 | 21.83 | 152.92 | 0.00 |
| Lu7 | 26.81 | 12.00 | 14.81 | 5.08 | 4.52 | 0.55 | 18.94 | 37.69 | 3.74 | 10.07 | 15.56 | 0.00 |
| Lu8 | 40.45 | 24.00 | 16.45 | 0.38 | 0.00 | 0.38 | 0.93 | 0.00 | 2.30 | 0.00 | 0.00 | 0.00 |
| Lu9 | 33.72 | 16.00 | 17.72 | 5.09 | 4.46 | 0.63 | 15.11 | 27.88 | 3.58 | 7.80 | 10.42 | 0.00 |
| Lu10 | 36.43 | 15.50 | 20.93 | 7.57 | 6.73 | 0.85 | 20.78 | 43.40 | 4.04 | 10.75 | 18.22 | 0.00 |
| Lu11 | 29.27 | 15.80 | 13.47 | 2.99 | 2.87 | 0.13 | 10.22 | 18.13 | 0.94 | 19.24 | 23.28 | 0.00 |
| Lu12 | 31.25 | 13.50 | 17.75 | 6.89 | 6.62 | 0.27 | 22.04 | 49.03 | 1.50 | 32.63 | 63.07 | 0.00 |
| Lu13 | 27.15 | 7.80 | 19.35 | 2.77 | 2.41 | 0.36 | 10.19 | 30.90 | 1.84 | 16.75 | 23.80 | 0.00 |
| Lu14 | 31.93 | 11.90 | 20.03 | 6.42 | 4.49 | 1.93 | 20.12 | 37.74 | 9.65 | 3.91 | 5.68 | 0.00 |
| Lu15 | 31.57 | 15.20 | 16.37 | 1.83 | 0.00 | 1.83 | 5.80 | 0.00 | 11.19 | 0.00 | 0.00 | 0.00 |
| Whole genome | 488.56 | 225.50 | 263.06 | 67.02 | 57.53 | 9.49 | 13.72 | 25.51 | 3.61 | 7.07 | 9.15 | 0.00 |

Lu: chromosome; LowRegion/Low: low-SNP/low-gene-density region; NonLowRegion/NonLow: non-low-SNP/low-gene-density region.

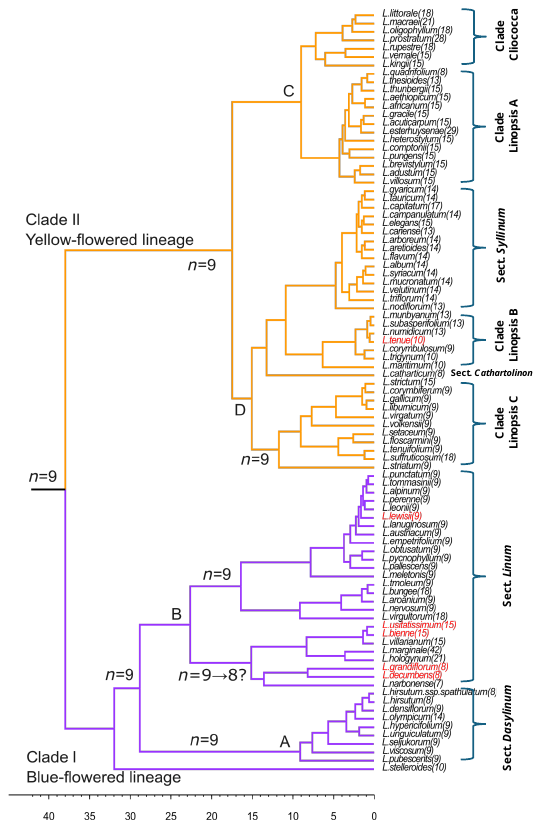
**Supplementary Figures**

**Supplemental Figure S1**. **Phylogenetic tree of the subfamily Linoideae based on S-DIVA analysis, adapted from Villalvazo-Hernandez et al. (2022**). Chromosome numbers for each species are provided in parentheses. Species analyzed in the present study are highlighted with orange boxes. The ancestral chromosome number of the *Linum* s.l. clade is visually inferred to be *n* = 9, based on the chromosome counts of species across the various subclades.

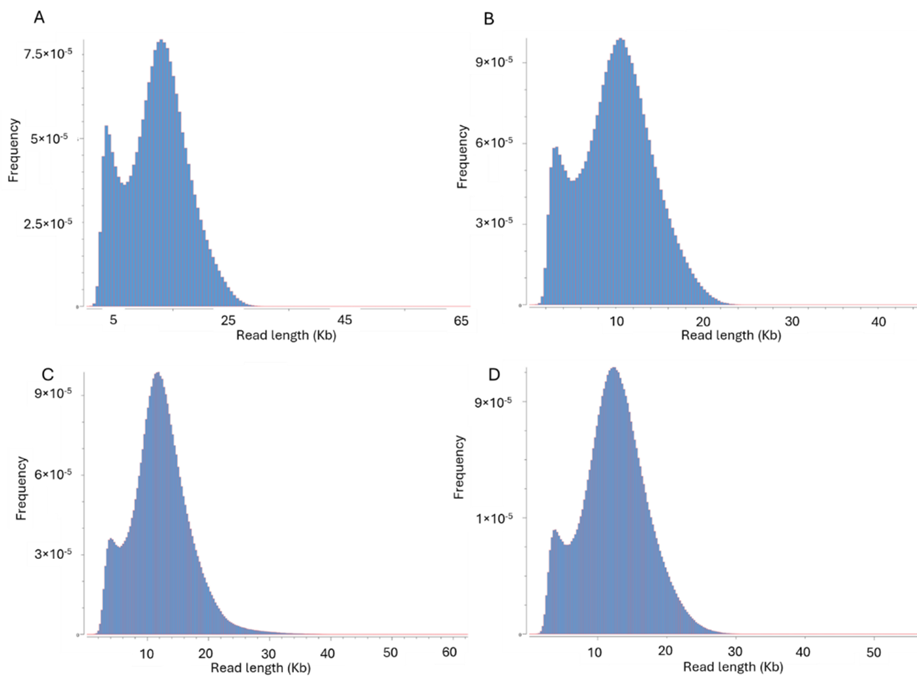

**Supplemental Figure S2.** **Read length distribution of PacBio HiFi CCS reads**. (**A**) L. usitatissimum cultivar CDC Bethune, (**B**) L. bienne accession LIN1917, (**C**) L. decumbens accession LIN1754, and (**D**) L. grandiflorum accession LIN1530.

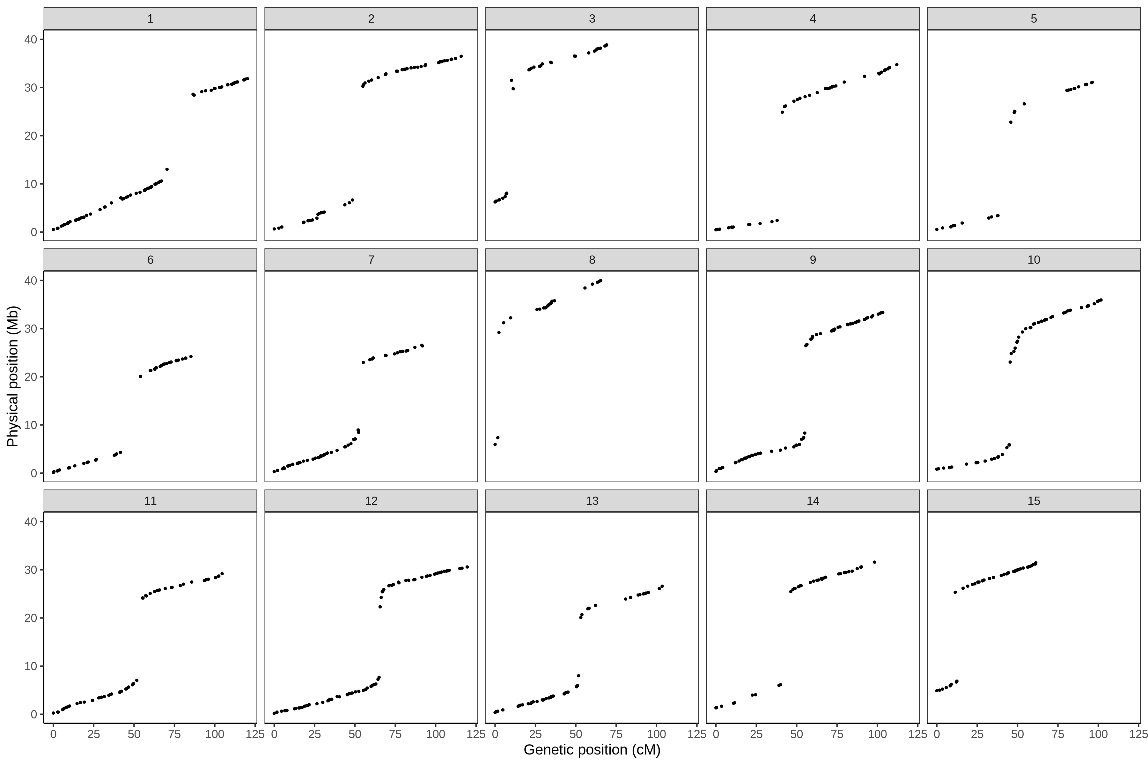

**Supplemental Figure S3. Colinear relationship between the genetic map (in centimorgans, cM) on the x-axis and the physical map (in base pairs, bp) on the y-axis of the Linda × Norman population (Supplemental Table S3)**. The physical positions are based on the *CDC Bethune v3.0* reference genome.

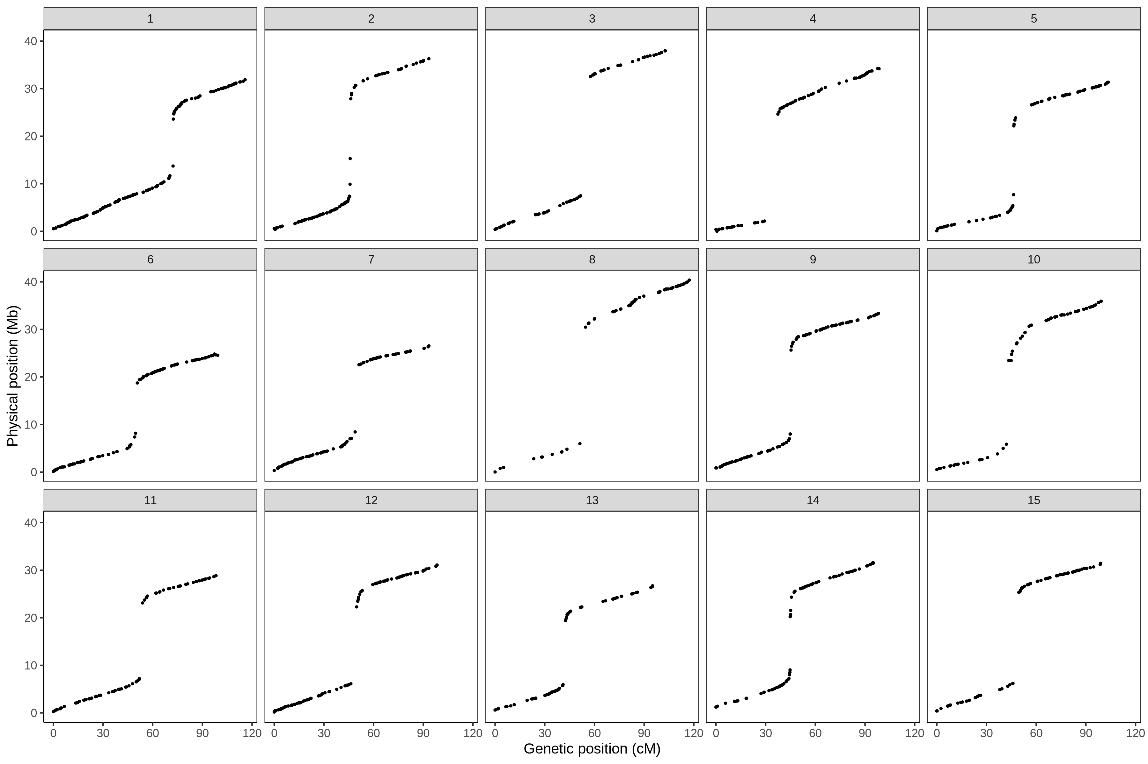

**Supplemental Figure S4.** **Colinear relationship between the genetic map (in centimorgans, cM) on the x-axis and the physical map (in base pairs, bp) on the y-axis of the Bison × Novelty population (Supplemental Table S3)**. The physical positions are based on the *CDC Bethune v3.0* reference genome.

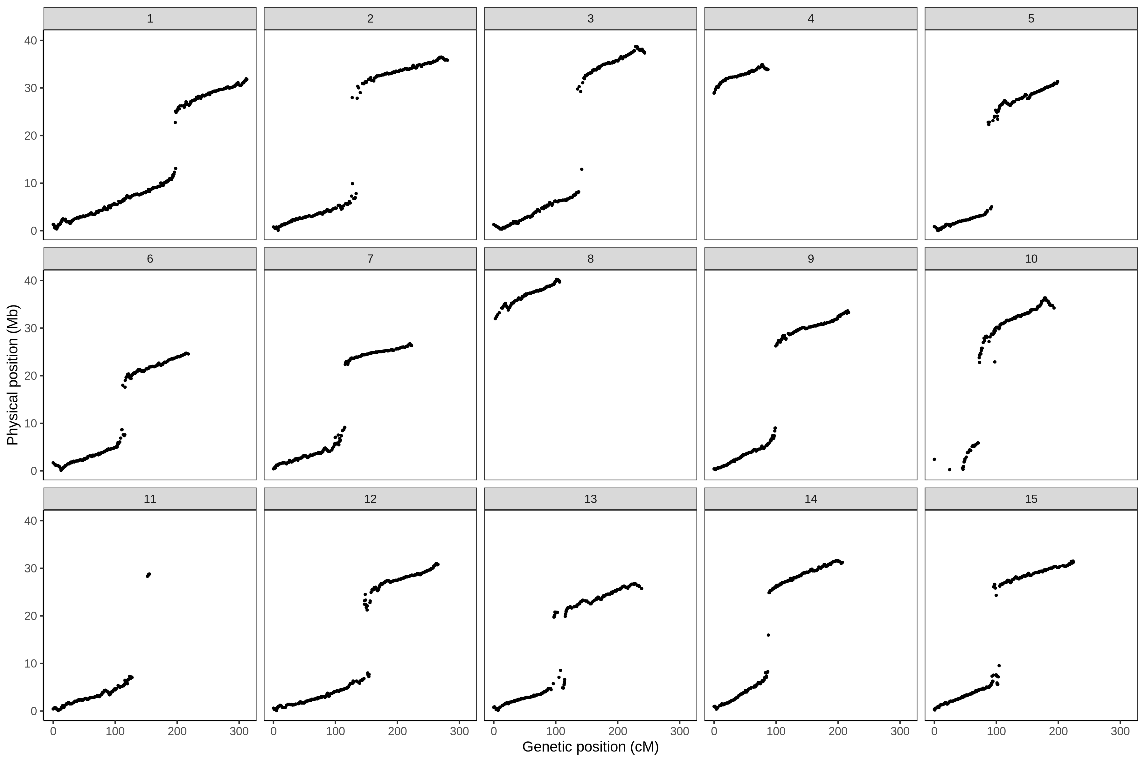

**Supplemental Figure S5. Colinear relationship between the genetic map (in centimorgans, cM) on the x-axis and the physical map (in base pairs, bp) on the y-axis of the LIN1917 (*L. bienne*) × CDC Bethune (*L. usitatissimum*) population (Supplemental Table S3).** The physical positions are based on the *CDC Bethune v3.0* reference genome.

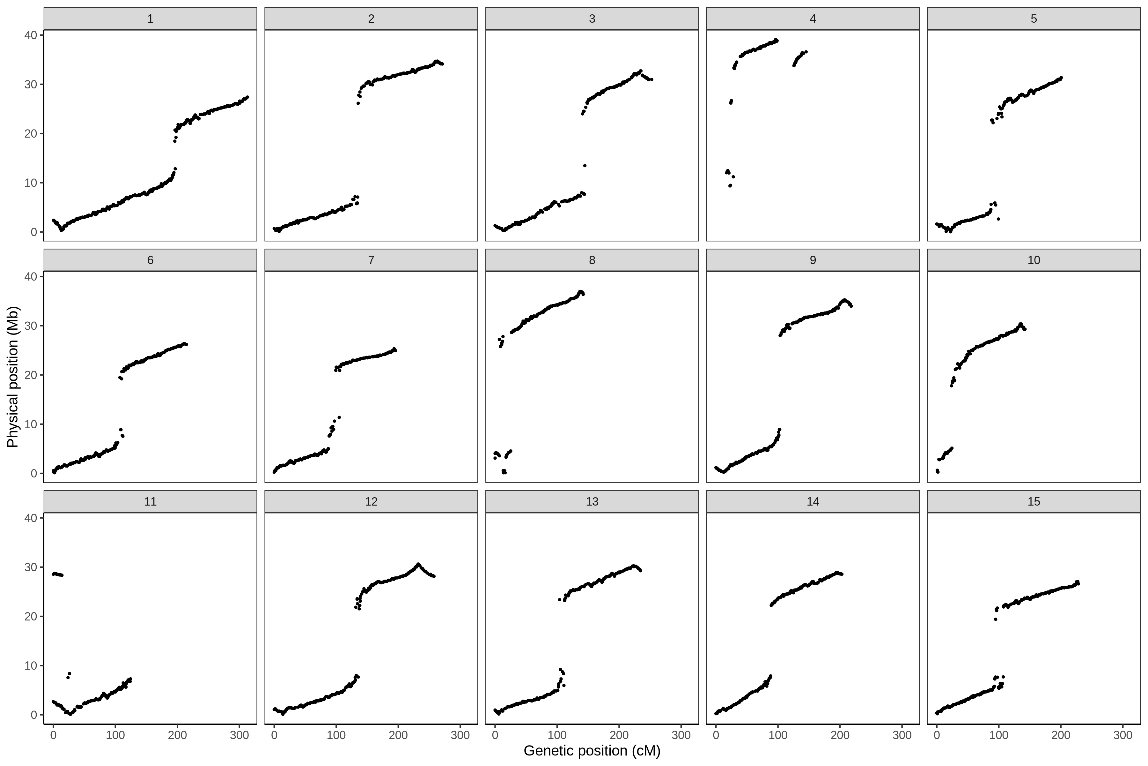

**Supplemental Figure S6. Colinear relationship between the genetic map (in centimorgans, cM) on the x-axis and the physical map (in base pairs, bp) on the y-axis of the LIN1917 (*L. bienne*) × CDC Bethune (*L. usitatissimum*) population (Supplemental Table S3).** The physical positions are based on the LIN1917 reference genome.

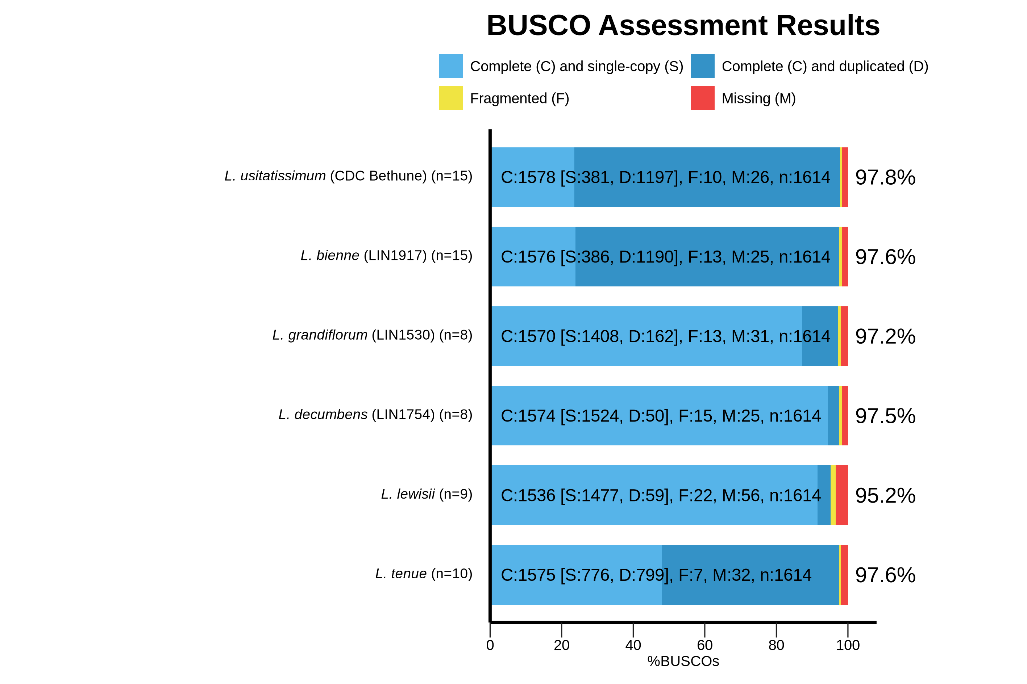

**Supplemental Figure S7.** **Benchmarking Universal Single-Copy Orthologs (BUSCO) analysis results of the genome assemblies of four *Linum* species and comparison with the previously reported assemblies of *L. tenue* (Gutierrez-Valencia et al. 2022) and *L. lewisii* (Innes et al. 2023)**. The embryophyte_odb10 dataset, comprising 1,614 conserved orthologs, was used.

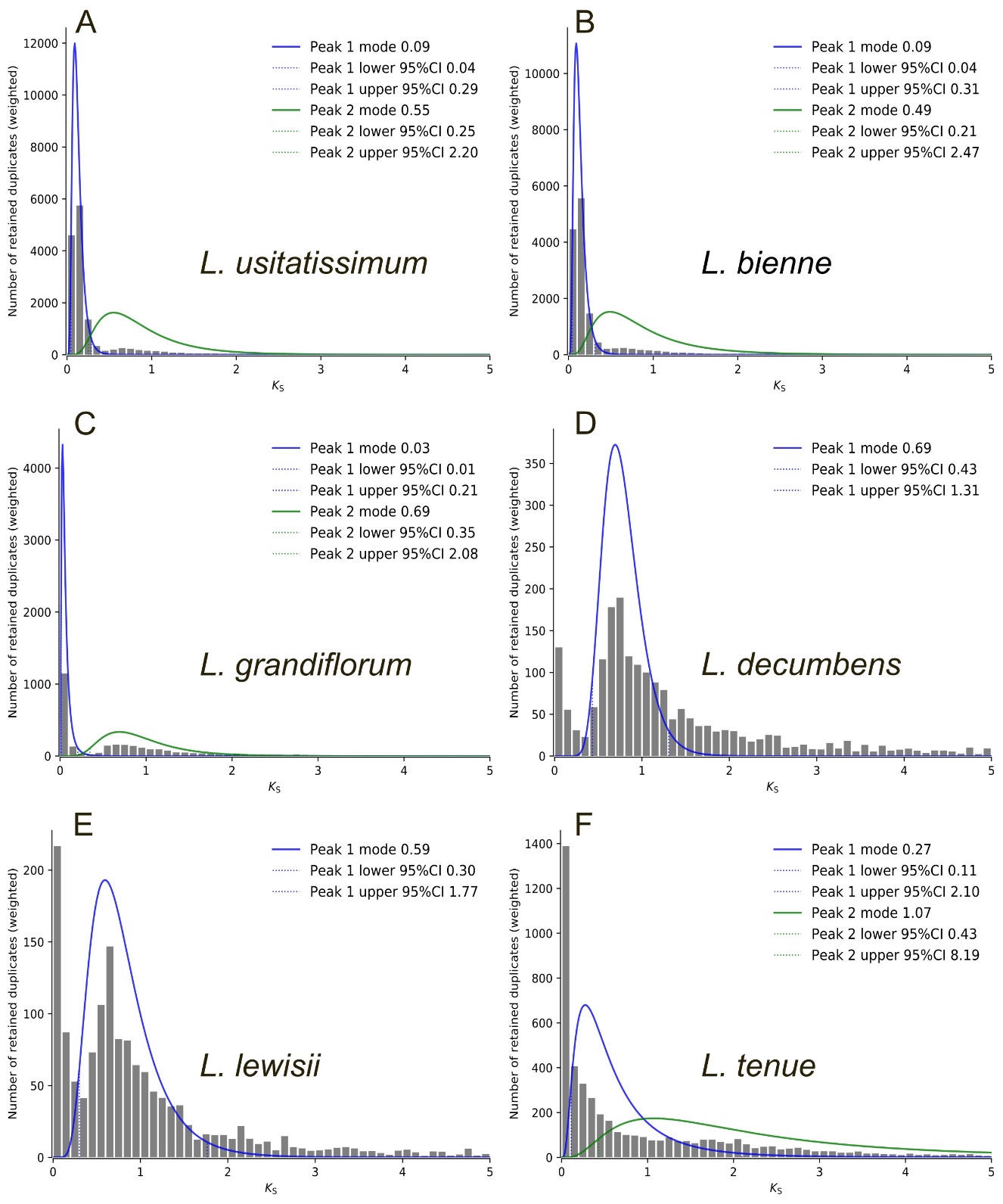

**Supplemental Figure S8. *Ks* peaks identified using WGD v2.0 based on the weighted *Ks* distributions of duplicated genes in six *Linum* species**: (**A**) *L. usitatissimum*, (**B**) *L. bienne*, (**C**) *L. grandiflorum*, (**D**) *L. decumbens*, (**E**) *L. lewisii*, and (**F**) *L. tenue*.

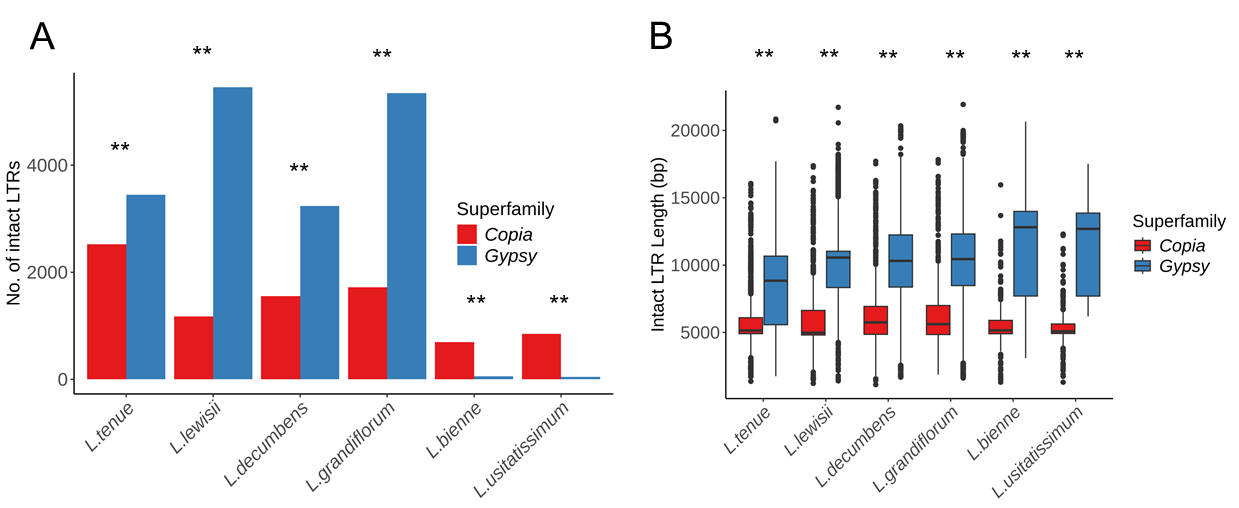

**Supplemental Figure S9. Comparison of the features of intact long terminal repeat (LTR) transposable elements across six *Linum* species**. (**A**) Number of intact LTR elements, and (**B**) length of intact LTR elements~~.~~ Letters above the boxes in (**A**) and (**B**) indicate significant differences among species. In (**B**), ** indicates significant differences between *Copia* and *Gypsy* at the 1% probability level and NS indicates no significant difference.

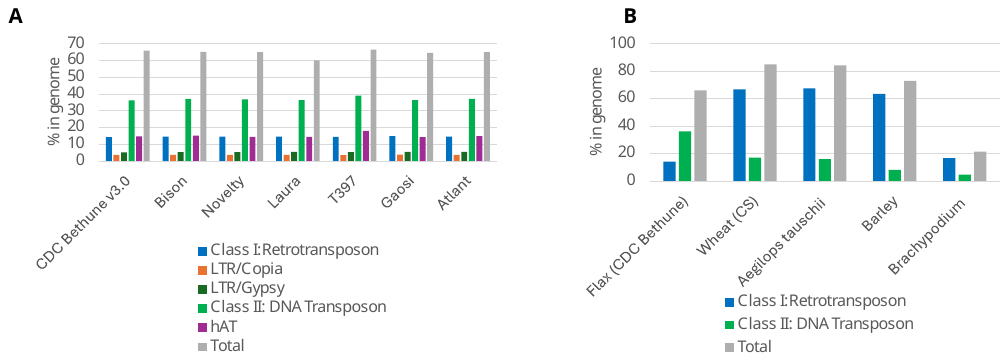

**Supplemental Figure S10. Transposable element (TE) composition across flax genomes and comparison with representative plant species**. (**A**) Proportions of major TE classes in eight flax genomes, including CDC Bethune v3.0, Bison, Novelty, Laura, T397, Gaosi, and Atlant. TE categories include Class I retrotransposons (LTR/*Copia* and LTR/*Gypsy*), Class II DNA transposons, and total TE content. (**B**) Comparison of TE composition between flax (*CDC Bethune v3.0*) and other plant genomes, including wheat (CS) (Zhu et al. 2021), *Aegilops tauschii* (Luo et al. 2017), barley (Mascher et al. 2017), and *Brachypodium* (International Brachypodium 2010). Values represent the percentage of each genome occupied by TE classes and total TE content. The same pipeline for TE identification was used for the flax cultivars, Bison (NCBI BioProject PRJNA1061532), Novelty (PRJNA1061538), Laura (PRJNA1061544), T397 (Yadav et al. 2025), Gaosi (Lu et al. 2025), and Atlant (Dmitriev et al. 2020). Data for the other plant species were obtained from previously published studies.

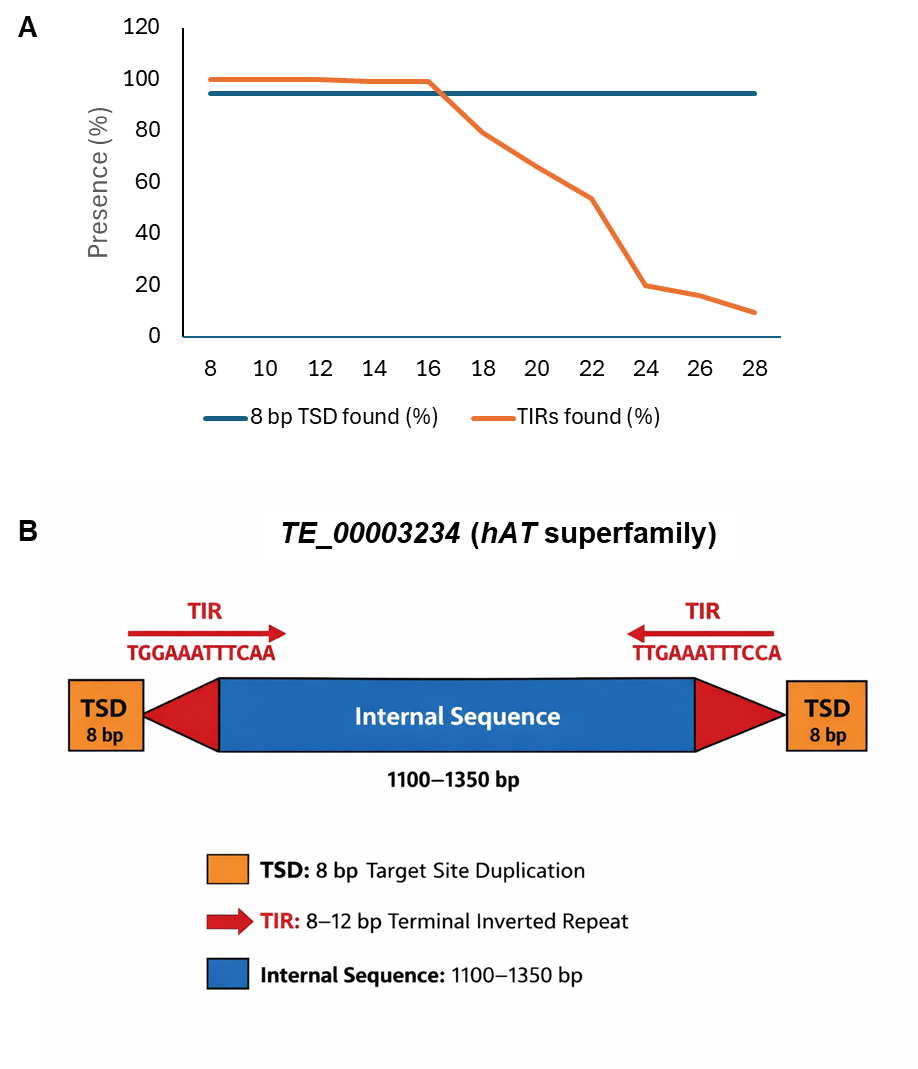

**Supplemental Figure S11. Structural model and terminal feature analysis of the *TE_00003234* DNA transposon in flax.** (A) Evaluation of target site duplications (TSDs) and terminal inverted repeats (TIRs) of *TE_00003234*, a DNA transposon belonging to the hAT superfamily. A total of 144 complete copies were identified in the *L. usitatissimum* CDC Bethune genome. The presence of 8 bp TSDs and TIRs was examined across a range of candidate TIR lengths. TSD detection parameters were window size = 80 bp, maximum mismatches = 0, and positional shift = 5 bp. TIR detection parameters were window size = 120 bp, maximum mismatches = 4 bp, and minimum sequence identity = 75%. (B) Structural model of *TE_00003234*. The element is flanked by 8 bp TSDs and 8-12 bp TIRs. The internal sequence spans approximately 1.1–1.35 kb overall representing the typical structural organization of hAT-superfamily DNA transposons.

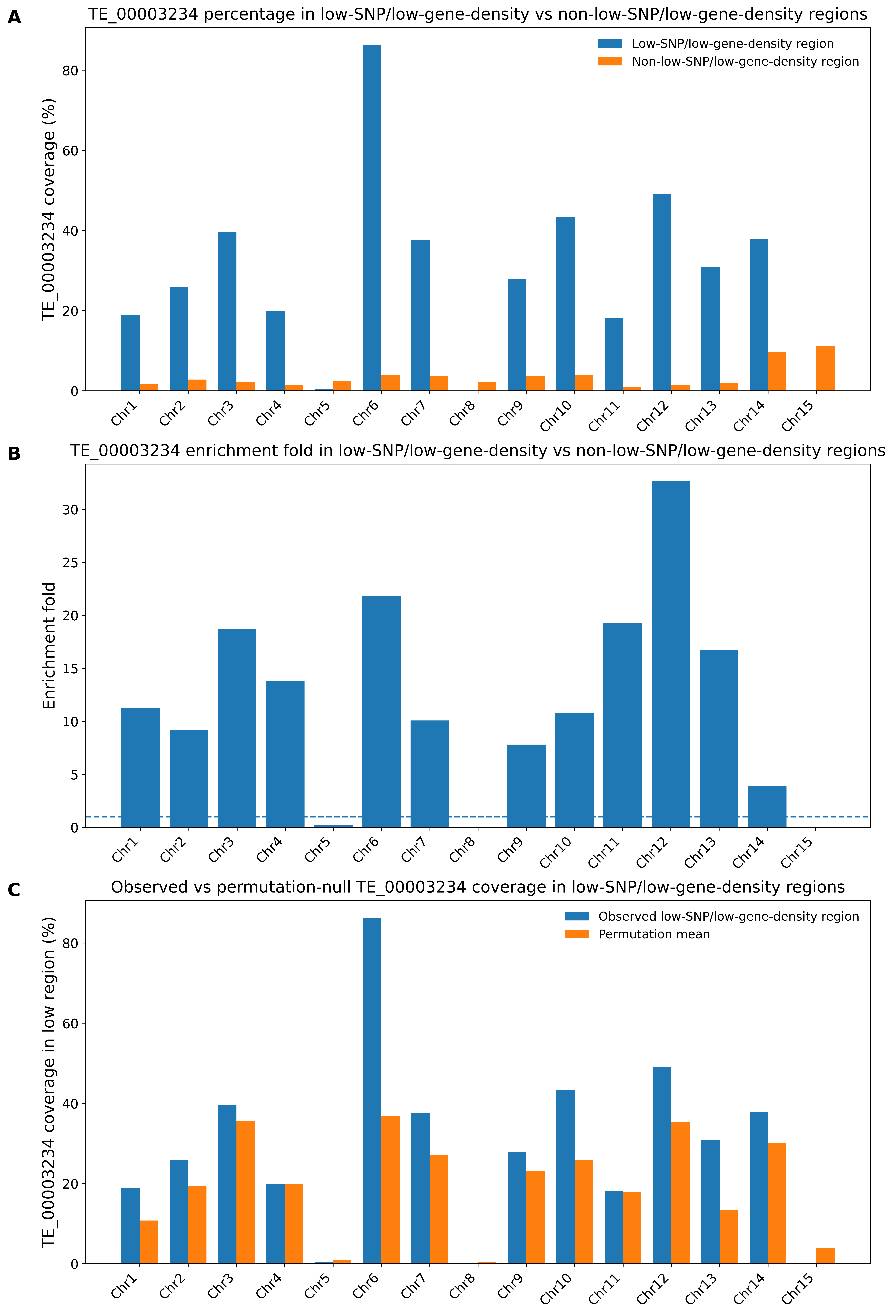

**Supplemental Figure S12. *TE_00003234* is strongly enriched in low-SNP/low-gene-density regions across flax chromosomes.**

(**A**) Percentage of genomic sequence occupied by *TE_00003234* in low-SNP/low-gene-density regions (blue) compared with non-low regions (orange) across the 15 chromosomes of *L. usitatissimum* (CDC Bethune v3.0). *TE_00003234* accounts for a substantial proportion of sequence in low-density regions, reaching up to ~85%, while remaining low in non-low regions.

(**B**) Enrichment fold of *TE_00003234* in low-SNP/low-gene-density regions relative to non-low regions. The dashed line indicates no enrichment (fold = 1). Most chromosomes show strong enrichment, ranging from ~8-fold to >30-fold.

(**C**) Observed *TE_00003234* coverage in low-SNP/low-gene-density regions (blue) compared with permutation-based expectations (orange). Permutation tests were performed by randomly redistributing low-region intervals within each chromosome while preserving interval size and number. Across all chromosomes, observed values exceed the permutation mean, indicating that the enrichment of *TE_00003234* is unlikely to arise by chance.

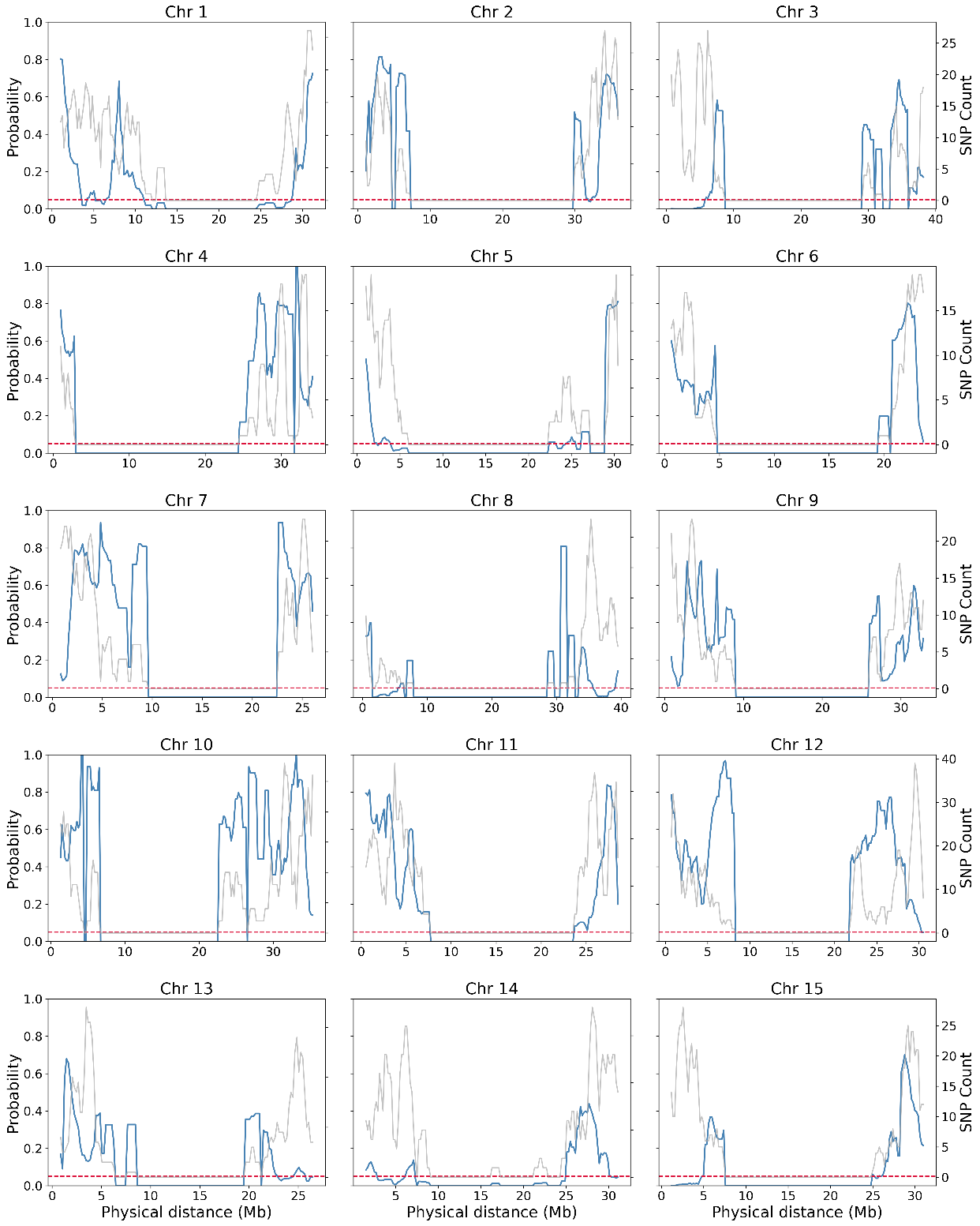

**Supplemental Figure S13.** **Segregation distortion probabilities (primary Y-axis, blue line) and SNP counts (secondary Y-axis, grey line) plotted along physical positions (in base pairs, bp; based on the reference genome CDC Bethune v3.0) for 158 Linda × Norman recombinant inbred lines (RILs).** A sliding window of 500 kb with a step size of 100 kb was used. The red dotted line indicates the 5% significance threshold for the Chi-square test of segregation distortion.

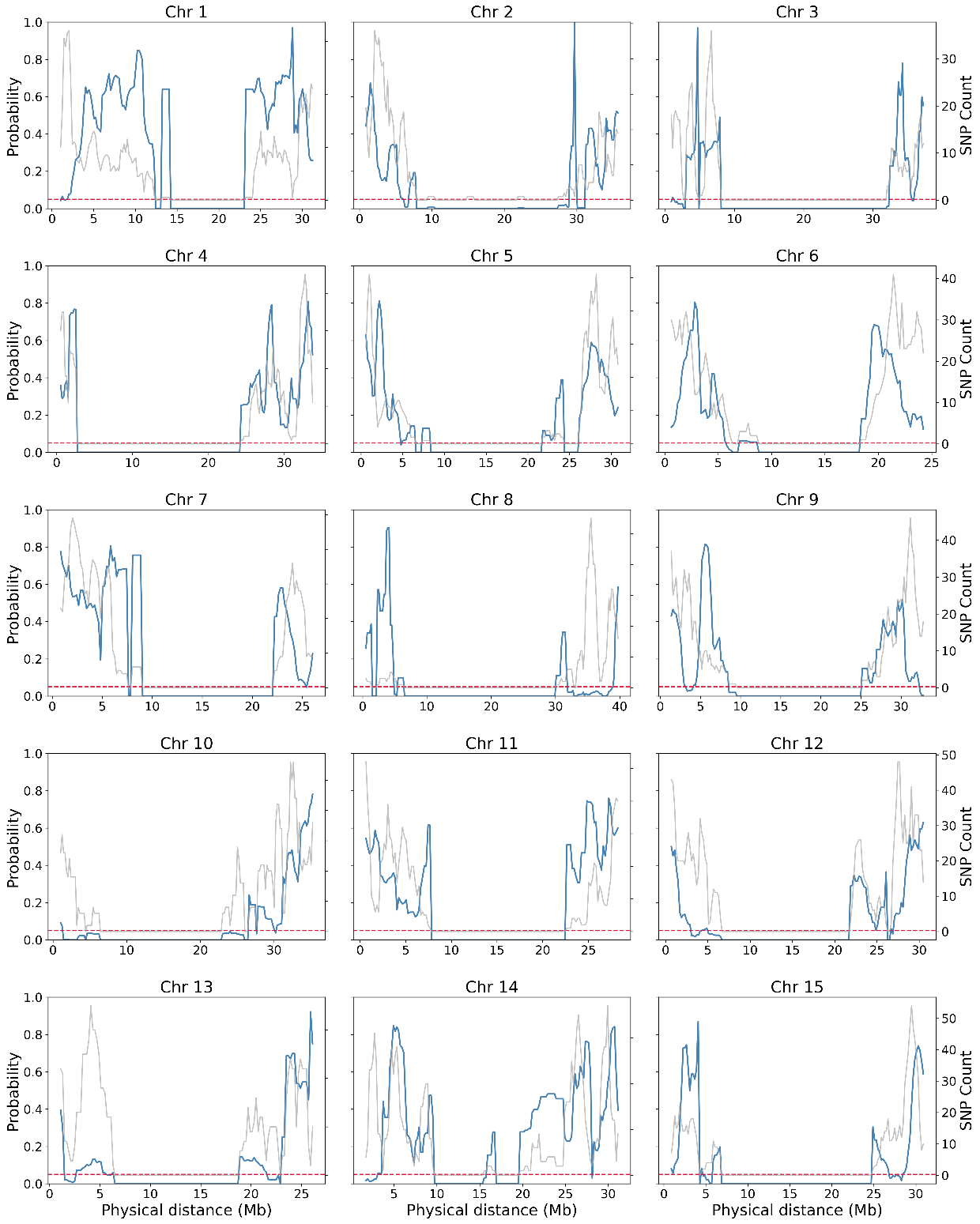

**Supplemental Figure S14.** **Segregation distortion probabilities (primary Y-axis, blue line) and SNP counts (secondary Y-axis, grey line) plotted along physical positions (in base pairs, bp; based on the reference genome CDC Bethune v3.0) for 702 Bison × Novelty recombinant inbred lines (RILs)**. A sliding window of 500 kb with a step size of 100 kb was used. The red dotted line indicates the 5% significance threshold for the Chi-square test of segregation distortion.

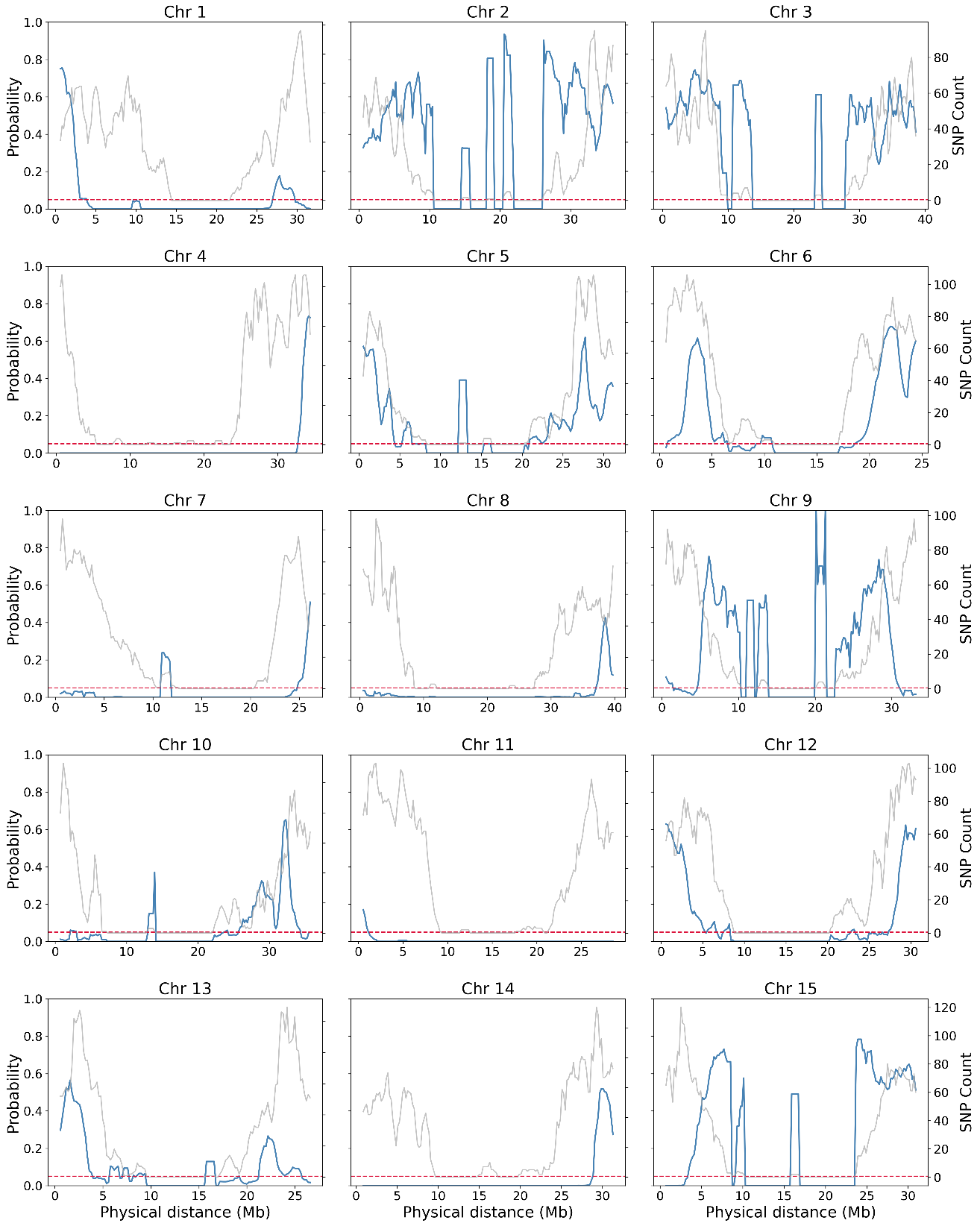

**Supplemental Figure S15.** **Segregation distortion probabilities (primary Y-axis, blue line) and SNP counts (secondary Y-axis, grey line) plotted along physical positions (in base pairs, bp; based on the reference genome CDC Bethune v3.0) for 166 LIN1917 × CDC Bethune recombinant inbred lines (RILs)**. A sliding window of 500 kb with a step size of 100 kb was used. The red dotted line indicates the 5% significance threshold for the Chi-square test of segregation distortion.

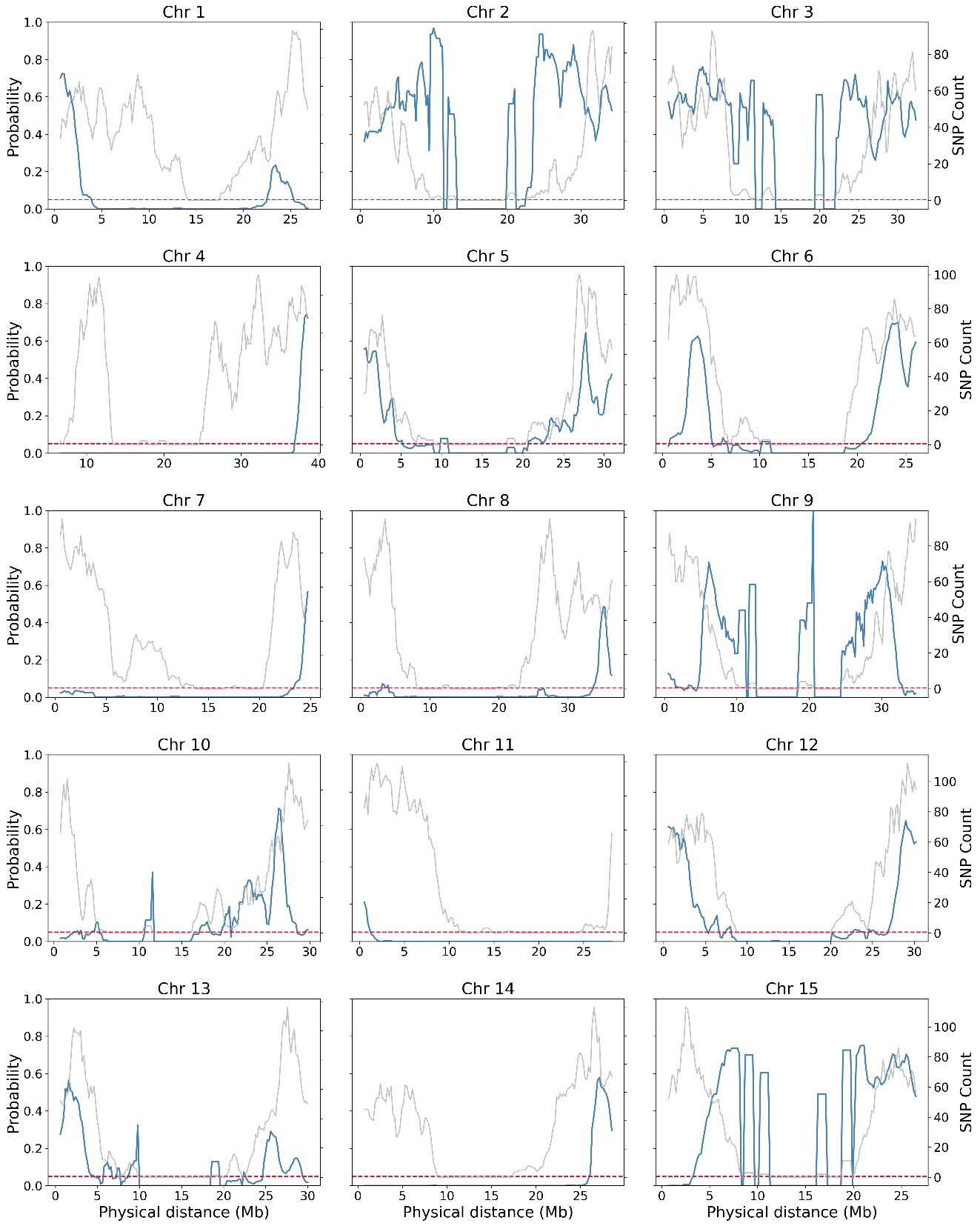

**Supplemental Figure S16.** **Segregation distortion probabilities (primary Y-axis, blue line) and SNP counts (secondary Y-axis, grey line) plotted along physical positions (in base pairs, bp; based on the reference genome LIN1917) for 166 LIN1917 × CDC Bethune recombinant inbred lines (RILs).** A sliding window of 500 kb with a step size of 100 kb was used. The red dotted line indicates the 5% significance threshold for the Chi-square test of segregation distortion.

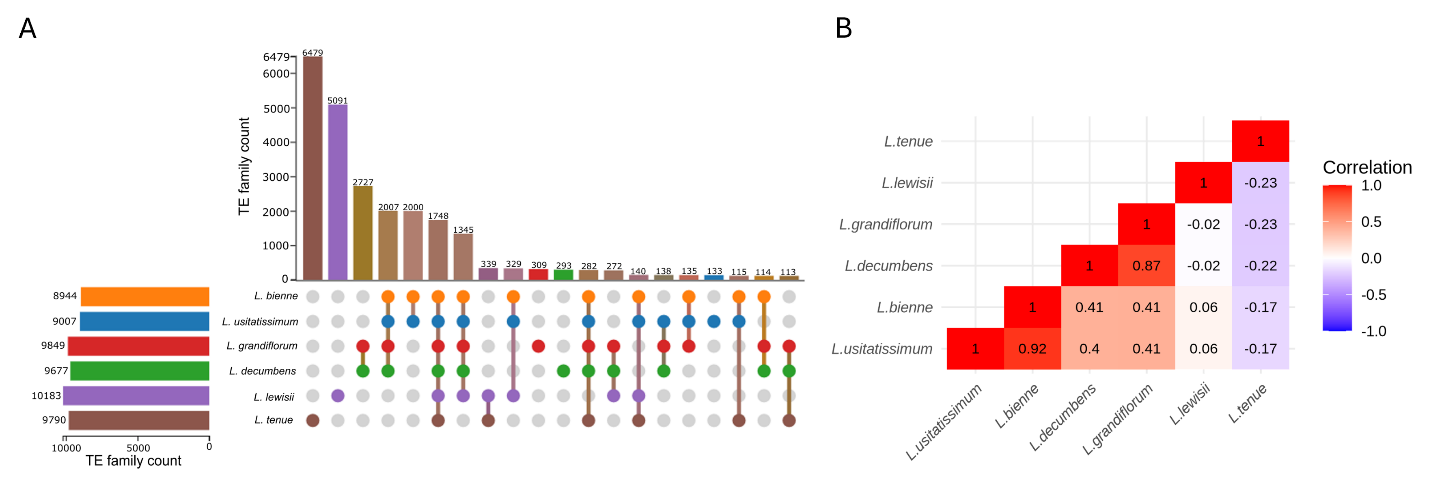

**Supplemental Figure S17. Comparison of the number of transposable element (TE) families among six *Linum* species.** (A) UpSet plot showing the total number of TE families in each species and the shared TE families across species. (B) Correlation matrix of TE families among the six species. Only TE families with sequences > 1 kb aligned to a genome were tallied.

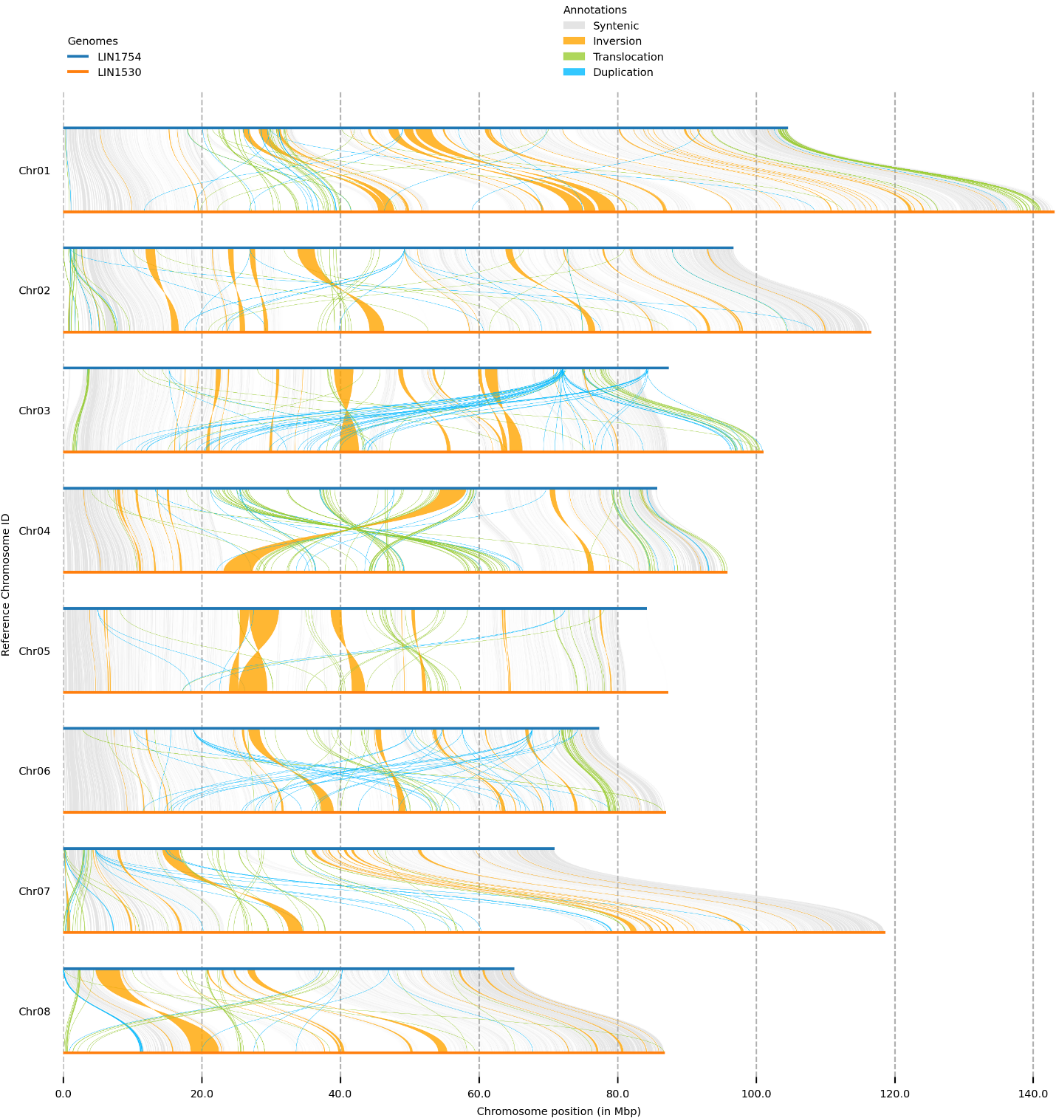

**Supplemental Figure S18.** Chromosome-by-chromosome comparisons between *L. decumbens* accession LIN1754 (upper chromosomes; blue) and *L. grandiflorum* accession LIN1530 (lower chromosomes; orange) illustrating syntenic regions (grey), inversions (yellow), translocations (green) and duplications (blue).

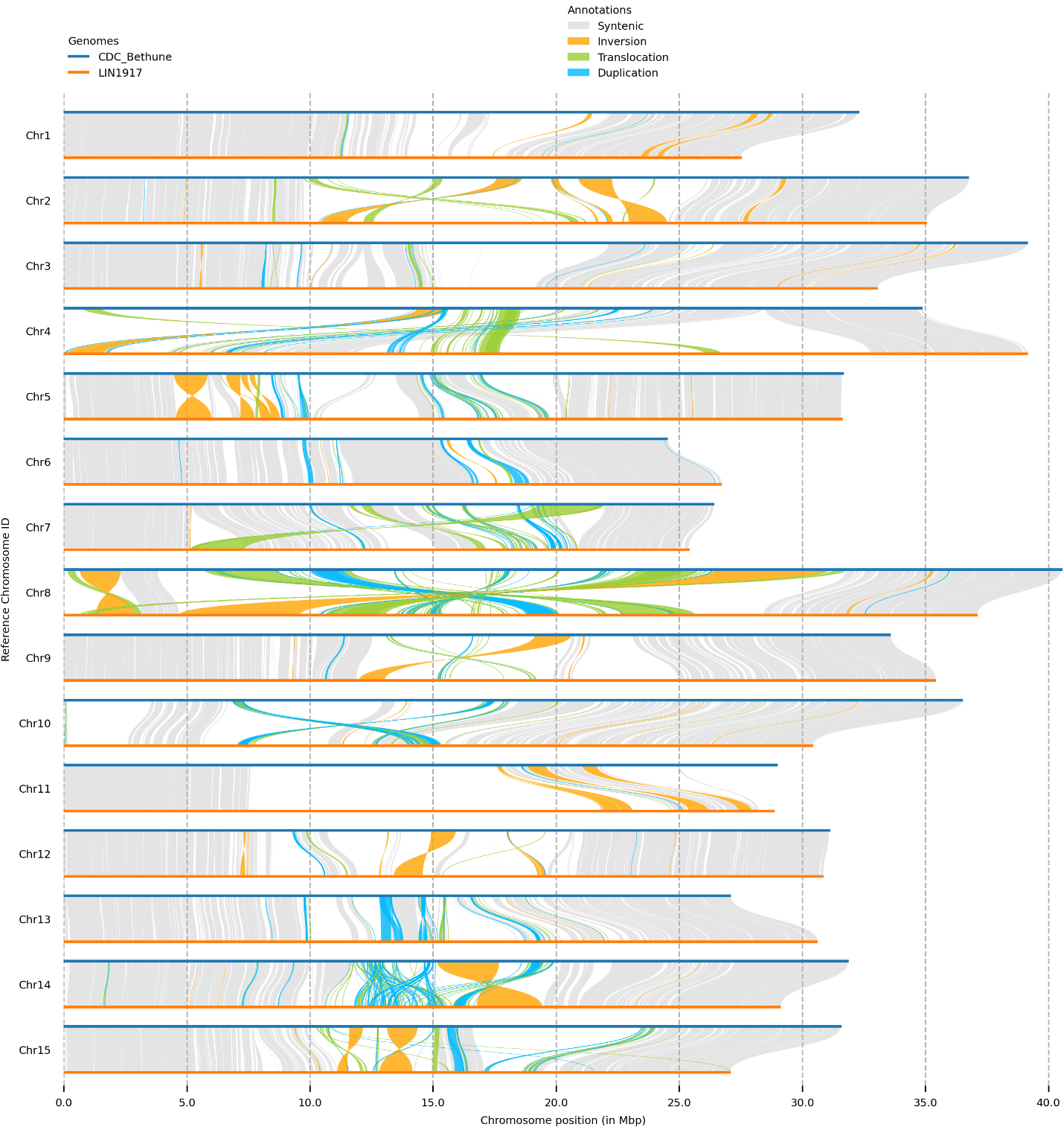

**Supplemental Figure S19.** Chromosome-by-chromosome comparisons between *L. usitatissimum* cultivar CDC Bethune (upper chromosomes; blue) and *L. bienne* accession LIN1917 (lower chromosomes; orange) illustrating syntenic regions (grey), inversions (yellow), translocations (green) and duplications (blue).

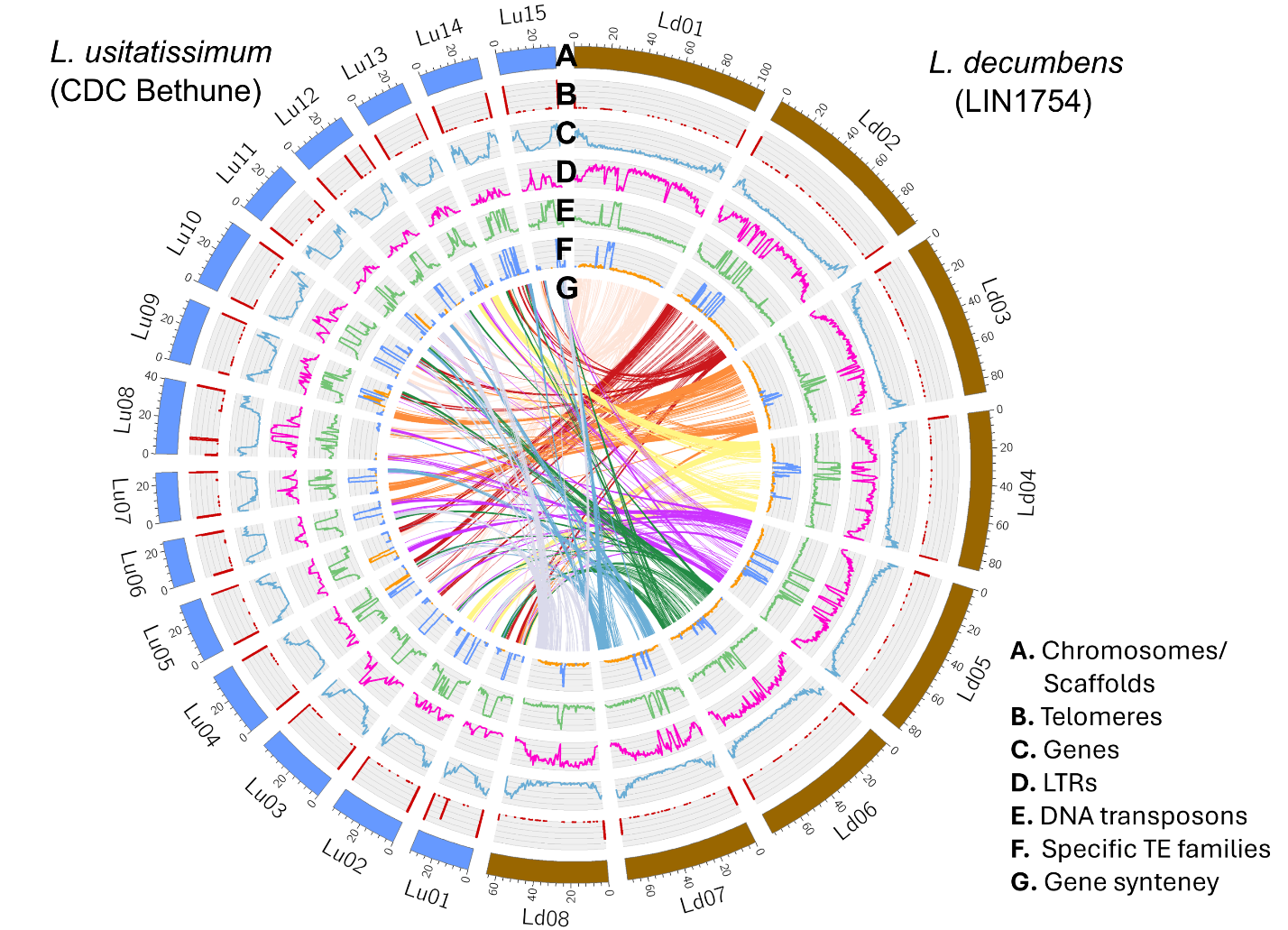

**Supplemental Figure S20. Circos map of the assemblies for *L. decumbens* accession LIN1754 and L. usitatissimum cultivar CDC Bethune.** The outermost track (A) represents the chromosomes. The second track (red) (B) shows the locations of telomeres, depicted by peaks of the simple repeat sequence ‘TTTAGGG’. The third track (light blue) (C) shows the gene density. The fourth track (pink) (D) represents the density of long terminal repeat (LTR) transposable elements (TEs). The fifth track (green) (D) represents the density of DNA transposons. The sixth track (F) shows the density of the predominant DNA transposon family *TE_00029583* (medium blue) for *L. decumbens*, and the density of the predominant DNA transposon family *TE_00003234* (medium blue) for *L. usitatissimum*. The central circle (G) illustrates gene collinearity (synteny) between the chromosomes of LIN1754 and CDC Bethune.

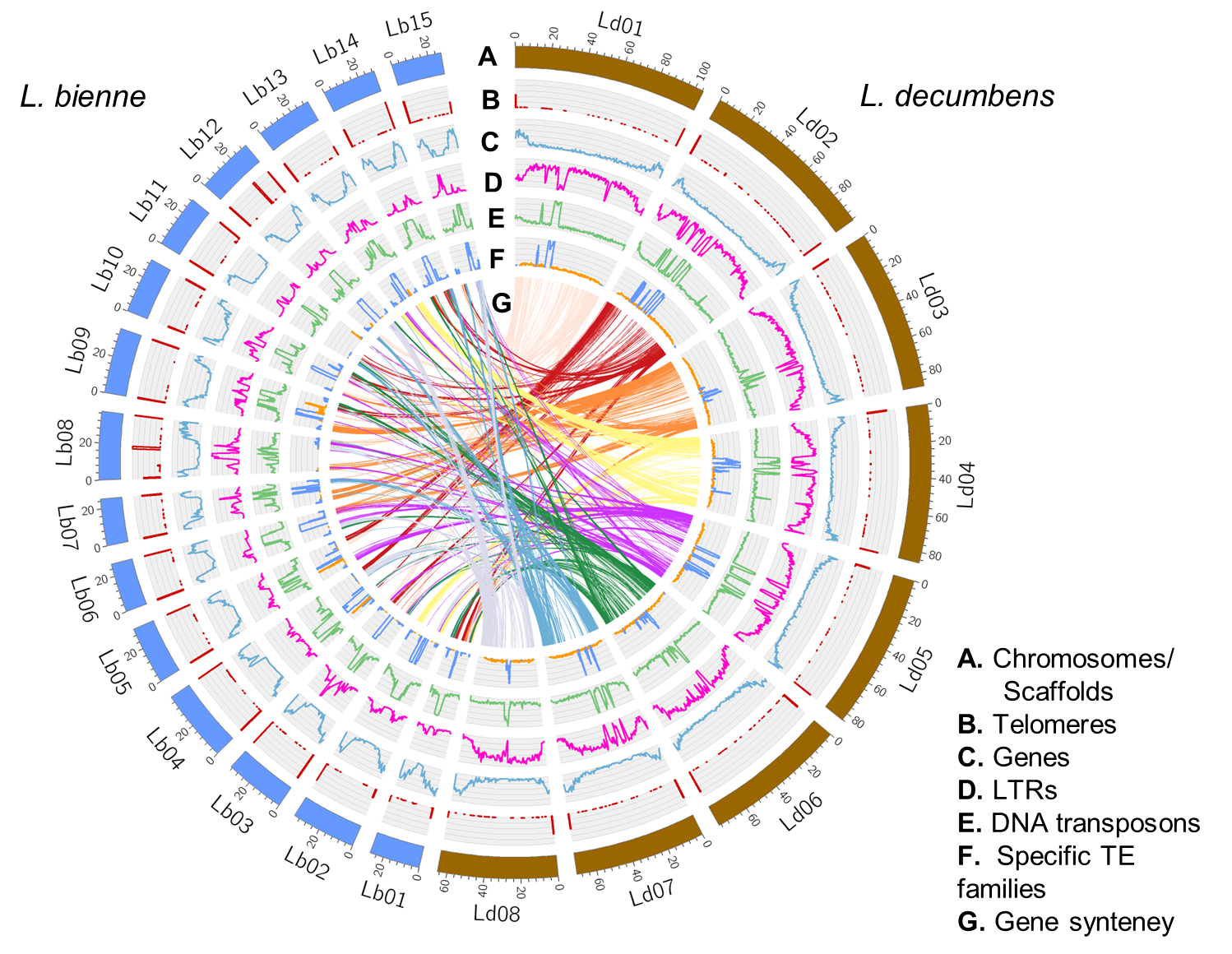

**Supplemental Figure S21. Circos map of the assemblies for *L. decumbens* accession LIN1754 and L. bienne accession LIN1917.** The outermost track (A) represents the chromosomes. The second track (red) (B) shows the locations of telomeres, depicted by peaks of the simple repeat sequence ‘TTTAGGG’. The third track (light blue) (C) shows the gene density. The fourth track (pink) (D) represents the density of long terminal repeat (LTR) transposable elements (TEs), and the fifth track (green) (E) represents the density of DNA transposons. The sixth track (F) shows the density of the predominant DNA transposon family *TE_00029583* (medium blue for *L. decumbens*, and the density of the predominant DNA transposon family *TE_00003234* (medium blue) for *L. bienne*. The central circle (G) illustrates the gene collinearity (synteny) between the chromosomes of LIN1754 and LIN1917.

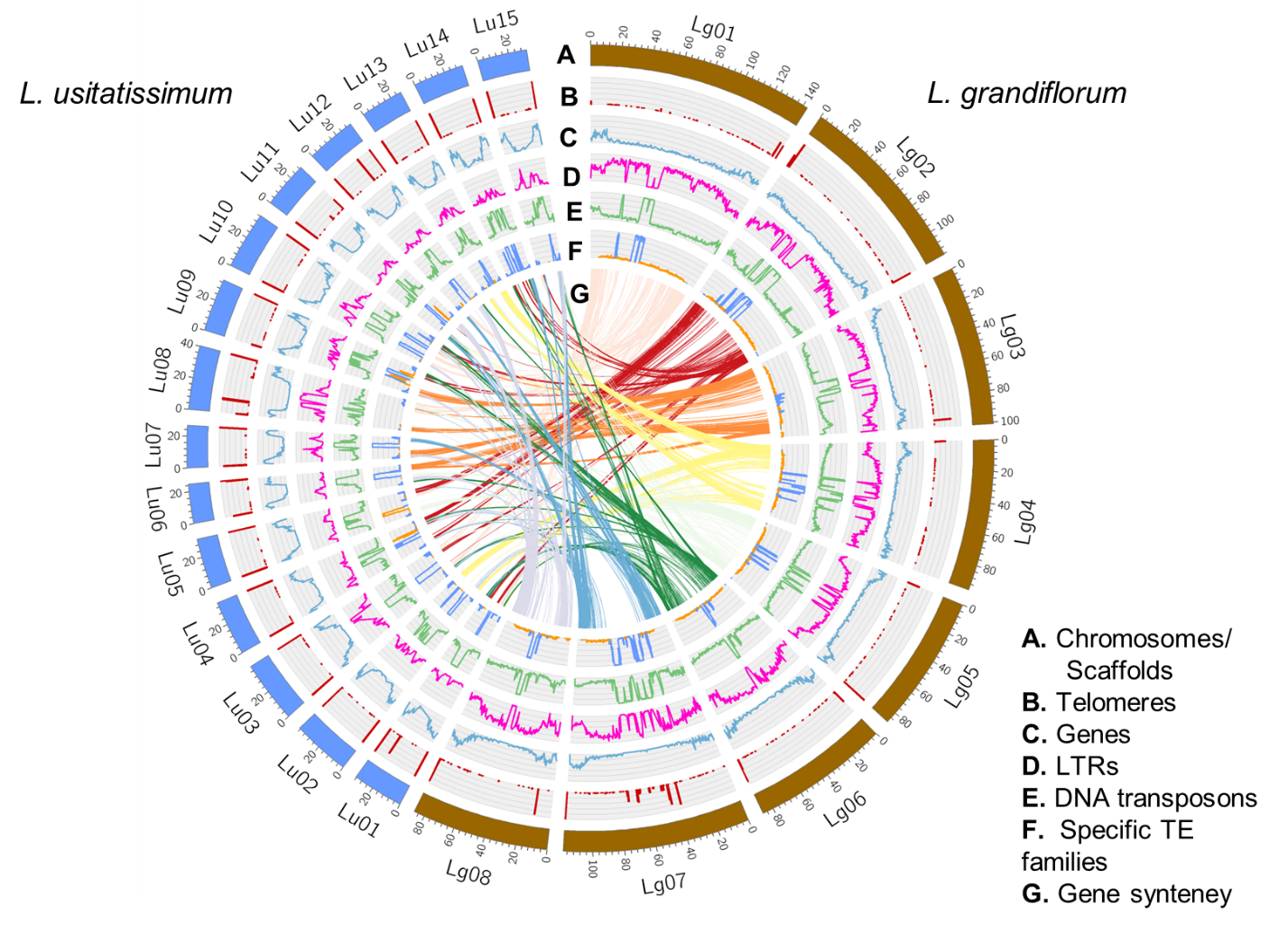

**Supplemental Figure S22. Circos map of the assemblies for *L. grandiflorum* accession LIN1530 and** L. usitatissimum **cultivar CDC Bethune.** The outermost track (A) represents the chromosomes. The second track (red) (B) shows the locations of telomeres, depicted by peaks of the simple repeat sequence ‘TTTAGGG’. The third track (light blue) (C) shows the gene density. The fourth track (pink) (D) represents the density of long terminal repeat (LTR) transposable elements (TEs). The fifth track (green) (D) represents the density of DNA transposons. The sixth track (F) shows the density of the predominant DNA transposon family *TE_00029583* (medium blue) for *L. grandiflorum*, and the density of the predominant DNA transposon family *TE_00003234* (medium blue) for *L. usitatissimum*. The central circle (G) illustrates gene collinearity (synteny) between the chromosomes of LIN1530 and CDC Bethune.

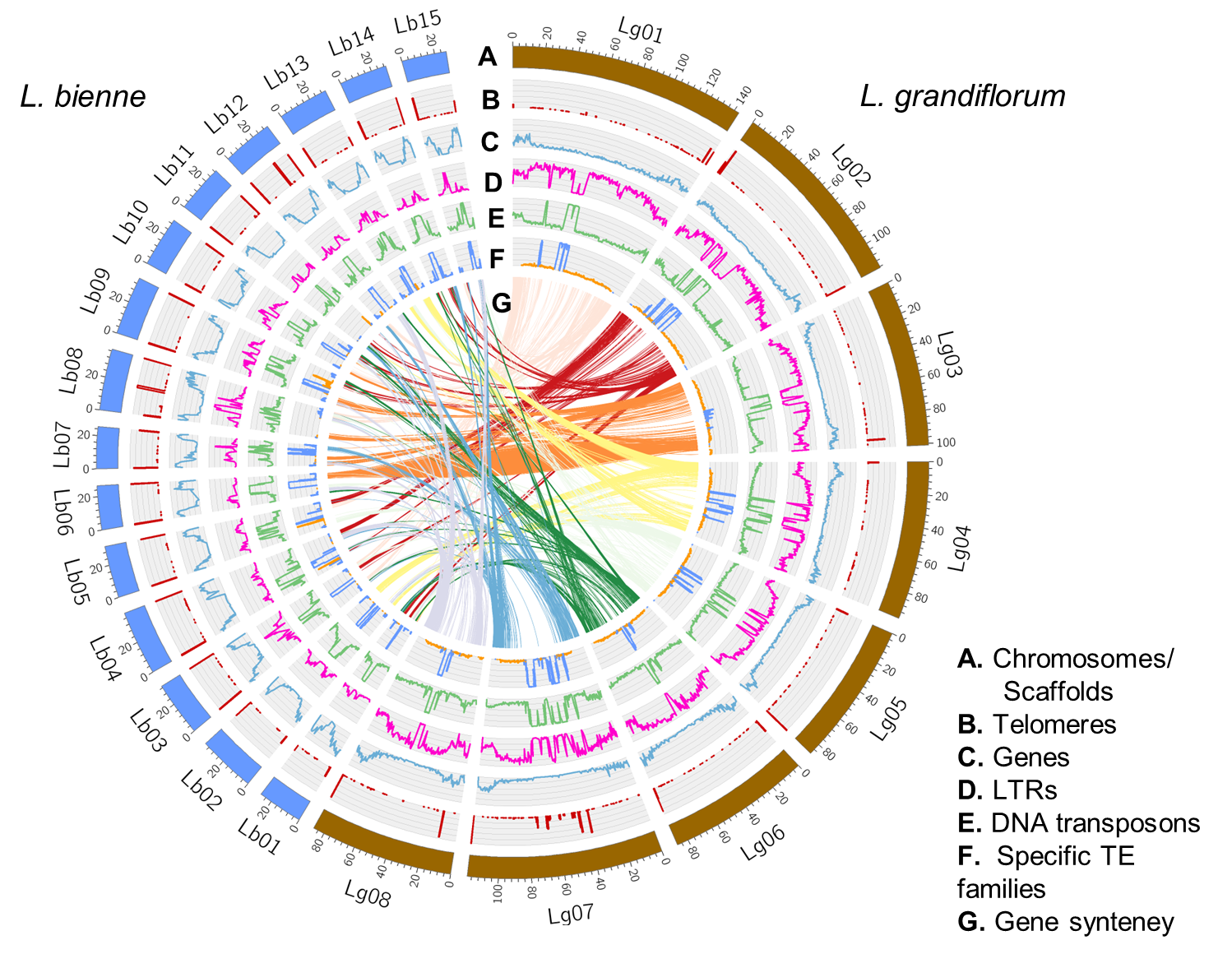

**Supplemental Figure S23. Circos map of the assemblies for *L. grandiflorum* accession LIN1530 and L. bienne accession LIN1917.** The outermost track (A) represents the chromosomes. The second track (red) (B) shows the locations of telomeres, depicted by peaks of the simple repeat sequence ‘TTTAGGG’. The third track (light blue) (C) shows the gene density. The fourth track (pink) (D) represents the density of long terminal repeat (LTR) transposable elements (TEs). The fifth track (green) (D) represents the density of DNA transposons. The sixth track (F) shows the density of the predominant DNA transposon family *TE_00029583* (medium blue for *L. grandiflorum*, and the density of the predominant DNA transposon family *TE_00003234* (medium blue) for *L. bienne*. The central circle (G) illustrates gene collinearity (synteny) between the chromosomes of LIN1530 and LIN1917.

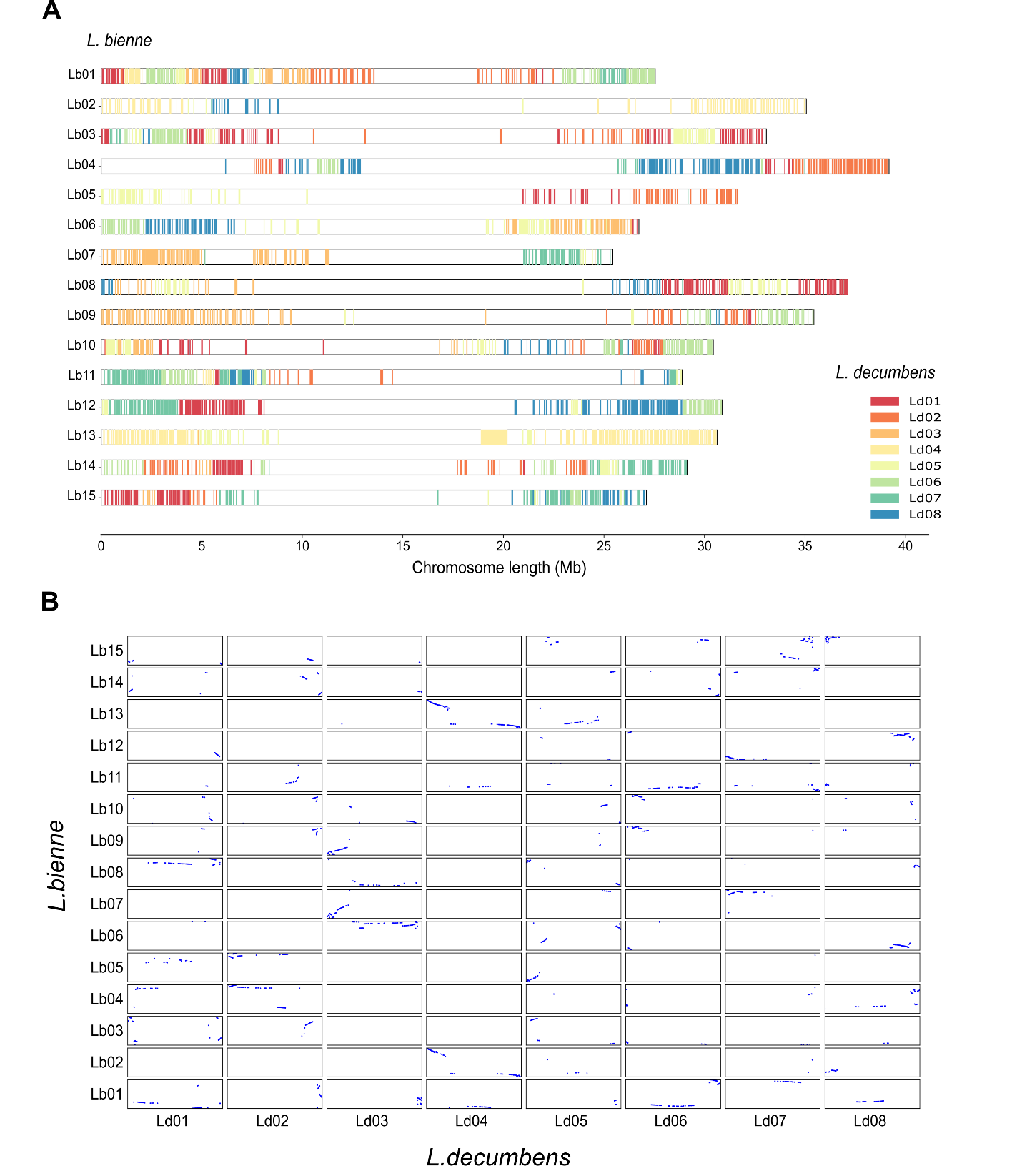

**Supplemental Figure S24. Distribution and collinearity of orthologous genes between *L. binne* and *L. decumbens*.** (**A**) chromosomal distribution of *L. decumbens* orthologs along the chromosomes of *L. binne*. Each horizontal line represents one *L. binne* chromosome, and colored ticks indicate the positions of orthologous genes, with colors corresponding to their chromosome of origin in *L. decumbens* (Ld01–Ld08). Multiple labels at chromosome termini denote the major contributing *L. decumbens* chromosomes. (**B**) Dot plot showing pairwise collinearity between *L. binne* (y-axis) and *L. decumbens* (x-axis) chromosomes based on orthologous gene pairs. Blue dots represent collinear gene matches.

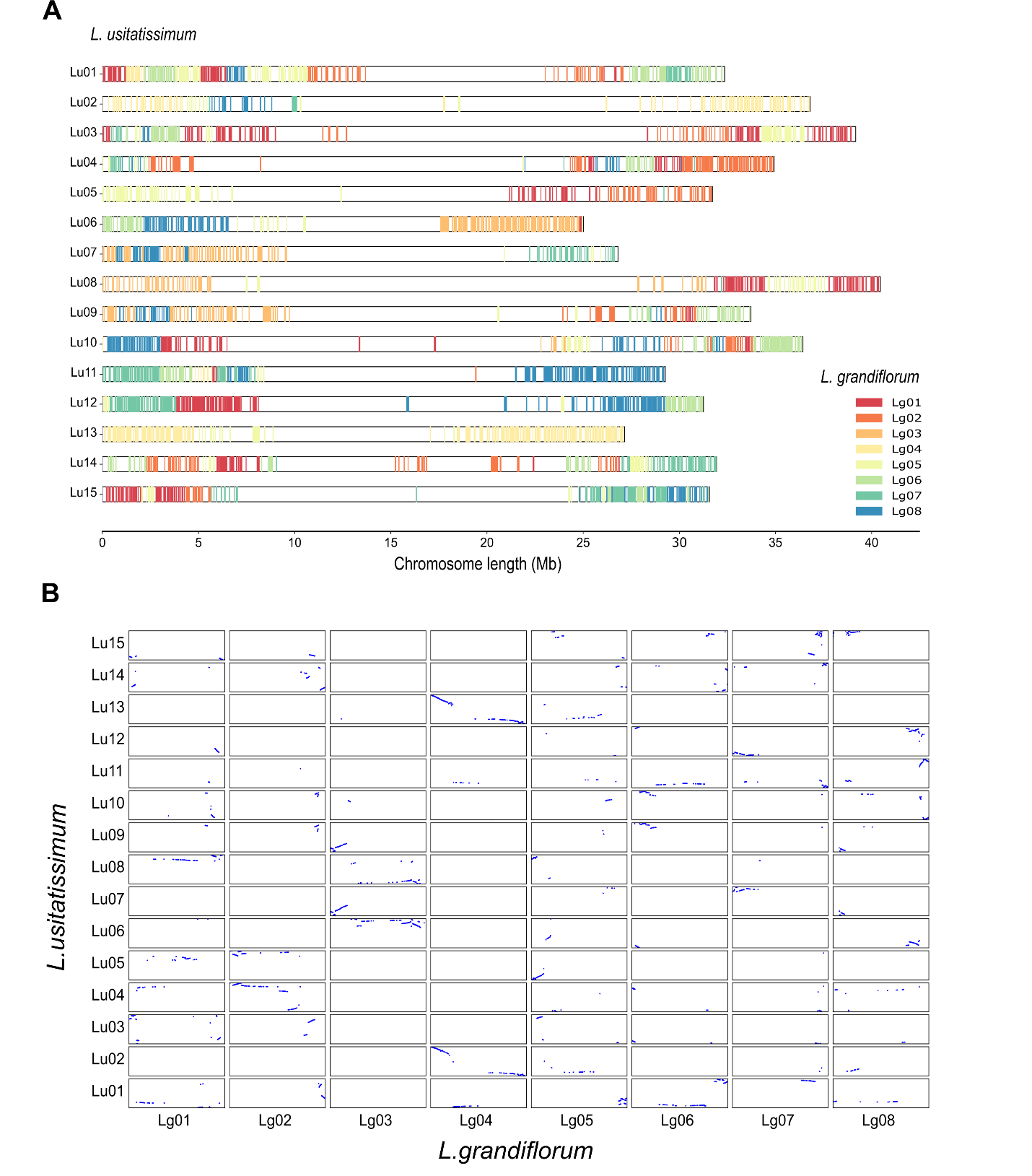

**Supplemental Figure S25. Distribution and collinearity of orthologous genes between *L. usitatissimum* and *L. grandiflorum*.** (**A**) Chromosomal distribution of *L. grandiflorum* orthologs along the chromosomes of *L. binne*. Each horizontal line represents one *L. usitatissimum* chromosome, and colored ticks indicate the positions of orthologous genes, with colors corresponding to their chromosome of origin in *L. grandiflorum* (Lg01–Lg08). Multiple labels at chromosome termini denote the major contributing *L. grandiflorum* chromosomes. (**B**) Dot plot showing pairwise collinearity between *L. usitatissimum* (y-axis) and *L. grandiflorum* (x-axis) chromosomes based on orthologous gene pairs. Blue dots represent collinear gene matches.

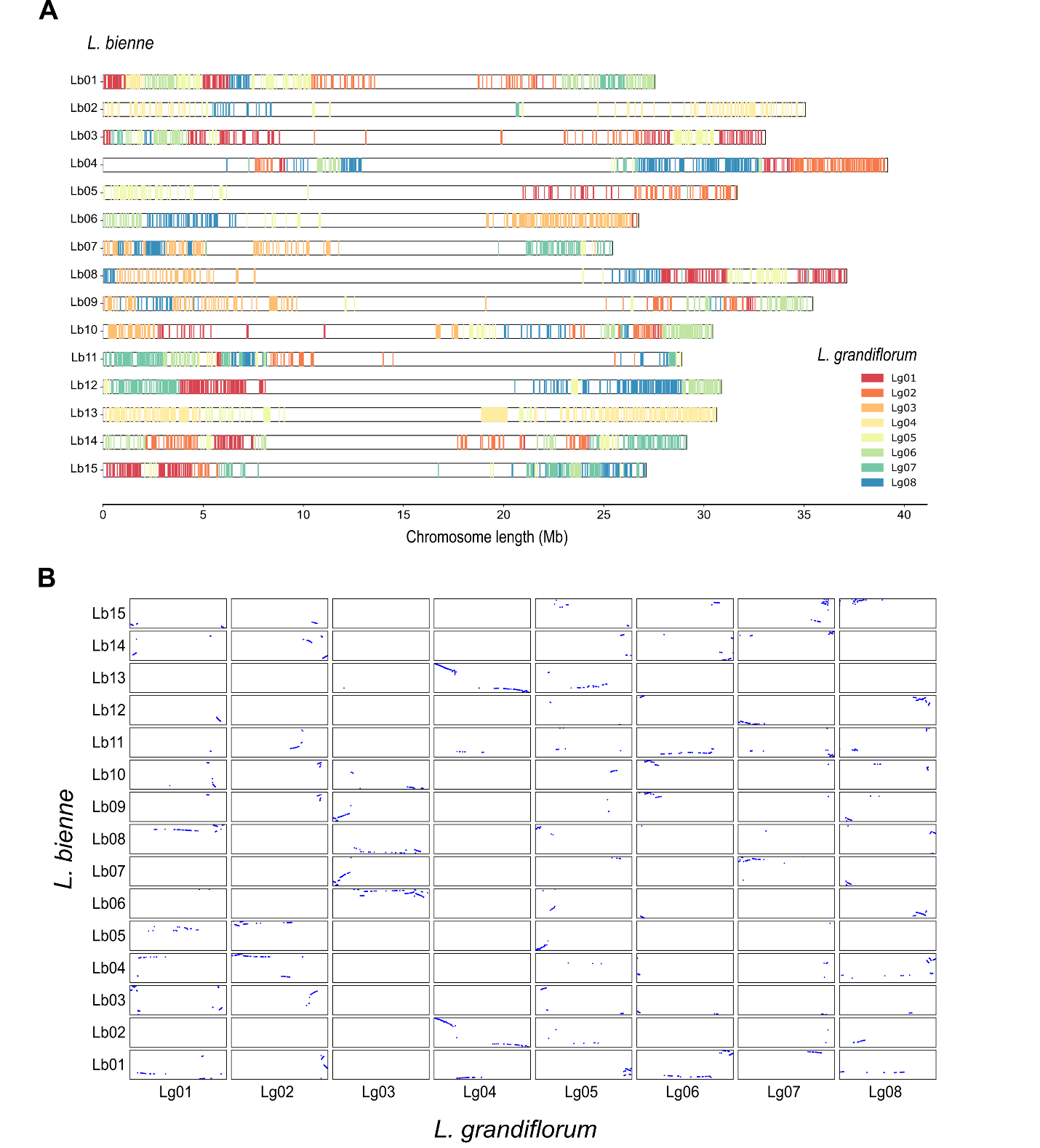

**Supplemental Figure S26. Distribution and collinearity of orthologous genes between *L. bienne* and *L. grandiflorum*.** (**A**) Chromosomal distribution of *L. grandiflorum* orthologs along the chromosomes of *L. binne*. Each horizontal line represents one *L. bienne* chromosome, and colored ticks indicate the positions of orthologous genes, with colors corresponding to their chromosome of origin in *L. grandiflorum* (Lg01–Lg08). Multiple labels at chromosome termini denote the major contributing *L. grandiflorum* chromosomes. (**B**) Dot plot showing pairwise collinearity between *L. bienne* (y-axis) and *L. grandiflorum* (x-axis) chromosomes based on orthologous gene pairs. Blue dots represent collinear gene matches.

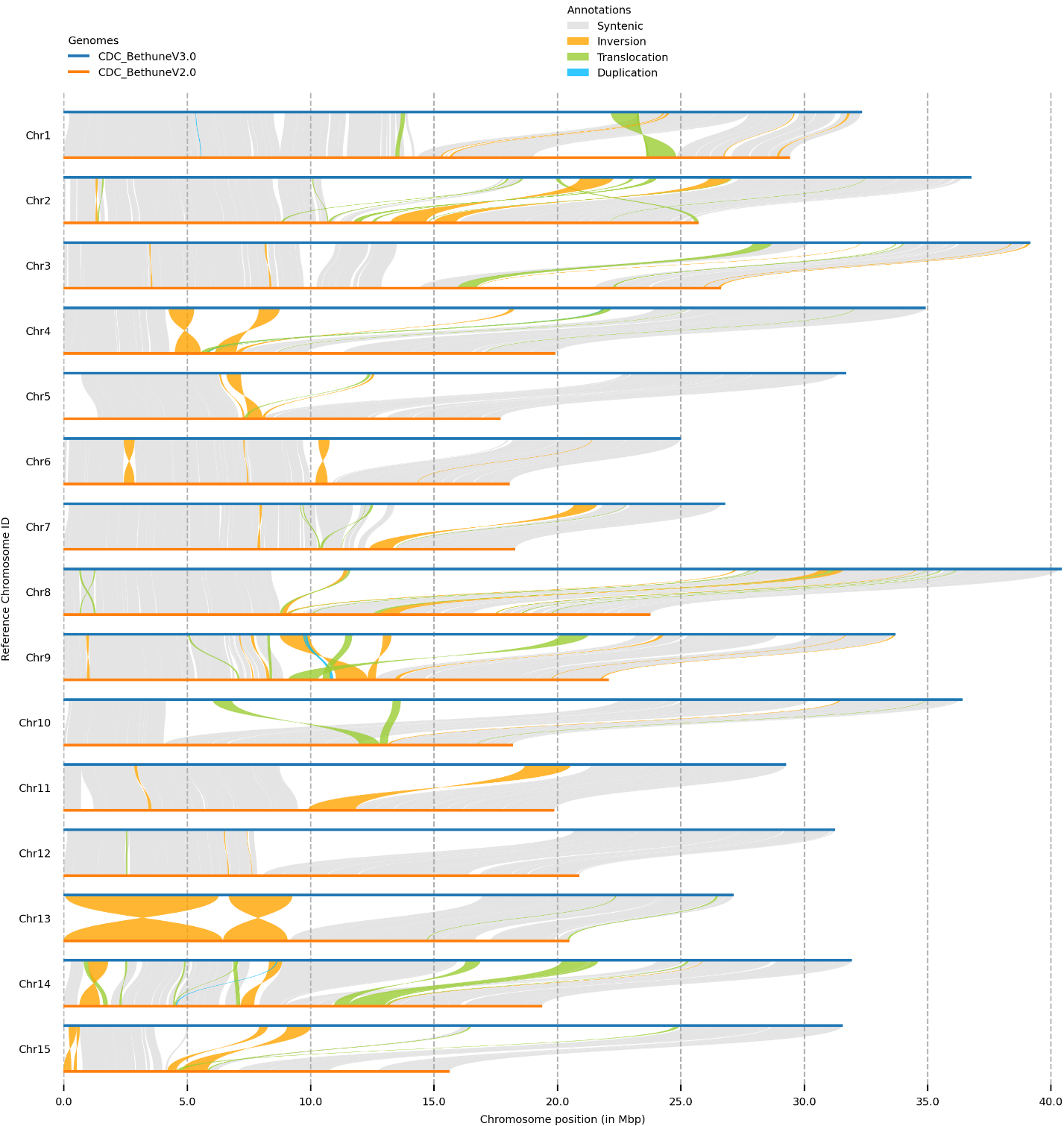

**Supplemental Figure S27.** Chromosome-by-chromosome comparisons between *L. usitatissimum* cultivar CDC Bethune v3.0 (upper chromosomes; blue) and v2.0 (lower chromosomes; orange) illustrating structural improvement and corrections.

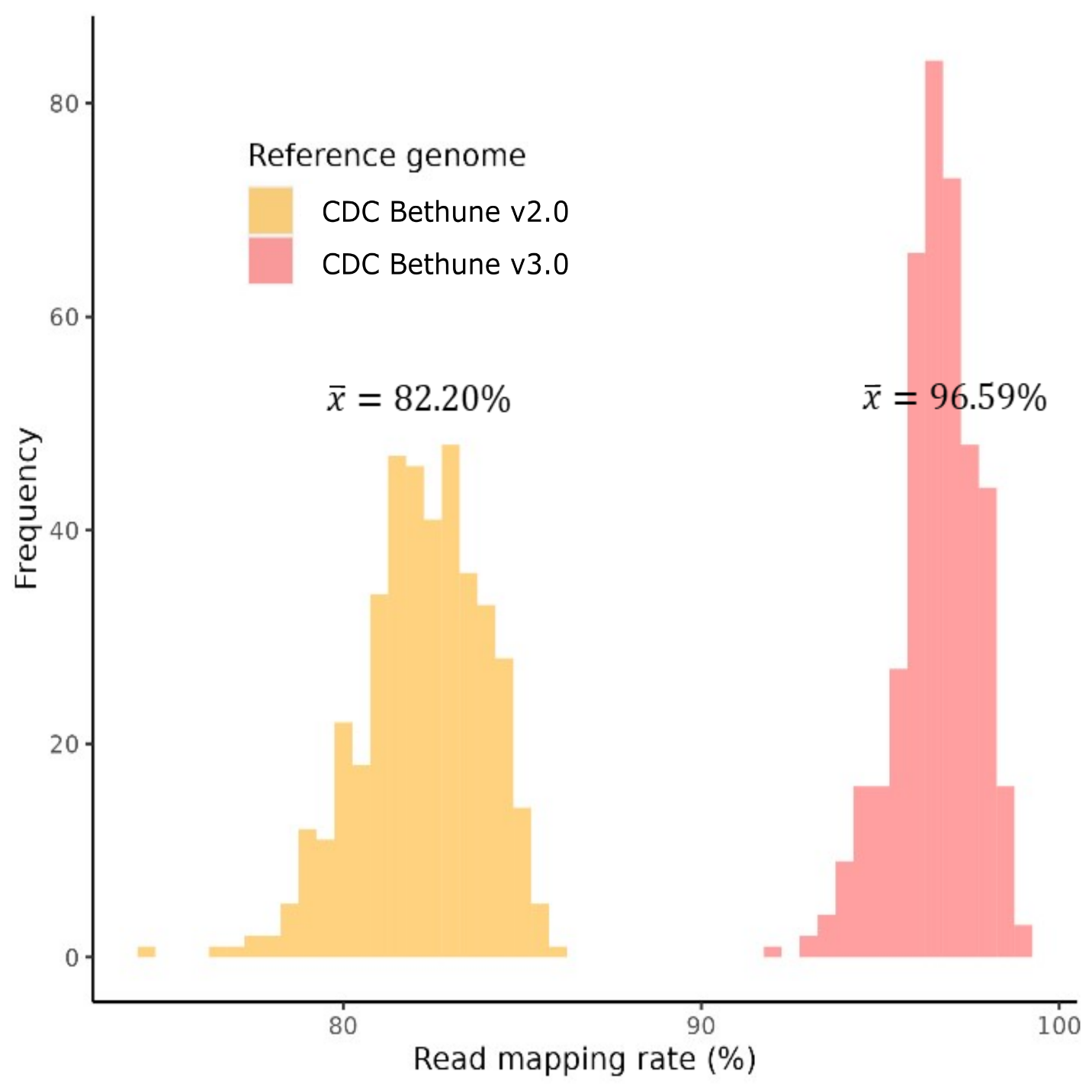

**Supplemental Figure S28. Improved read mapping rates to the telomere-to-telomere CDC Bethune v3.0 flax genome assembly.** Distribution of short-read mapping rates for 407 flax core accessions (You et al. 2022) aligned to two versions of the flax reference genome: CDC Bethune v2.0 (You et al. 2018) and the telomere-to-telomere CDC Bethune v3.0 assembly generated in this study. Mapping to v3.0 resulted in consistently higher read mapping rates (mean = 96.59%) compared with v2.0 (mean = 82.20%), indicating substantially improved genome completeness and sequence continuity in the v3.0 assembly. Histograms show the frequency distribution of mapping rates across accessions, highlighting reduced variability and fewer unmapped reads when using the v3.0 reference.
